# The Regional Vulnerability Index (RVI) as a Neuroimaging-Based Biomarker for Autism: Associations with Likelihood, Cognition, and Longitudinal Social Outcomes

**DOI:** 10.64898/2026.05.19.726341

**Authors:** Antonio Pagán, Katherine E. Lawrence, Jan Buitelaar, Si Gao, Paul M. Thompson, Yizhou Ma, Kelly T. Cosgrove, Fernanda Laezza, D.M. Hafeman, Brian Donohue, Bhim M. Adhikari, Anilkumar Pillai, Neda Jahanshad, Wen Li, Sophia Thomopoulos, Katherine A. Loveland, Ron Acierno, Alia Warner, Cecilia Montiel-Nava, Carly Demopoulos, Jeffrey R. Temple, Jair C. Soares, Shuo Chen, L. Elliot Hong, The ENIGMA autism Working Group, Peter Kochunov

## Abstract

Autism spectrum disorder (ASD) is a complex neurodevelopmental condition with symptoms spanning cognitive, social and behavioral domains and leading to lifelong challenges. Autism is heritable but specific genetic and environmental factors that shape its early neurodevelopmental trajectory remain unknown. Despite the clinical variability, neuroimaging findings from Enhancing Neuro Imaging Genetics through Meta Analysis (ENIGMA)-ASD consortium identified stable and replicable patterns of cortical and subcortical differences that were consistent with those reported by an independent consortium, the Cognitive Genetics Collaborative Research Organization (COCORO) in Japan. Here we developed and evaluated a regional vulnerability index (RVI), a similarity metric that quantifies how closely a person’s brain resembles the specific pattern of an individual with autism. RVI-ASD is based on combining the regional effect sizes for regional brain measurements published by the ENIGMA-ASD group with microstructural white matter integrity differences reported by COCORO. RVI-ASD showed significantly higher effect size for case-control differences relative to any individual brain measure (d=0.30 vs. d=0.01–0.21) in individuals with autism, particularly in the adolescent-to-adult sample (N=2,577; Mean age = 16.1; SD=5.0). We next calculated RVI-ASD in the baseline and follow-up (ages 10 and 12) data from normally developing participants of the ABCD study (N=4,201). We tested the longitudinal stability, heritability, genotype-by-age interaction and sensitivity of RVI-ASD to known factors and cognitive measurements. RVI-ASD were stable on the 2-year follow up (ICC=0.76–0.92); showed significant heritability (h²=0.55-0.83, p<10^-16^) but was not affected by gene-by-age interaction. RVI-ASD showed significant positive correlation with paternal age, while correlation with maternal age was non-significant. Baseline and follow-up RVI-ASD were negatively correlated with cognitive measures including total, fluid and crystallized intelligence (r=-0.09 to -0.11, p<10^-6^). RVI-ASD scores tracked with core autism phenotypes including poor eye contact and rigid routines (p < .01). In a sub-sample of children with symptoms of autism (N=20), we found that earlier age of symptom onset was strongly correlated with higher White Matter RVI (r = -0.61), linking early behavioral emergence to the neuroanatomical signature. Longitudinal changes in subcortical RVI-ASD are significantly correlated with change in social functioning. The RVI-ASD is a neuroimaging-based biomarker linked to stable and reproducible brain patterns in autism. RVI-ASD allows researchers to study associations with factors associated with the likelihood for autism and cognition across large and inclusive non-clinical samples, thus moving beyond simple case-control models to understand the biological pathways of autism.

## Introduction

Autism spectrum disorder (ASD) is a complex neurodevelopmental condition characterized by challenges in cognitive, social, communication and behavioral domains [1]. Autism prevalence rates continue to rise, being diagnosed in ∼1 in 30 children [2] and there is a critical need to understand the neurobiological underpinnings to support the well-being of individuals with autism and their families [3]. Small samples sizes in studies of autism and high clinical variability has led to variable neuroimaging findings [4–6]. Here, we adopted a ‘big data’ neuroimaging approach to demonstrate that individuals with autism share a stable and reproducible pattern of neuroanatomical differences. We used this pattern to pilot the first regional vulnerability index in autism (RVI-ASD), a quantitative metric that measures how closely an individual’s brain morphology resembles the characteristic neuroanatomical “signature” of autism. Next, we applied this metric to a large cohort of normally developing adolescents to show that this signature scales dimensionally with subthreshold autistic traits, identifies known risks of autism and is predictive of cognitive performance.

Autism traits are identifiable across general population^1^ [7, 8] [9, 10]. We aim to validate a stable neural signature for autism to pivot to a dimensional, Research Domain Criteria (RDoC) approach [11]. We posit that the risks and trait that contribute to autism diagnosis are quantifiable in the general population, with a clinical diagnosis representing the most severe tail of this distribution [4–6]. This approach is supported by decades of research on the “Broader Autism Phenotype” (BAP), which quantified the severity of autism traits in relatives of individuals with autism [12, 13]. This dimensional approach offers greater statistical power to detect and quantify the association of factors associated with the likelihood of autism using large and inclusive population samples [14]. We show that RVI-ASD allows to investigate how genetic and environmental factors contribute to the expression of the brain signature of autism in the general population and inform about sources of heterogeneity [4–6]. Unlike categorical diagnoses, quantitative measures such as RVI can capture the graded nature of ASD-related biology, moving beyond the limitations of clinically defined cut-offs [4, 15].

We, and others, recently proposed to use the regional differences (pattern similarity metrics) as the basis for a new class of biomarkers that measure individual similarity to the expected autism patterns to study the continuum of autism traits across the population [16]. Biomarkers, such as RVI, measure agreement with the expected brain autism pattern rather than focusing on a single region or a circuit by aggregating information across multiple metrics and brain systems. The resulting composite measure is hypothesized to provide a more sensitive and specific biomarker compared to specific measures from any one individual brain region.

The RVI is a simple linear method that gauges the similarity of an individual’s brain measurements to the established differences reported in our foundational ENIGMA and mega-analytic studies [17–19]. By calculating a single, continuous score for each participant, the RVI-ASD treats the autism neuro-signature as a dimensional trait, aligning with the clinical shift in dimensional construct of autism [14].

The regional effect sizes used to calculate RVI-ASD were derived from the meta-analyses performed by the autism working group of the ENIGMA consortium of the cortical gray matter thickness and subcortical volumes (N=3,671) in individuals with autism compared to neurotypical controls [19–21]. We replicated our ENIGMA-ASD patterns (RVI) in an independent international cohort of autistic individuals, from the COCORO consortium in Japan [22, 23]. The COCORO consortium also reported effect sizes for the regional white matter differences [18]. We combined ENIGMA cortical and subcortical effects with COCORO white matter effect sizes combining to construct “whole-brain” RVI-ASD.

Autism runs in families with heritability of the disorder reported to be ∼70% (h^2^=0.6-8) with genetic predisposition playing a foundational role [24, 25]. Having a first degree relative with autism increases individual likelihood by up to ten times [26]. Autism is also influenced by other factors [27]. For example, higher paternal age (>35 years) is robustly associated with autism across studies [28, 29]. This is primarily linked to a multidimensional etiology, including a combination of de novo mutations and epigenetic mechanisms [30–33]. Finally, there is strong evidence for environment by genetic interactions, especially during prenatal and perinatal periods [34]. For example, immune-mediated conditions during pregnancy and birth can elevate the likelihood for autism [34]. The complex nature of these varied factors underscores the need to apply our validated, dimensional biomarker to a large, longitudinal population sample to test for etiological convergence outside of a clinical setting.

We employed a novel analytic approach to evaluate the utility of the RVI-ASD as a neuroimaging-based biomarker in both case-control and populational samples. Our objectives were two-fold, structured into two complementary studies. In Study I, we sought to confirm the stability of this neuroanatomical signature across diverse international clinical cohorts. We refined the autism brain signature using an updated ENIGMA-ASD dataset and evaluated its reproducibility against the independent COCORO consortium. In Study II, we applied the refined RVI-ASD signature to a sample of youth from the ABCD study to investigate its expression in the general population. By applying the RVI-ASD to the longitudinal ABCD cohort, we aimed to disentangle biological liability for autism from clinical confounds. Specifically, we evaluated: A) the effects of genetic factors, gene-by-environment interactions, and developmental history on baseline RVI; B) longitudinal changes in the RVI signature during adolescence; and C) the relationship between this neural signature and cognitive development.

## Methods

### Overview

To characterize RVI-ASD as a neuroimaging-based biomarker for autism, we conducted two distinct but complementary studies. Study I focused on establishing the robustness and stability of the autism neuroanatomical signature using large-scale clinical datasets from the ENIGMA-ASD and COCORO working groups. Study II applied this refined signature to a large, longitudinal population-based sample of youth from the ABCD Study to investigate its heritability, genetic architecture, and relationship to dimensional traits in the general population.

### Study I: Defining and Confirming the autism Brain Signature

#### 2.1 Participants (ENIGMA-ASD)

Participants were drawn from the ENIGMA-ASD Working Group, a large-scale, multi-site effort, aggregating data from distinct international sites. See Supplementary Table 1 for full information regarding image acquisition parameters for each cohort. The final analytic dataset included 3,389 individuals drawn from 51 independent sites. The sample included 1,655 individuals diagnosed with autism under the age of 30 years old (1,358 males, Age: M = 13.8, SD = 5.6) and 1,734 neurotypical (NT) controls (1,282 males, Age: M = 14.5, SD = 5.8). This represents a substantial expansion and refinement of previous cohorts, incorporating data from new independent sites and excluding participants over age 30 to focus on developmental trajectories.

### Clinical Characterization

Full-Scale IQ (FSIQ) scores were available for 1,395 participants with autism (M = 104.39, SD = 18.40) and 1,521 NT controls (M = 111.52, SD = 14.90). FSIQ was significantly lower in the autism group (t = -11.45, p < .001), though averages for both groups remained within the normative range. Clinical symptom measures, available for subsets of the sample, confirmed expected group differences. For the autism group, the mean ADOS [35] Calibrated Severity Score (CSS, N = 519) for autism participants was 6.97 (SD = 2.03), indicating moderate-to-high severity and differing significantly from controls (t = 41.10, p < .001). The mean Social Responsiveness Scale (SRS; Constantino, 2013) Total Raw score for the autism group (N = 698) was 90.93 (SD = 29.22), which was substantially higher than in the NT control group (N = 729, M = 19.94; t = 57.09, p < .001). Demographic characteristics for all three cohorts (updated ENIGMA-ASD, COCORO, and ABCD) are summarized in Table 1.

**Table 1.** Demographic Characteristics of the Study Cohorts.

| Cohort | N<br>(Total) | N (ASD) | N (Control) | Age (Mean ± SD) | % Male | Primary Measures |
| --- | --- | --- | --- | --- | --- | --- |
| Updated<br>ENIGMA-ASD | 3,389 | 1,637 | 1,752 |  |  |  |
| Childhood | 812 | 483 | 329 | ASD: 7.7 ± 1.9 | ASD:<br>~84% | Cortical Thickness |
| (Under 10 years<br>old) |  |  |  | CNT: 7.8 ± 1.9 | CNT:<br>~62% | Subcortical Volume |
| Adolescent/Adult | 2,577 | 1,154 | 1,423 | ASD: 16.1 ± 5.0 | ASD:<br>~82% | Cortical Thickness |
| (10–30 years<br>old) |  |  |  | CNT: 16.2 ± 5.0 | CNT:<br>~76% | Subcortical Volume |
| COCORO-ASD | 5,604 | 206 | 3,068 | ASD: 28.5 ± 10.9 | ASD: 82% | Cortical Thickness |
|  |  |  |  | CNT: 38.3 ± 14.2 | CNT: 55% | Subcortical Volume |
|  |  |  |  |  |  | White Matter (DTI) |
| ABCD | 4,201 |  | 4,201 | 9.49 ± 0.51 | 54% | RVI Evaluation |
|  |  |  |  |  |  | (Genetics, Cognition) |

*Note.* The ABCD analysis focused on a population-based sample of developing children; CNT = neurotypical control participants.

#### 2.2 Cross-Cohort Reproducibility (COCORO-ASD)

Data for the COCORO sample were derived from two separate mega-analyses: cortical thickness and surface area comparisons were obtained from Matsumoto et al. [22], which included 206 individuals with autism (Mean Age = 28.5, SD = 10.9 years; 18.0% Female) and 3,068 NT controls (Mean Age = 38.3, SD = 14.2 years; 45.4% Female). Subcortical volume comparisons were obtained from Okada et al. [23], which included a similar cohort of 193 individuals with autism and 3,078 controls. See the Supplementary Methods for full details on studies. The white matter RVI was constructed using regional effect sizes for fractional anisotropy (FA) reported by Koshiyama et al. (2020) following the same within-site adjustment and meta-analytic protocol.

### Study II: Population-Based Investigation (ABCD Study)

#### 2.3 Participants (ABCD Study)

Data for the dimensional analysis were drawn from the ABCD Study, a multi-site longitudinal study of brain development and child health in the United States [37]. The present ABCD sample included individuals (N = 4,201) who met the following criteria to ensure high reproducibility: (1) data collected using a Siemens MRI scanner only, as prior studies identified low reproducibility for data collected using Phillips and GE scanners [38, 39]; (2) evaluation at the same study sites during follow-up visits; (3) completed structural and diffusion MRI scanning; and (4) had genetic data available. Additional disqualifying factors included a history of traumatic brain injury, severe perinatal complications, or uncorrected vision and hearing deficits [40]. The present study was limited to participants who had high-quality (see Fan et al., 2023a for information quality control) structural and diffusion neuroimaging data available at both the baseline (Mean Age = 9.9 SD = 0.6 years) and the 2-year follow-up (Mean Age =12.0 SD = 0.7 years) assessments (further information regarding the ABCD sample is included in the Supplementary Methods.

#### 2.4 Neuroimaging Data Acquisition and Preprocessing

For the ENIGMA cohorts, we utilized summary statistics derived from multi-site data. Site-specific acquisition parameters are detailed in the primary publication and provided in our Supplementary Table 1 [19]. The ABCD imaging data were collected using only Siemens Prisma 3T MRI scanners. For the 3D volumetric T1-weighted MRI session, a matrix of 256 × 256, 176 slices, 1.0 mm isotropic resolution, a repetition time (TR) of 2500 milliseconds, and an echo time (TE) of 2.88 milliseconds were used. Diffusion images were acquired with a matrix of 140 × 140, 81 slices, and 1.7 mm isotropic resolution. Full details of the imaging acquisition protocols are described elsewhere [38] and in our Supplementary Methods.

#### 2.5 Regional Vulnerability Index (RVI) Calculation

The ENIGMA consortium provided the meta-analytical ranks of the severity of differences associated with autism, and other disorders (e.g., SSD, BD, MDD, OCD) on gray matter thickness (33 cortical areas), subcortical volumes (7 structures) and fractional anisotropy (24 major white matter regions) [20, 42, 43, 43–50]. The updated ENIGMA RVI-ASD results from this paper were provided from each working group and have not been added to the ENIGMA toolbox. For ABCD analysis the whole-brain RVI was constructed using updated ENIGMA-ASD cortical and subcortical and COCORO white matter effect sizes. Four dependent variables were selected as primary neuroanatomical outcomes: Gray Matter Thickness RVI, Gray Matter Volume RVI, whole-brain RVI-ASD, White Matter RVI-ASD. The calculations for the unimodal RVIs are included in the Supplementary Methods. Higher whole-brain RVI-ASD values imply that the pattern of regional values more closely followed the regional vulnerability pattern for autism as determined by the ENIGMA meta-analysis. RVI calculation was based on previously established methods [45] using RVIpkg [51]. All statistical analyses were performed in RStudio v2025.12.1+563 [52] and R v4.4.3 [53]. Participants were removed from the analysis if their RVI measurements were identified as statistical outliers, defined as being more than two standard deviations from the sample mean. In addition, participants who were not diagnosed with autism nor neurotypical, were removed from the analysis (See Supplementary Figure 1).

#### 2.6 Genotyping (ABCD Study)

All ABCD participants were genotyped using the Affymetrix Axiom Smokescreen Array. This array was designed to characterize global ancestry variations in addition to the whole genome coverage. Participants who have more than 20% missing rates on genotype calls were excluded from the downstream analyses. Variants that had more than 10% missing rates were excluded as well. Overall, 516,598 genetic variants passed quality control [54]. The genetic data for the ABCD sample (Data Release 2022.09) were obtained from the ABCD 5.0 data release available in the NIH-NDA archive (nda.nih.gov). An empirical kinship matrix was calculated from the ABCD genomic data using weighted allelic correlations (WAC) via the SOLAR-Eclipse software. The statistical power of the resulting pedigree for subsequent heritability analyses was evaluated using the Expected Likelihood Ratio Test (ELRT). We utilized polygenic modeling via SOLAR-Eclipse to partition the variance of baseline and longitudinal RVI changes into additive genetic, environmental, and gene-by-age interaction components. Genetic correlation analyses were additionally employed to interrogate these components and screen potential environmental contributors, while empirical heritability estimates derived from genomic data were validated against self-reported relationships. Full methodological details are provided in the Supplementary Methods. Linear mixed-effects models were applied to covariate-adjusted, inverse-normalized data to account for genetic relatedness among subjects. Overall phenotypic correlations between traits were partitioned into shared additive genetic and residual environmental components to identify shared genetic influences. Full methodological details and equations are provided in the Supplementary Methods. We modeled developmental changes in additive genetic control (h^2^) to differentiate the effects of shifting environmental variance from biologically programmed genotype-by-age (GxA) interactions. GxA interactions were quantified as the additive genetic variance in response to development (*σ*^2^*GΔ*) between baseline and follow-up assessments. Full methodological details and equations are provided in the Supplementary Methods.

#### 2.7 Behavioral and Clinical Measures (ABCD)

Environmental factors were extracted from the ABCD Parent Demographic Survey and the developmental history questionnaire administered at baseline. ‘Birth complications’ was a composite score of adverse perinatal events and delivery method (e.g., C-section) was derived from the medical history questionnaire. ‘Lead exposure’ was assessed via the ‘Lead Risk’ variable provided by the ABCD study, which is derived from geocoded data estimating the age of housing stock in the participant’s census tract (a standard proxy for lead paint exposure). Youth-reported negative life events were assessed using the ABCD Life Events Scale (PLE), quantifying the total number of adverse experiences reported by the participant. The ABCD study also captures Income-to-Needs Ratio, calculated by dividing the total household income by the federal poverty threshold corresponding to the household size. Financial difficulty was assessed using a cumulative adversity score derived from the ABCD Parent Demographics Survey. Dimensional measures of psychopathology were assessed using the Child Behavior Checklist (CBCL) Syndrome Scales (e.g., ‘Social Problems’, ‘Anxious/Depressed’, ‘Withdrawn/Depressed’) and DSM-Oriented Scales (e.g., ‘Depressive Problems’, ‘Anxiety Problems’). Social traits were measured using the brief Social Responsiveness Scale (SRS), which was administered to parents starting at the Year 1 assessment. For a full list of measures see the Supplementary Methods.

##### Psychiatric and Behavioral Measures

Parents reported on their child’s current and lifetime mental health using the computerized Kiddie Schedule for Affective Disorders and Schizophrenia for DSM-5 (KSADS-COMP). Given that the ABCD study protocol excluded participants with severe autism unable to comply with scanning procedures (Barch et al., 2021), and did not use gold-standard observational instruments (e.g., ADOS), our analysis prioritized specific symptom dimensions over categorical diagnostic label to capture phenotypic variance within this population-based sample. Notably, the ABCD study transitioned from KSADS-COMP version 1.0 (used at Baseline through Year 2) to version 2.0 at the 3-Year Follow-up. This update was implemented to correct scoring algorithms and enhance the sensitivity of specific modules, including autism (Barch et al., 2021). In our specific analytic sample, no participants met full diagnostic criteria for autism as defined by the KSADS-COMP. However, we interpret this finding within the contents of the instrument’s methodology as a computerized, parent-report tool, which limits the clinical and observational sensitivity to capture the full spectrum of the autism diagnosis. To comprehensively characterize ASD-related phenomenology, we derived three categories of variables from the KSADS autism spectrum disorder supplement, including diagnostic categories, dichotomous symptom dimensions, and continuous measures and developmental trajectories. For a comprehensive list of all specific variable names, item descriptions, scoring criteria, and derivation formulas, please see Supplementary Methods.

### Cognitive Measures

Cognitive measures included the NIH Toolbox Cognition Battery [55]. This battery included seven tasks that were summarized into three composite scores: a Fluid Cognition Composite (derived from flanker, card sort, picture sequence memory, list sorting, and pattern comparison tasks), a Crystallized Cognition Composite (derived from picture vocabulary and oral reading tests), and a Total Cognition Composite (derived from all seven tasks).

#### 2.8 Statistical Analysis

##### Study I Analysis

To assess the stability of the autism neuroanatomical signature, we performed independent samples *t*-tests comparing RVI scores between autism and NT groups in the updated ENIGMA dataset. To control for the family-wise error rate associated with multiple comparisons across the 40 neuroanatomical regions of interest (33 cortical regions and 7 subcortical volumes), a Bonferroni correction was applied. Accordingly, the threshold for statistical significance in the regional follow-up analyses was strictly defined as p < .00125 (α = .05 / 40). We also examined effect size for RVI-ASD by separating the ENIGMA sample into adolescent-to-young adults (age=10-30; N = 2,577) and children (age=under 10 years old, N=812). We further evaluated the cross-cohort reproducibility of this signature by calculating the Pearson correlation coefficient between the regional effect sizes derived from our updated ENIGMA sample and those reported by the independent COCORO consortium.

##### Study II Analysis

In the ABCD cohort, we investigated the relationships between the four RVI metrics (Cortical, Subcortical, White Matter, and whole-brain) and the selected independent variables (environmental factors, cognitive measures, and clinical phenotypes). Our analytic framework was structured to first characterize the RVI-ASD against known factors associated with autism traits and behavioral correlates, and then to examine its longitudinal stability and predictors of change. Longitudinal reproducibility of the RVI-ASD was assessed via the Intraclass Correlation Coefficient (ICC) using the irr package (version 0.84.1) in R. We reported the single-measure ICC using a two-way model for absolute agreement (Model: twoway; Type: agreement; Unit: single), which provides an index of reliability that reflects both the degree of correlation and the agreement between measurements [56]. Independent samples t-tests were conducted to examine the relationship between specific autism symptoms/diagnoses and whole-brain RVI scores. Significant findings are reported such that higher RVI scores (indicating greater neuroanatomical vulnerability) were observed in the group endorsing the symptom compared to the group without.

## Results

### Study I: Autism Brain Signature

#### Autism brain pattern in updated ENIGMA and COCORO samples

The cortical and subcortical effect sizes for the updated ENIGMA-ASD sample are shown in Figure 1 and Supplementary Figure 3. We observed significantly (after Bonferroni correction) lower cortical thickness of the entorhinal cortex (d = -0.23, p = 2.96 x 10^-10^), the fusiform gyrus (d = -0.17, p = 2.50 x 10^-6^), and the middle temporal gyrus (d = -0.15, p = 2.64 x 10^-5^; see Supplementary Table 2).

**Figure 1.**
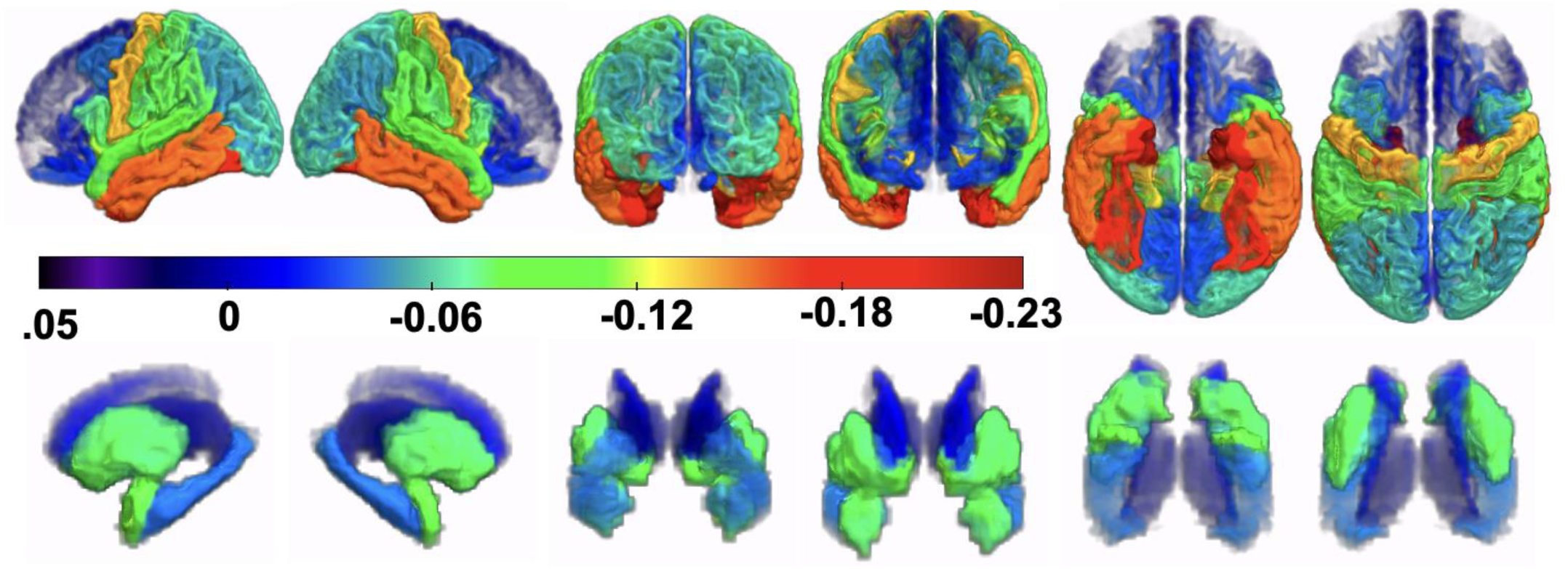
The neuroanatomical signature of autism. Surface projection maps illustrating the regional effect sizes for cortical thickness and subcortical volume differences between autism cases and neurotypical controls in the updated ENIGMA-ASD consortium sample. The color scale represents the magnitude of the decrease (blue/cool) or increase (red/warm) in autism. This stable, replicable pattern serves as the clinical template for the RVI-ASD, capturing the characteristic “brain signature” of the disorder across the lifespan.

Regional effect sizes in the updated ENIGMA sample were significantly correlated (r = 0.88, p = 3.79 10^-14^) with findings reported in the previous study (Van Rooij et al., 2018; see Supplementary Figure 4; Supplementary Table 3). The updated ENIGMA-ASD effect sizes showed concordance with the smaller (N=206 individuals with ASD) COCORO consortium results across all regions (r = 0.40, p = .009; Supplementary Figure 5). Cortical thickness alterations were significantly correlated between the updated ENIGMA and COCORO datasets (r = 0.35, p = .045; Supplementary Table 3). Subcortical volumetric effects showed a strong positive correlation (r = 0.66, p = .074; Supplementary Table 3).

#### Effect Size of RVI-ASD in the ENIGMA Case-Control Sample

Whole-brain RVI-ASD (cortical and subcortical) was significantly elevated in autism individuals with larger effect sizes than any individual neuroimaging measure (t=8.04, p<1×10^-15^, d=0.27; see Supplementary Table 2) in the updated ENIGMA sample. In the adolescent-to-young adults (Ages 10-30), the whole-brain RVI-ASD was also significantly elevated in autism individuals (t = 7.60, p<1×10^-13^, d = 0.30; Figure 2; Supplementary Table 4). In the younger cohort under 10 years old), the whole-brain RVI-ASD was significantly elevated in cases (t = 2.90, p = 0.004, d = 0.20) (Supplementary Figure 6; Supplementary Table 5).

**Figure 2.**
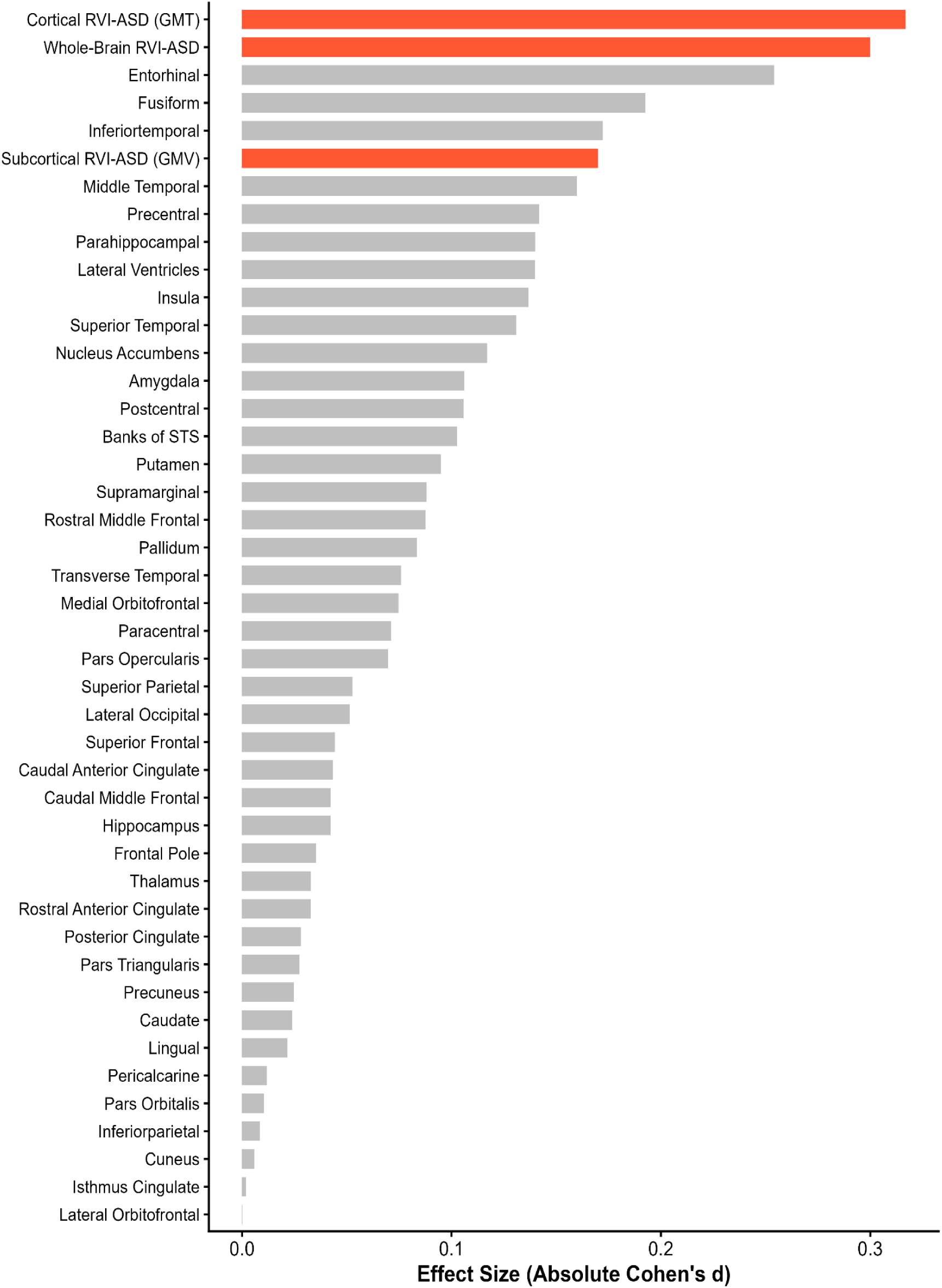
Effect sizes of RVI-ASD and regional brain measures in adolescents and young adults (Ages 10–30). Bar chart ranking the sensitivity (absolute Cohen’s d) of various neuroimaging markers in separating autism cases from neurotypical controls within the adolescent and young adult ENIGMA cohort.

#### Specificity vs. Other Neuropsychiatric Disorders

We calculated the spatial correlation between the group-level Cohen’s d maps for autism and those for other major psychiatric and neurological conditions (Figure 3). The autism pattern showed a significant positive correlation with these for major depressive disorder (MDD, r=0.56, p<10^-6^) and negatively correlated with the difference pattern for anorexia nervosa (AN, r=-0.66, p<10^-9^; Figure 3). In post-hoc analysis, we observed no significant difference in RVI-MDD or RVI-AN for cases vs. controls.

**Figure 3.**
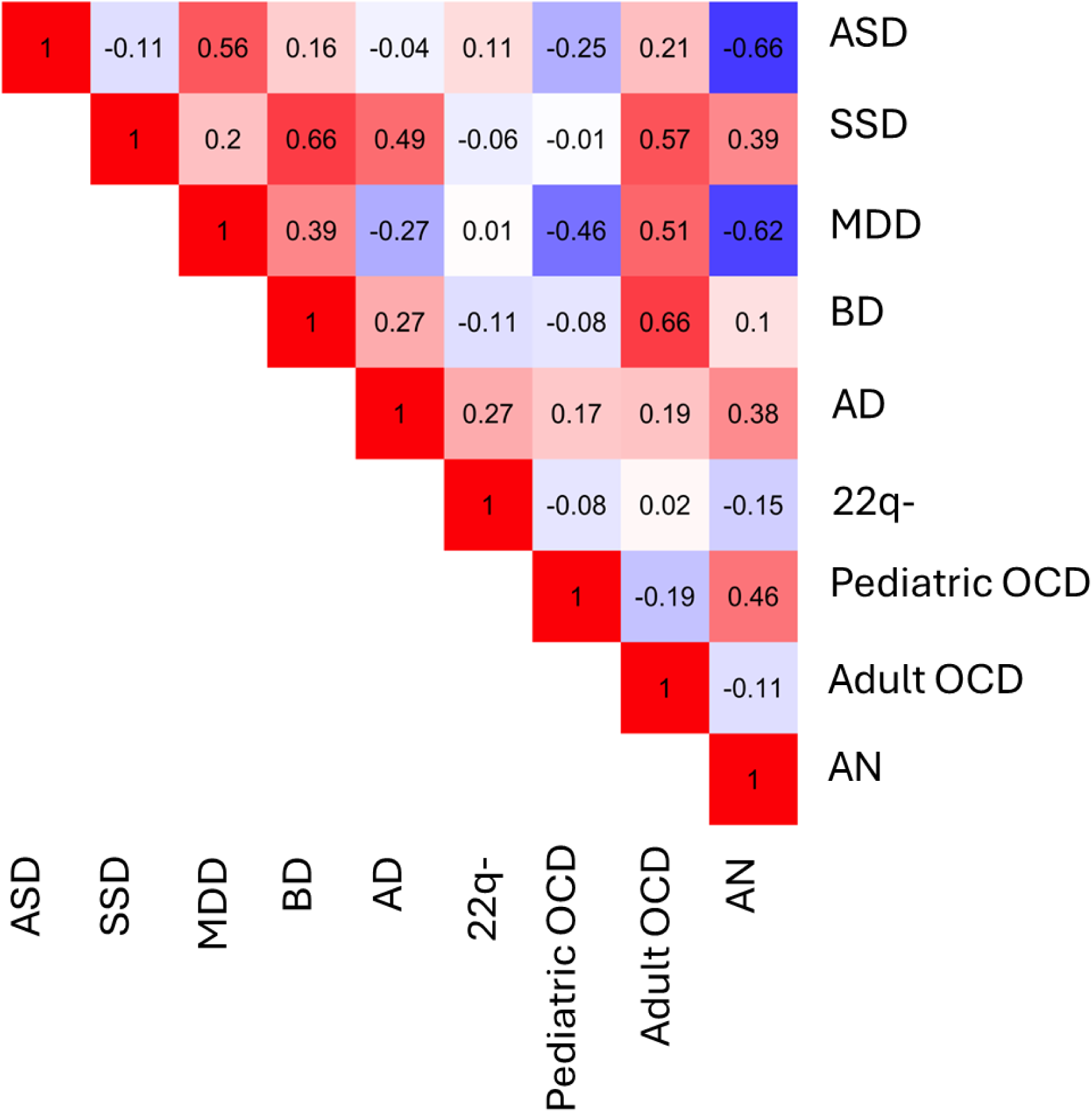
Specificity of the autism neuroanatomical signature across neuropsychiatric disorders. Heatmap illustrating the spatial correlation (Pearson’s r) between the group-level neuroanatomical effect size map of autism and those of other major psychiatric and neurological conditions (Schizophrenia [SSD], major depressive disorder [MDD], bipolar disorder [BD], Alzheimer’s Disease [AD], 22q11.2 Deletion Syndrome [22q], obsessive-compulsive disorder [OCD] in adult and pediatric populations, and Anorexia Nervosa [AN]).

### Study II: Population-Based RVI-ASD analysis (ABCD Study)

#### Longitudinal Stability, Heritability, and Genotype-by-Age Interaction

RVI-ASD pattern was stable and reproducible over the two-year follow-up period in the ABCD cohort (ICC = 0.84). Baseline and follow-up RVI values were significantly heritable (h^2^= 0.83 and 0.83, p<1×10^-16^) and showed no significant genotype-by-age interaction (p=0.488). Tissue-specific analyses showed that cortical and subcortical RVIs were significantly heritable and showed no significant genotype-by-age interaction (Table 2). The White Matter RVI was also heritable and exhibited a significant genotype-by-age interaction (p=0.026).

**Table 2.**
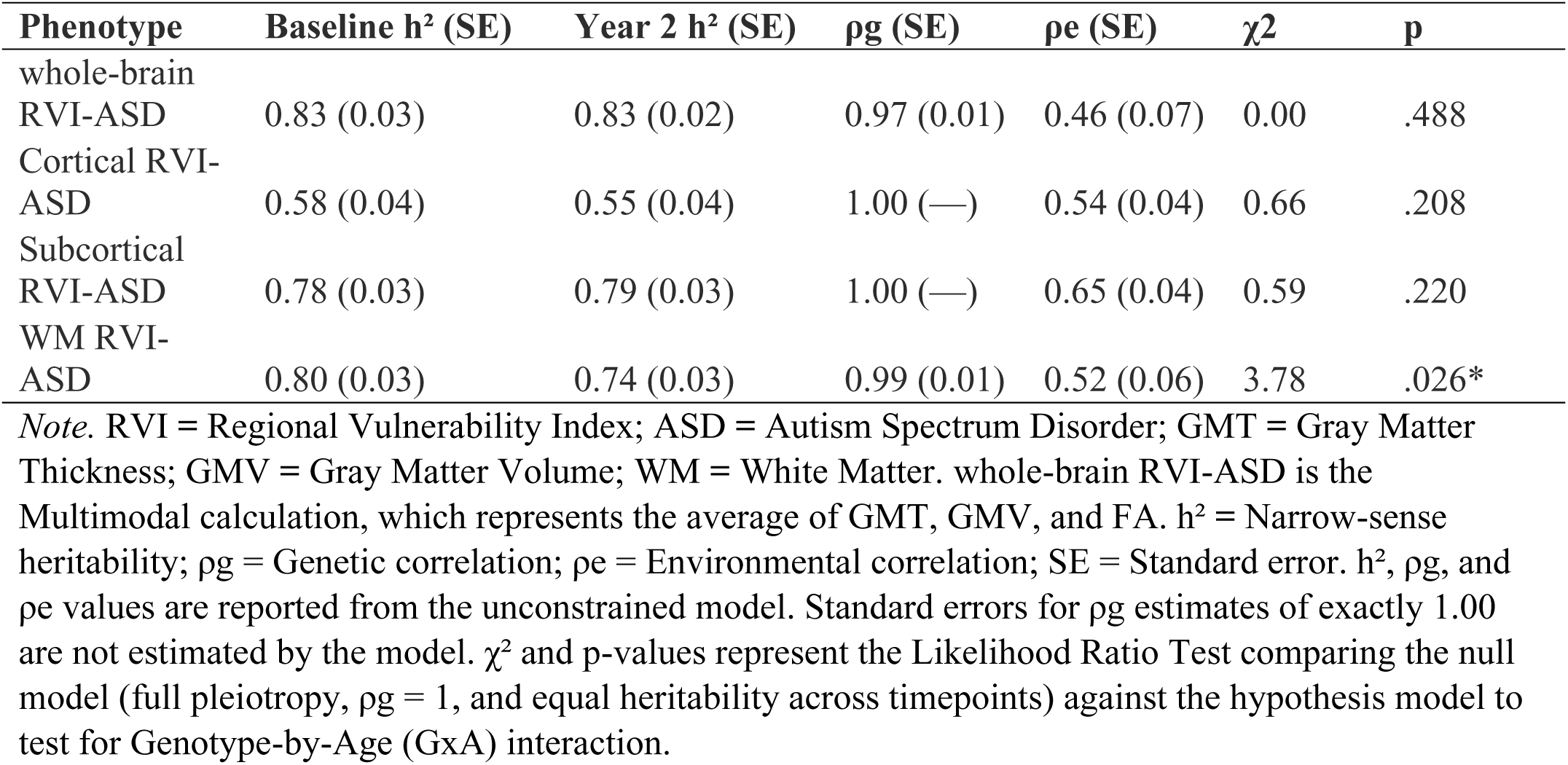
Longitudinal Stability, Heritability, and Genotype-by-Age for RVI-ASD Patterns.

| Phenotype | Baseline $h^2$ (SE) | Year 2 $h^2$ (SE) | $\rho_g$ (SE) | $\rho_e$ (SE) | $\chi^2$ | p |
| --- | --- | --- | --- | --- | --- | --- |
| whole-brain<br>RVI-ASD | 0.83 (0.03) | 0.83 (0.02) | 0.97 (0.01) | 0.46 (0.07) | 0.00 | .488 |
| Cortical RVI-ASD | 0.58 (0.04) | 0.55 (0.04) | 1.00 (—) | 0.54 (0.04) | 0.66 | .208 |
| Subcortical<br>RVI-ASD | 0.78 (0.03) | 0.79 (0.03) | 1.00 (—) | 0.65 (0.04) | 0.59 | .220 |
| WM RVI-ASD | 0.80 (0.03) | 0.74 (0.03) | 0.99 (0.01) | 0.52 (0.06) | 3.78 | .026* |

*Note.* RVI = Regional Vulnerability Index; ASD = Autism Spectrum Disorder; GMT = Gray Matter Thickness; GMV = Gray Matter Volume; WM = White Matter. whole-brain RVI-ASD is the Multimodal calculation, which represents the average of GMT, GMV, and FA. $h^2$ = Narrow-sense heritability; $\rho_g$ = Genetic correlation; $\rho_e$ = Environmental correlation; SE = Standard error. $h^2$ , $\rho_g$ , and $\rho_e$ values are reported from the unconstrained model. Standard errors for $\rho_g$ estimates of exactly 1.00 are not estimated by the model. $\chi^2$ and p-values represent the Likelihood Ratio Test comparing the null model (full pleiotropy, $\rho_g = 1$ , and equal heritability across timepoints) against the hypothesis model to test for Genotype-by-Age (GxA) interaction.

#### RVI-ASD and Risk Factors

Paternal age was associated with significant elevation in individual RVI-ASD (r=0.04, p=.005). Maternal age showed negative but non-significant correlation with RVI-ASD (r=-0.02, p=0.13). We screened for available environmental factors (e.g., birth complications, lead exposure, and neighborhood safety) but found no significant associations. We did find a significant negative association between RVI-ASD and income-to-needs ratio (r = -0.04, p = 0.01) but post-hoc analysis found that paternal age was significantly associated with this ratio (r=0.27, p < 10^-10^) and association with RVI was not significant after correcting for paternal age (Supplementary Figure 7).

#### RVI-ASD, Cognition, and General Psychopathology

We calculated association between RVI-ASD and cognitive performance. RVI-ASD was negatively associated with Total Cognition (r = -0.11, p < 10^-10^), Fluid (r = -0.10, p < 10^-10^), and Crystallized (r = -0.09, p < 10^-10^) intelligence scores (Figure 4). The negative association between RVI-ASD and cognition was primarily driven by cortical and subcortical grey matter (Figure 4) while white matter RVI-ASD was not significantly associated with Total or Fluid cognition, and showed a positive association with Crystallized intelligence (r = 0.04, p = 0.031) and List Sorting working memory (r = 0.04, p = 0.01; see Supplementary Figure 8 for the comparison between groups). In post-hoc analysis, the associations with RVI-ASD and cognition were compared with RVI for SSD and MDD (Supplementary Figure 8) and showed significantly stronger association with cognition than RVI for disorders with known cognitive impact. RVI-ASD was not significantly associated with social problems, depression, or anxiety syndrome scales as measured by the CBCL. Similarly, we found no significant association with withdrawn/depressed symptoms, ADHD problems, or SRS scores (see Supplementary Table 6).

**Figure 4.**
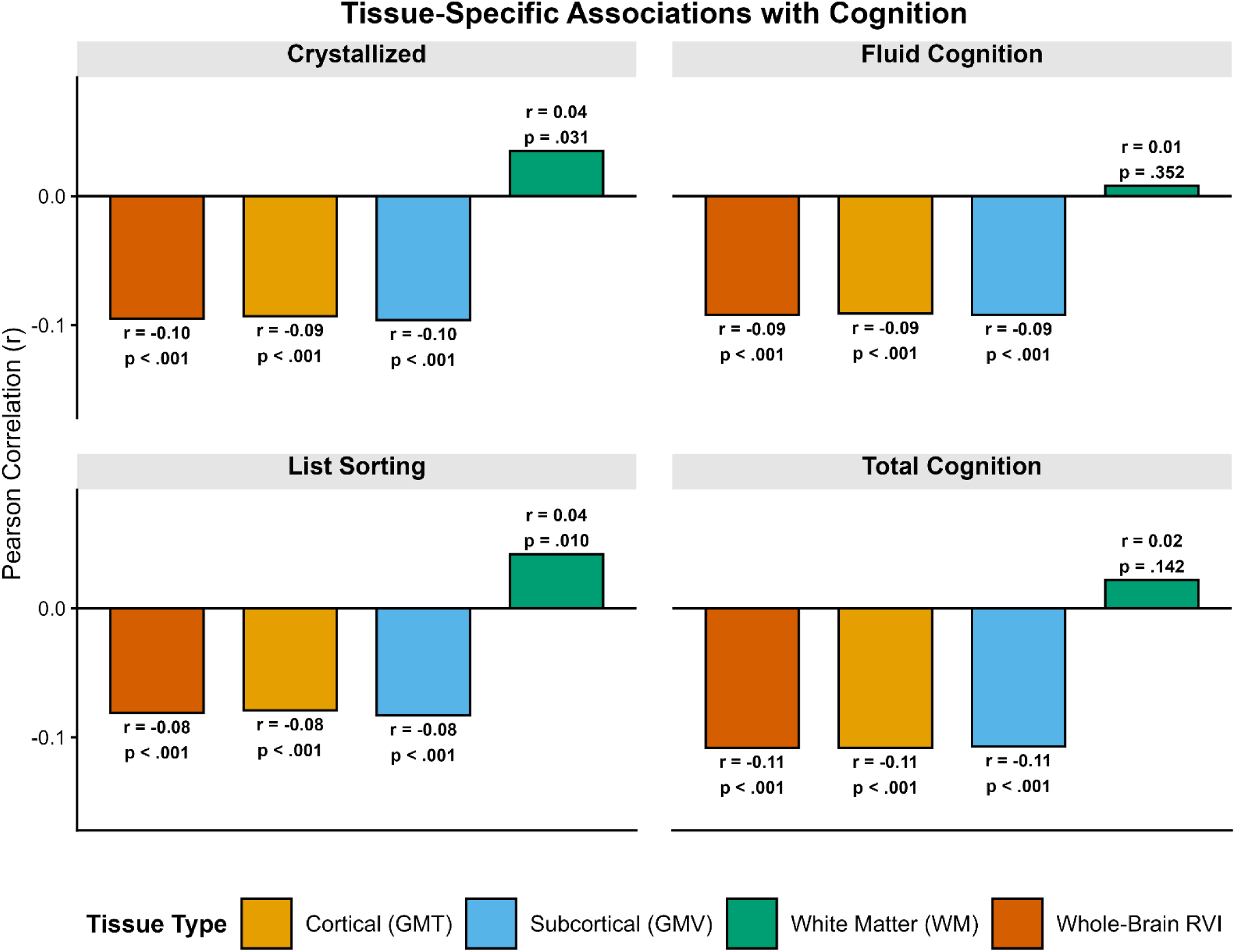
Tissue-specific associations between baseline RVI-ASD and cognitive performance in the ABCD cohort. Grouped bar chart illustrating the distinct roles of cortical thickness (GMT), subcortical volume (GMV), and white matter (WM) similarity indices (calculated at the baseline assessment) in cognitive outcomes.

#### RVI-ASD and autism-linked phenotypes

In the present sample, there were no ABCD participants who were diagnosed with autism based on the KSADS algorithm. However, among the subsample of children whose parents endorsed the initial KSADS autism screening items, significant associations were observed between specific autistic traits and structural vulnerability (Supplementary Table 7). Specifically, children with challenges in social-emotional reciprocity at the Year 4 follow-up (N=55) showed significantly higher whole-brain RVI scores at the Year 2 timepoint (t = 2.34, p = .022). Similarly, a lifetime history of trouble maintaining eye contact reported at Baseline (N=526) was significantly associated with elevated whole-brain RVI scores at Baseline (t = 2.20, p = .028). Furthermore, current poor eye contact reported at the Year 4 follow-up (N=30) was associated with significantly higher whole-brain RVI scores at Year 2 (t = 2.39, p = .023). Post-hoc analyses revealed that a past history of trouble maintaining eye contact was significantly associated with structural alterations localized to the isthmus of the cingulate gyrus (t = 2.21, p = .027; Supplementary Table 8). Current symptoms of poor eye contact were associated with widespread structural alterations, including significant reductions in cortical thickness localized to the entorhinal cortex (t = 2.55, p = .016) and several other regions (see Supplementary Table 8). Post-hoc regional analyses regarding challenges in social-emotional reciprocity did not reveal any statistically significant localized structural alterations (See Supplementary Table 8).

#### Developmental Timing and RVI-ASD

Twenty ABCD subjects had parental report of “Unusual Body Movements” and “Trouble Maintaining Eye Contact”. On average, these parents reported that these symptoms emerged at age 5.16 (SD = 3.15), and the age of onset was significantly correlated with baseline White Matter RVI-ASD (r = -0.61, p = .004; Supplementary Table 9).

Likewise, 82 ABCD subjects had parental report of “Trouble Maintaining Eye Contact” alone and earlier onset (Mean Age = 4.74, SD = 3.15) was significantly associated with White Matter RVI-ASD (r = -0.35, p = .001). Presence of “Strict/Rigid Routines” was associated with more similarity to White Matter RVI-ASD Baseline (N=47 vs 1,900; t = 2.40, p = .020) and Year 2 (t = 2.73, p = .009).

## Discussion

Our study demonstrates that individuals with autism share a pattern of brain differences that is replicable and stable. Individual similarity to this pattern may serve as a neuroimaging-based biomarker for epidemiological samples to study and understand variables associated with autism. We developed RVI-ASD that is a linear and easy to understand index of individual similarity to autism. RVI-ASD provided stronger effect sizes for separating cases from controls than any individual brain structure [57]. We used a population-based cohort of normally developing adolescents (ABCD) to show that this degree of individual similarity is heritable and stable during development. We used RVI-ASD to search for factors contributing to higher similarity and observed that paternal age was only significantly associated factors. In turn, higher RVI-ASD was significantly associated with lower cognitive scores and specific autistic traits. Overall, this study supported the previous finding that autism is a genetic condition with contribution from some environmental factors where higher brain similarity to this illness was associated with significantly lower cognitive scores.

### Study I: Stability and Specificity of the Autism Brain Signature

We observed that effect sizes derived from ENIGMA studies on RVI-ASD (van Rooij et al., 2018; Boedhoe et al., 2020) predicted effect sizes in the updated ENIGMA and independent COCORO cohorts (r = 0.88 and 0.40, respectively). Visual inspection of the replication plot (Supplementary Figure 4) confirmed the agreements in the patterns despite sample-related variability in the magnitude of effect sizes. For example, regional effect sizes in COCORO sample were smaller than in the ENIGMA samples. ENIGMA protocols were used across many neuropsychiatric disorders thus allowing us to compare the patterns of regional effects. Many disorders such as schizophrenia and bipolar disorder co-share 60-80% of regional difference patterns [58–60]. We observed that the patterns in autistic individuals’ brains were unique and did not match those in other mental health disorders. Instead, the autism pattern shared some correlation with the patterns in MDD and AN. It is challenging to interpret this finding as it did not extend to the level of individual tissues. Overall, we conclude that neuroanatomical alterations in subjects with autism are stable and reproducible despite small-to-moderate effect sizes relative to other neuropsychiatric conditions (Okada et al., 2023; Matsumoto et al., 2023).We used this multivariate pattern as the “autism brain signature” to derive the RVI-ASD and to test the “Broader Autism Phenotype” hypothesis that posits that autists traits are widely distributed and can be tested in generate population (Sucksmith et al., 2011).

### Study II: Population-Based Investigation in ABCD

The whole-brain and tissue specific RVI-ASD (ICC=0.72-0.92) were stable from the baseline and two-years follow-up. We calculated the significance of the additive genetic (G), genotype by age (GxA) interaction and environmental (E) and random sources (e) to the variance in RVI. The whole-brain and tissue specific RVI-ASD were significantly heritable for both the baseline and follow-up. The *h^2^*estimate (h^2^=0.83) generally agreed with reported heritability of autism diagnosis in clinical practice (Ramaswami & Geschwind, 2018). We observed significant genotype-by-age interaction for white matter RVI-ASD (p=.026) that likely reflects the protracted and non-linear nature of fiber tract maturation during the transition to adolescence. As highlighted in a comprehensive review by Lebel et al. (2017), association tracts, particularly those facilitating frontal-temporal communication, undergo significant microstructural refinements (e.g., increased myelination and axonal packing) well into young adulthood. This finding aligns with general neurodevelopmental principles which demonstrate that white matter is the most genetically dynamic tissue during the transition to adolescence, with heritability of fiber tract microstructure (e.g., fractional anisotropy) fluctuating according to genetically-timed programs of myelination [61, 62]. This suggests that structural connectivity represents a period of continued neuroplasticity, or vulnerability, where genetic influences on brain wiring are still actively unfolding during the ABCD study’s age range (Lebel & Beaulieu, 2011).

We found a significant association between elevated RVI-ASD and higher paternal age while its association with maternal age was negative and not significant. This specific pattern mirrors findings from large-scale epidemiological studies identifying advanced paternal age as a risk for autism liability [29, 31]. For instance, Reichenberg et al. (2006) found that offspring of men over 40 were 5.75 times more likely to have autism compared to those of men under 30. Sandin et al. (2016) reported that autism rates were 66% higher among children born to fathers over 50, compared to only a 15% increase for children born to mothers in their 40s. The biological mechanisms involve increased rate of de novo genetic mutations in the male germline. Unlike female oocytes, which are formed prior to birth, sperm undergo continuous division throughout the male lifespan, leading to an accumulation of replication-driven genetic errors (Sandin et al., 2016). Conversely, this etiology is likely multidimensional, involving not only germline mutations but also epigenetic and genetic factors associated with delayed fatherhood (Janecka et al., 2017).

Higher RVI-ASD was robustly associated with lower cognitive scores including fluid and crystalized intelligence. The negative association between RVI-ASD and cognition was significantly stronger than that for RVI-SSD and RVI-MDD (see Figure 3). These two disorders were chosen because SSD-like pattern shows strong association with cognition, while subjects with MDD show only modest cognitive differences [63–66]. From a neurobiological perspective, our results support the conceptualization of autism liability as being linked to cognition and consistent with models positing that cognitive differences are the core features of autism [67, 68]. We also observed associations between RVI-ASD and autism-linked phenotypes, including poor eye contact and unusual body movements. We observed that early-onset social-communicative challenges, emerging before age five, were correlated with the White Matter RVI, whereas the broader clinical presentation tracked with the whole-brain RVI-ASD. This aligns with the “Developmental Cascade” model of autism, which posits that early disruptions in white matter development constrain the subsequent functional maturation of social-processing networks (Wolff et al., 2012; Faraji et al., 2023). Specifically, the correlation the timing (earlier) symptoms onset and higher White Matter RVI suggests that white matter integrity may act as a sensitive marker for disruptions occurring during critical windows of rapid myelination in the first years of life. In contrast, the whole-brain RVI likely captures a more global, macro-scale network deviation that emerges as these early connectivity differences propagate across distributed neural circuits. By identifying these links in a community sample, we demonstrate that the RVI-ASD signature is sensitive to the biological markers of autism even in sub-threshold presentations, representing a continuous neuroanatomical phenotype that exists beyond formal diagnostic boundaries.

Higher RVI-ASD robustly predicted lower cognitive performance but were not associated with any general behavioral symptoms (e.g., anxiety, depression). This suggests that the RVI-ASD captures a latent biological liability that scales linearly with cognitive architecture but does not result in overt behavioral psychopathology until a clinical threshold is exceeded. This finding supports the conceptualization of the RVI as a stable, neuroimaging-based biomarker of likelihood of the expression of core autism traits, reflecting the underlying neural substrate of the disorder, rather than a state-dependent marker of current symptom severity.

### Strengths, Limitations, and Future Directions

The primary strength of this study is its use of the ABCD cohort, which provides the statistical power to conduct a dimensional, transdiagnostic investigation of a neurobiological marker. By moving beyond a traditional case-control framework, we uncovered nuanced relationships among genetic liability, brain structure, and cognition across the developmental spectrum. However, several limitations must be acknowledged. First, the RVI is a correlational measure of similarity and cannot be interpreted as a causal biomarker. Second, the two-year follow up period in the ABCD cohort is relatively short; longer term tracking is needed. Third, our study revealed conflicting results between the clinical ENIGMA and general population ABCD cohorts. This highlights a critical need for future research to understand *why* the RVI-ASD behaves differently in clinical versus general population samples. Future research could use the RVI to explore neurobiological heterogeneity within the autism spectrum, examining how the signature varies across individuals with different symptom profiles, cognitive abilities, and etiological backgrounds. The lack of significant environmental associations in our analysis may be attributable to the timing of data collection rather than a true lack of effect. For example, while the ABCD protocol tracks exposures such as lead, these are measured in late childhood (ages 9–10), whereas the critical window for such neurodevelopmental impacts likely occurred substantially earlier. Another limitation is that, while the K-SADS is a validated semi-structured interview, the computerized version relies on parent report rather than direct clinician observation of social behaviors. Consequently, the specific symptom variables utilized in our analysis (e.g., “Poor Eye Contact,” “Strict Routines”) likely offer greater phenotypic granularity for neuroimaging correlations than the categorical diagnostic labels, which were not strictly validated by expert clinical consensus. Furthermore, we must acknowledge limitations regarding the demographic composition of the COCORO autism cohorts, which were predominantly male (e.g., only 18.0% female in the cortical thickness sample). Finally, while the RVI provides a robust macro-scale signature of autism liability, the underlying cellular and molecular mechanisms driving these patterns remain to be fully elucidated. Future studies should aim to bridge this gap by employing “virtual histology” approaches, which integrate neuroimaging data with high-resolution gene expression atlases (e.g., Allen Human Brain Atlas).

## Conclusion

In conclusion, this study establishes the RVI-ASD as a complex neuroimaging-based biomarker that captures the neurobiological liability for autism. By integrating clinical findings from ENIGMA and COCORO with population data from ABCD we show that the autism brain signature is a stable, highly heritable trait that tracks with cognitive performance and autism traits in a population-based sample. This work underscores the power of a population neuroscience approach to reveal the biological pathways involved in neurodevelopmental conditions and provides a robust tool for future research into the dimensional nature of autism. Overall, this study is the first to show that autism is associated with stable and replicable brain differences and that similarity to these patterns can serve as an informative biomarker. In autism cases, similarity index provided stronger effect size than any individual brain region. In typically developing individuals, higher similarity to autism was associated with lower cognitive performance including low scores on the fluid and crystalized intelligence. Our approach allowed us to evaluate the RVI-ASD as a dimensional neuroimaging-based biomarker, confirming that the biological liability for autism is quantifiable even in non-clinical populations (Lenroot & Young, 2013). Our finding aligns with the research which suggests that autism represents an earlier, more generalized perturbation of neural development affecting broad cognitive capacity, whereas SSD-related deficits may emerge during later developmental windows or affect more specific functional domains [65, 69, 70]. These findings suggest that the neuroanatomical patterns characteristic of autism capture a dimension of neural organization that is fundamental to broad cognitive development [71], whereas the patterns associated with other disorders may be more specific to other functional domains or later developmental stages.

Supplementary information is available at MP’s website

## Acknowledgements

LEH has received or plans to receive research funding or consulting fees on research projects from Mitsubishi, Your Energy Systems LLC, Neuralstem, Taisho, Heptares, Pfizer, Luye Pharma, Sound Pharma, IGC Pharma, Takeda, and Regeneron. None of these companies was involved in the design, analysis or outcomes of the study.

This work was supported by the National Institutes of Health grants R01MH123163, R01MH112180, R01MH116948, S10OD023696, R01EB015611, R01MH117601, R01AG095874, R01MH116147, P50MH103222 and U01MH108148. These funding sources provided financial support to enable design and conduct of the study or collection, management, or analysis of the data. None of the funding agencies had a role in the interpretation of the data. None had a role in the preparation, review, or approval of the manuscript. None had a role in the decision to submit the manuscript for publication.

## Conflict of Interest

None to report.

## Footnotes

1 Note that the present study chose to use person- and identity-first language throughout the paper because we wanted to include multiple preferences of the autistic community. We refer the reader to recent articles for more in-depth discussions on this subject (Keating et al. 2023; Singer et al. 2023).

---

**Supplementary Figure 1.**
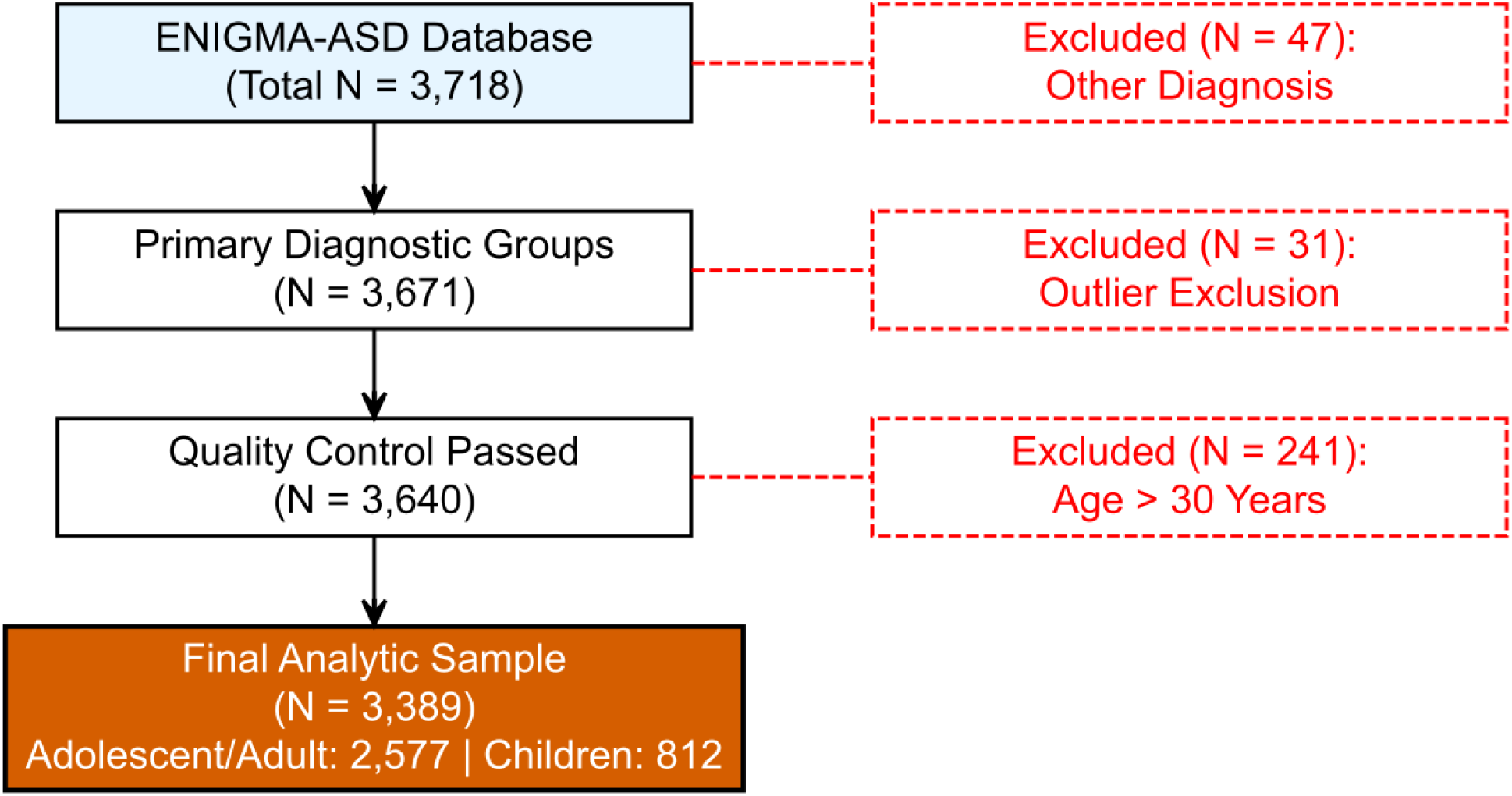
Flow diagram illustrating the step-by-step exclusion criteria applied to the initial ENIGMA-ASD database (N=3,718) to derive the final analytic sample (N=3,389). Exclusions were performed sequentially based on the presence of other primary diagnoses (N=47), exclusion of outliers in the RVI data (N=31), and exclusion of individuals who were greater than 30 years old (N=241). The final sample consisted of 2,577 adolescent/young-adults (10-30 years old) and 812 children (under 10 years old).

**Supplementary Figure 2.**
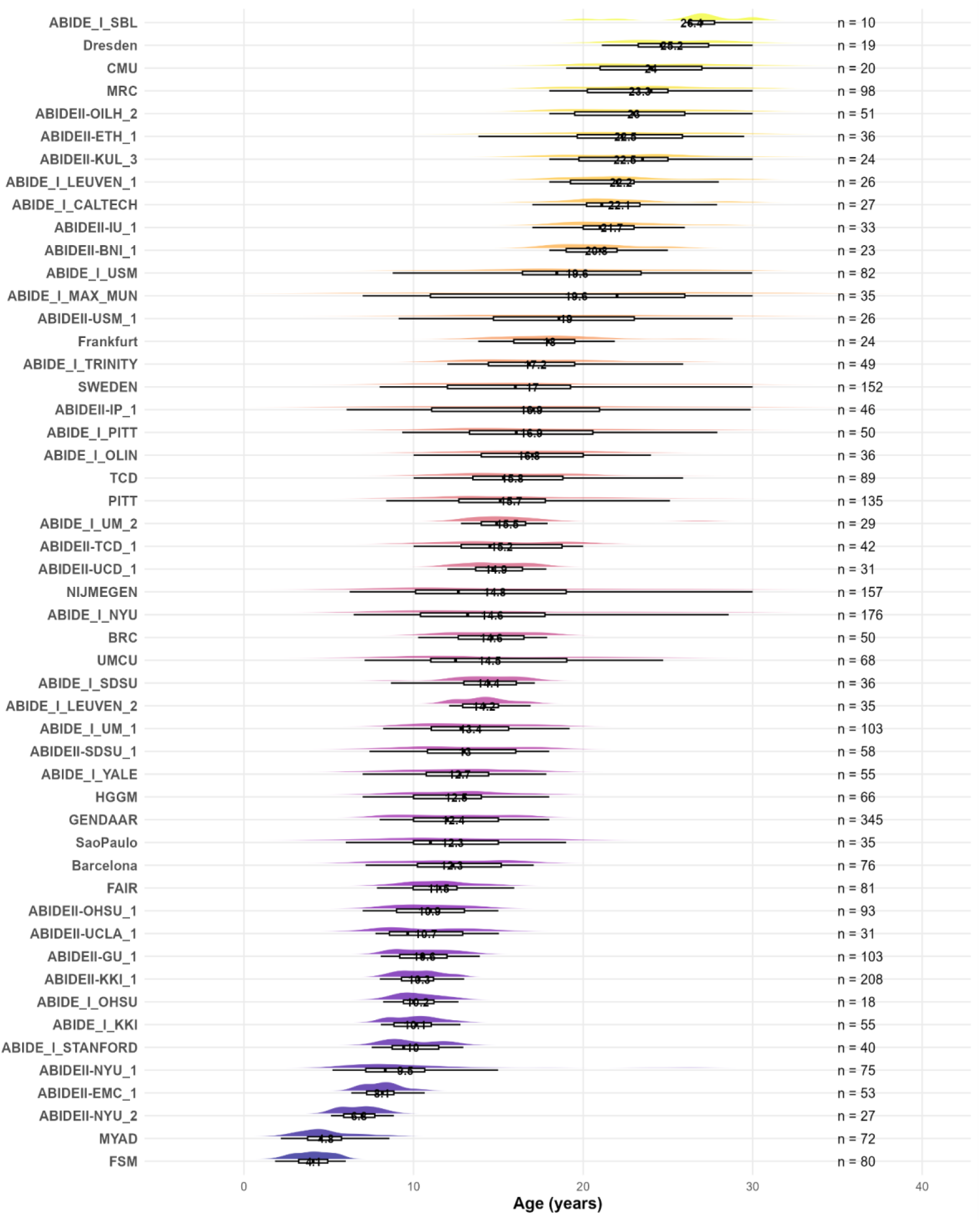
Age distribution across participating sites. Raincloud plots illustrating the distribution of age (in years) for each scanning site included in the final analytic sample. Sites are sorted by ascending mean age. For each site, the visualization includes a half-violin plot representing the density distribution of age and a box plot indicating the interquartile range. The numeric value centered on each box plot denotes the mean age for that site, while the value to the right indicates the total sample size (n) for that specific site. Note that “cattes,” “liehoe,” and “wougro” were combined in the NIJMEGEN site.

**Supplementary Figure 3.**
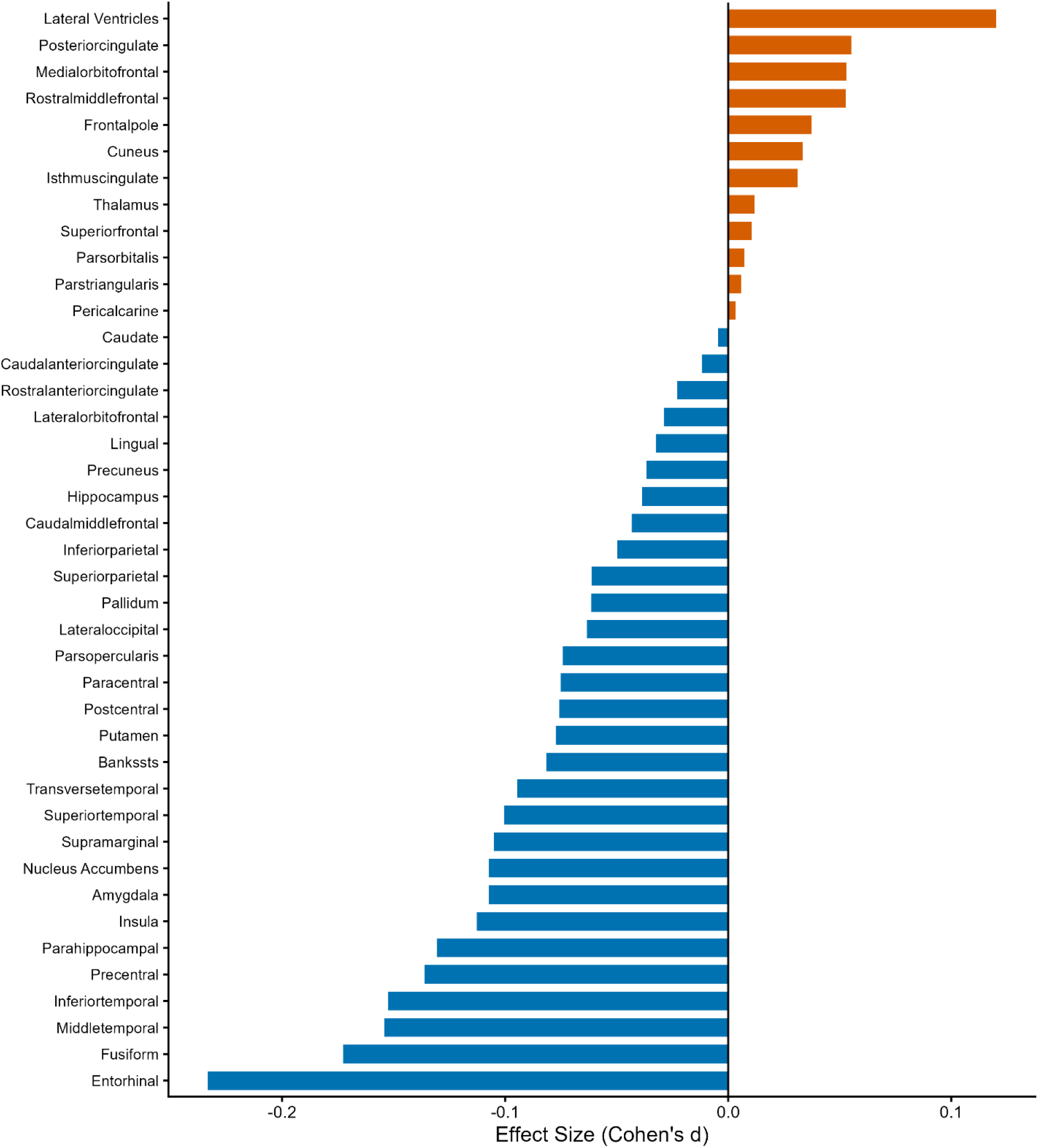
Quantitative profile of the autism neuroanatomical signature. Regional effect sizes (Cohen’s d) for cortical thickness and subcortical volume differences between individuals with autism and neurotypical controls in the updated ENIGMA-ASD analytic sample. Regions are ordered by effect magnitude. Blue bars indicate regions where the autism group exhibits significantly reduced thickness or volume (decreases), while red bars indicate regions of increase (excess).

**Supplementary Figure 4.**
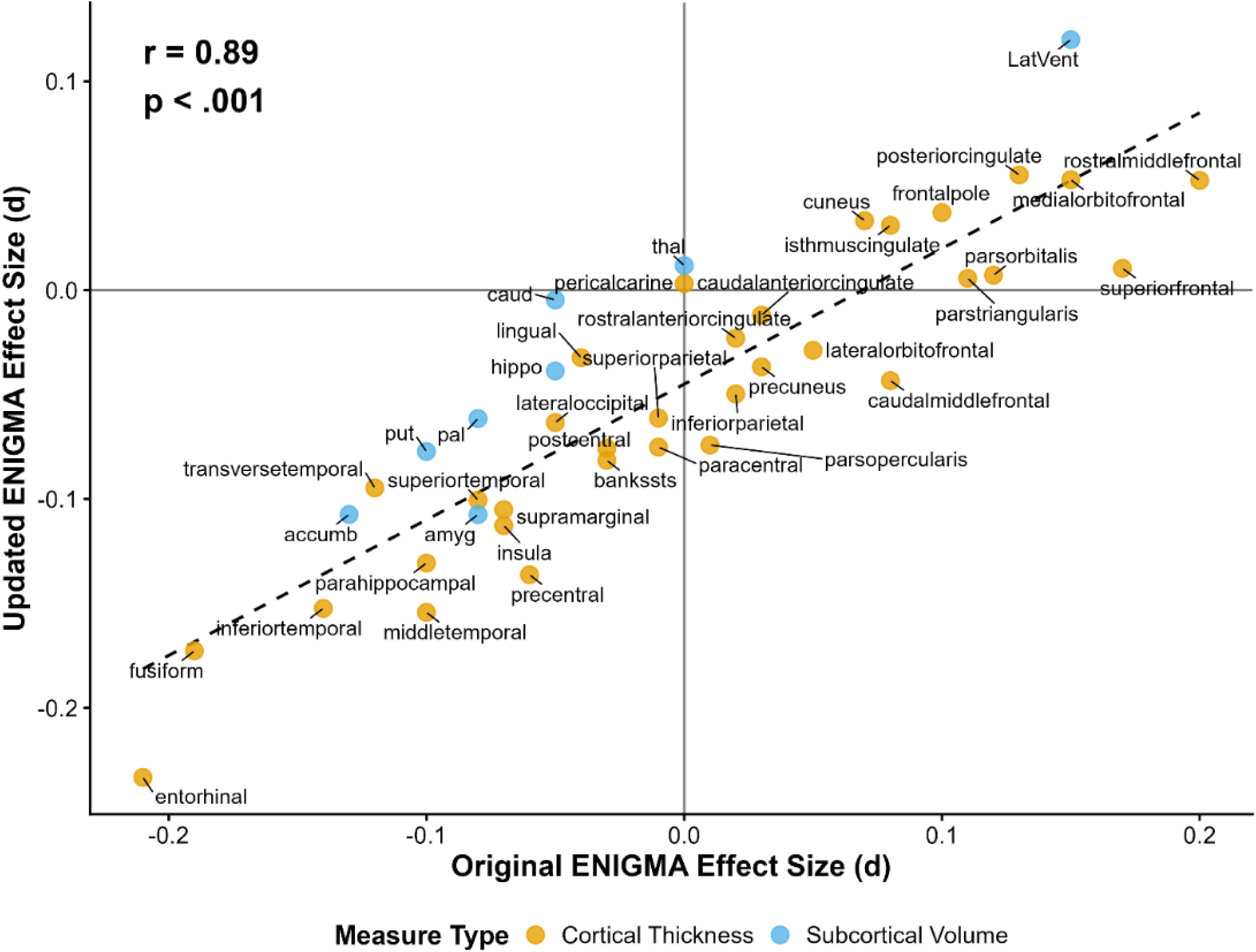
Replication of ENIGMA Findings, comparing effect sizes from the original (van Rooij et al., 2018) and updated ENIGMA findings.

**Supplementary Figure 5.**
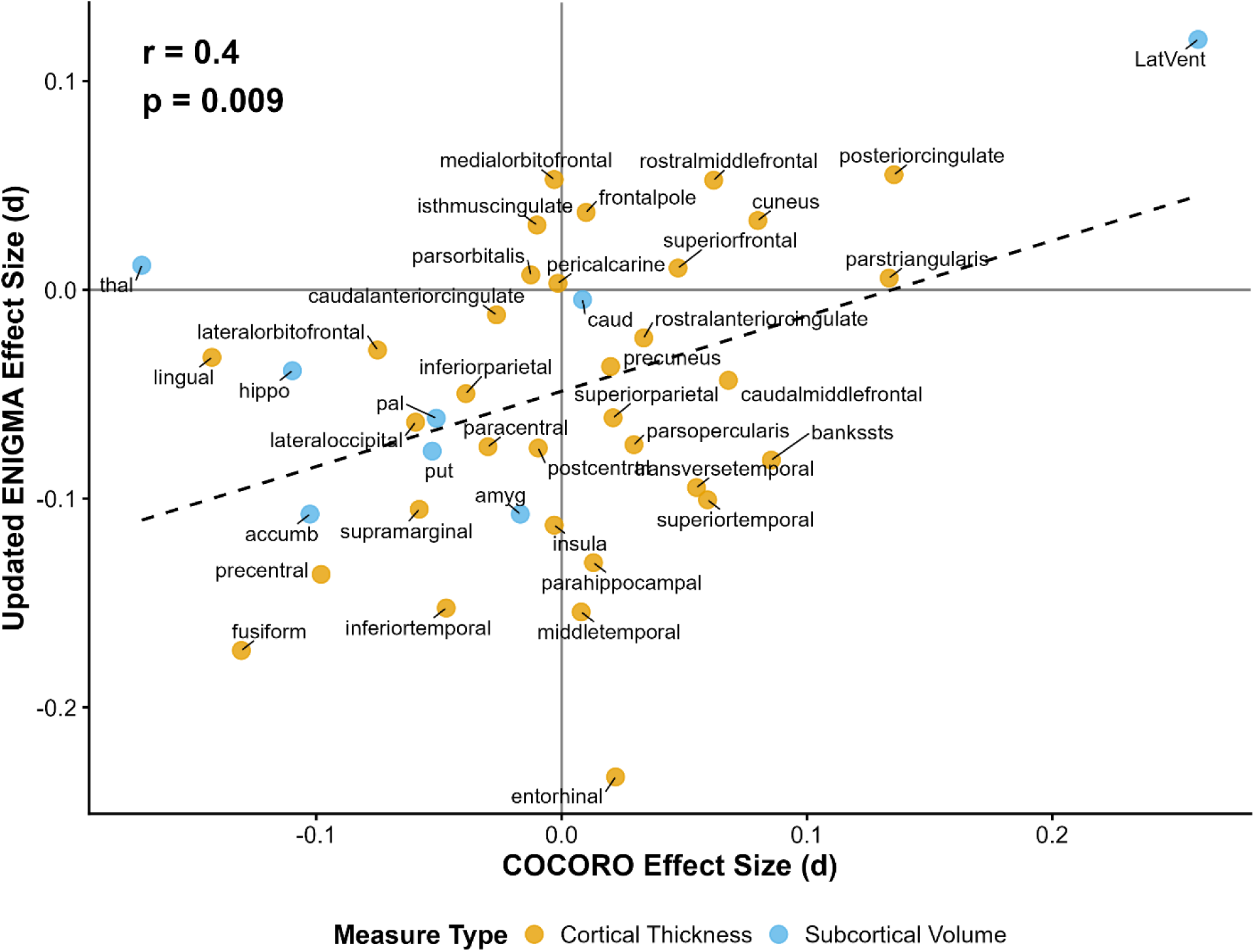
Cross-cohort Replication, comparing effect sizes from the COCORO vs. the Updated ENIGMA data sets.

**Supplementary Figure 6.**
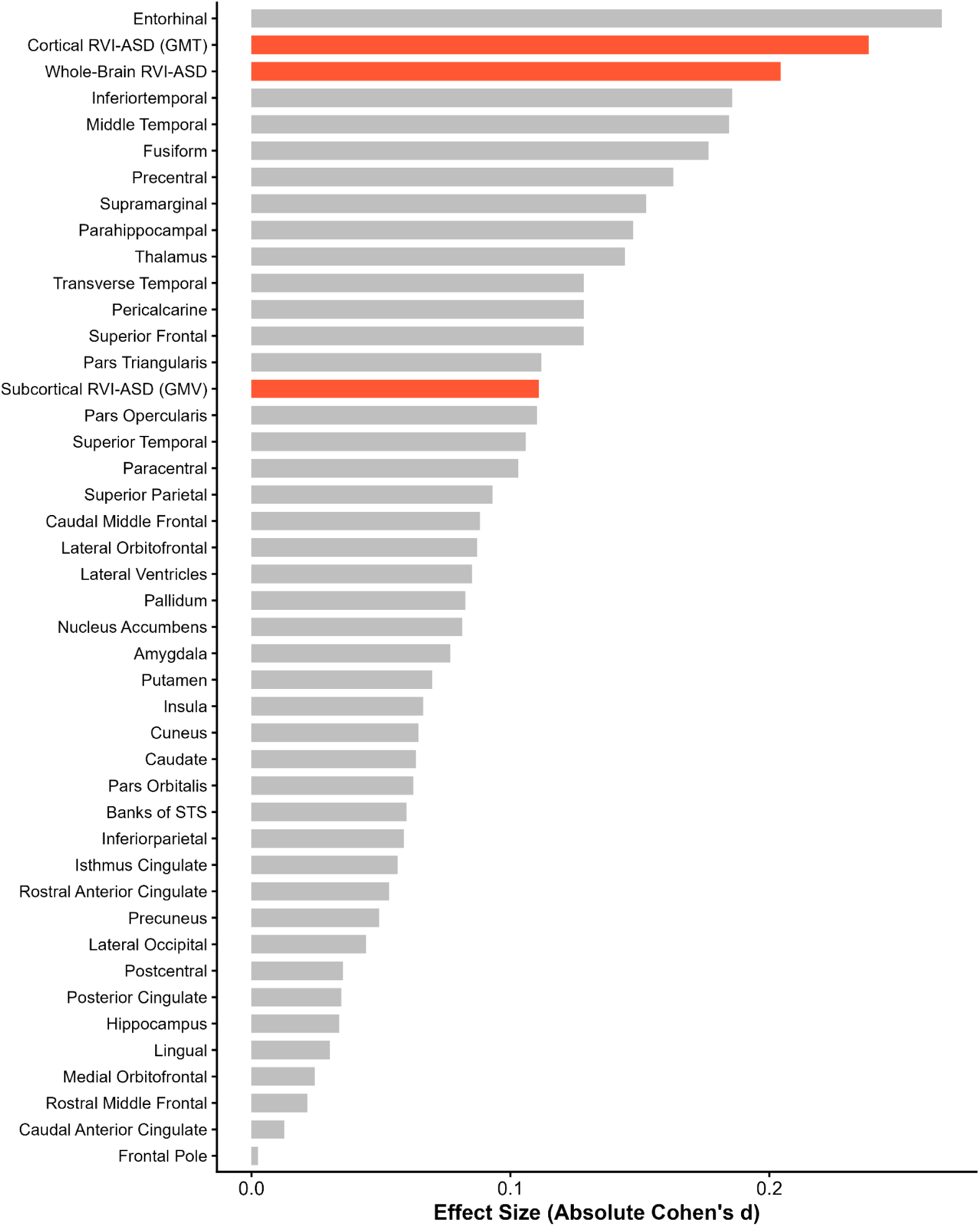
Effect sizes of RVI-ASD and regional brain measures in children (under 10 years old). Bar chart ranking the sensitivity (absolute Cohen’s d) of various neuroimaging markers in separating autism cases from neurotypical controls within the younger ENIGMA cohort.

**Supplementary Figure 7.**
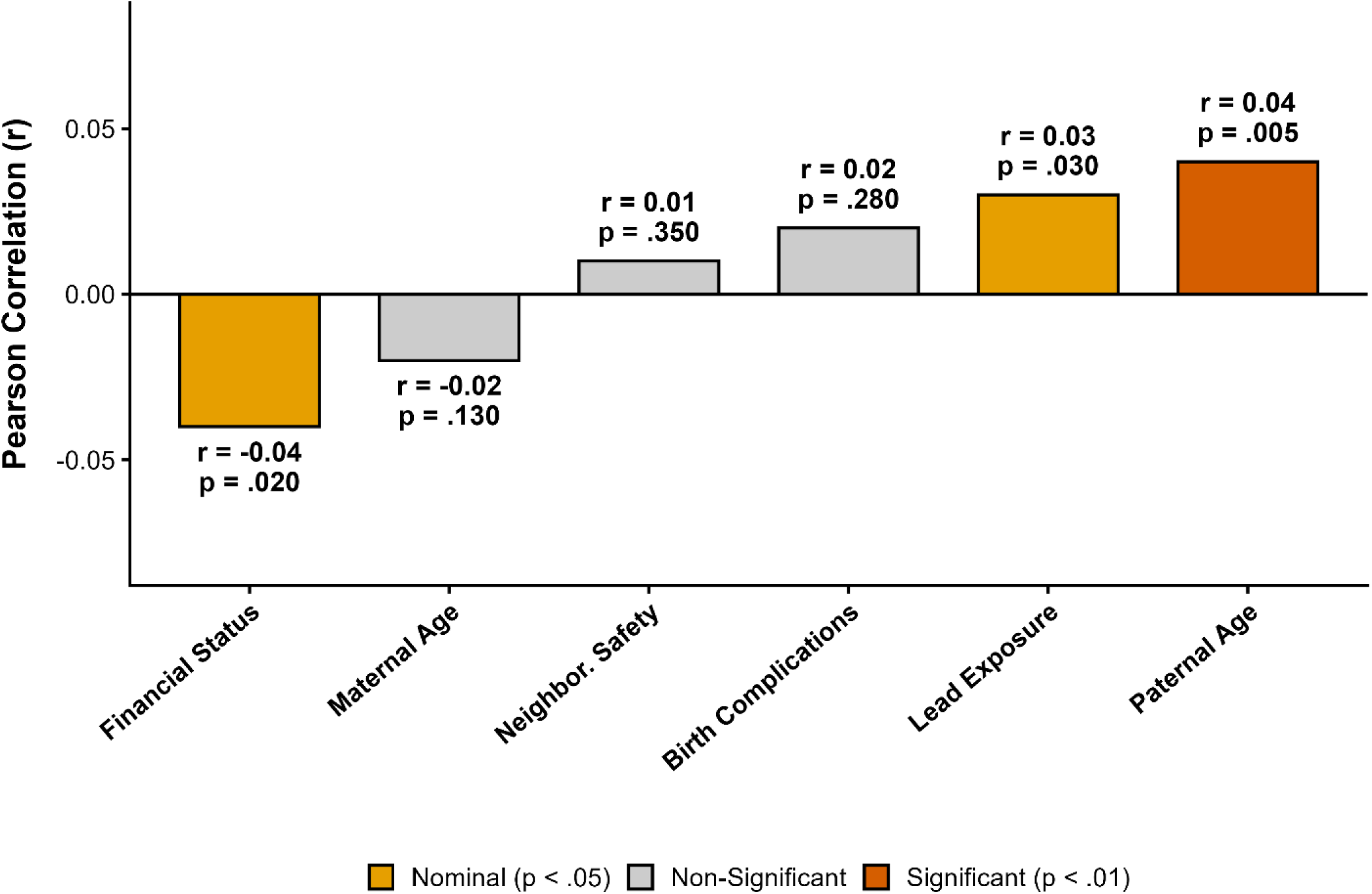
Association of whole-brain RVI-ASD with etiological factors associated with the likelihood of autism in the ABCD cohort. Bar chart representing the Pearson correlation coefficients (r) between the neuroanatomical autism signature and known factors associated with the likelihood of autism.

**Supplementary Figure 8.**
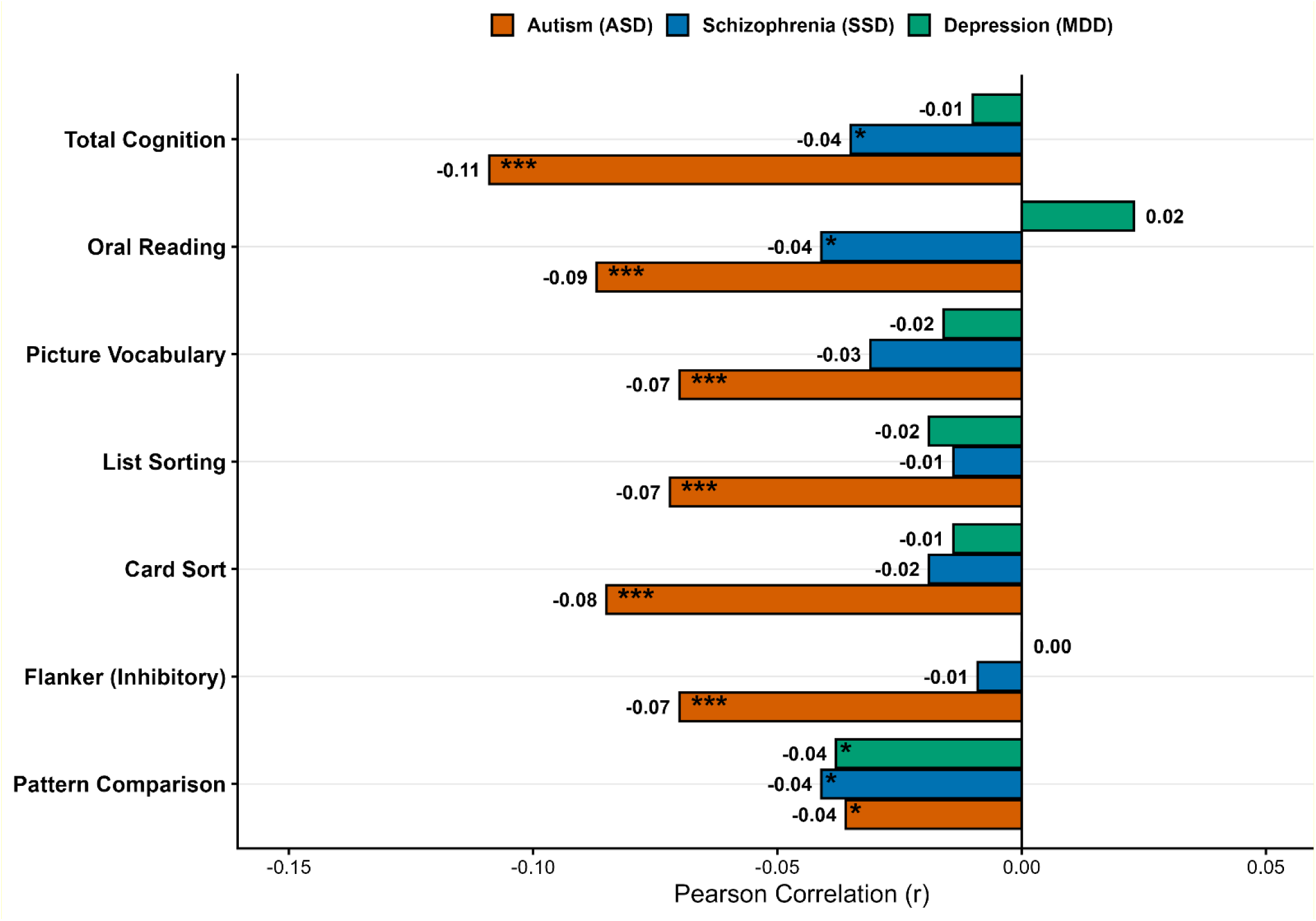
Comparative specificity of brain-cognition associations. Bar chart illustrating the Pearson correlation coefficients for various variables for autism, Schizophrenia (SSD), and Major Depressive Disorder (MDD). p < 0.001 = ***; p < 0.01 = **; p < 0.05 = *.

**Supplementary Table 1.**
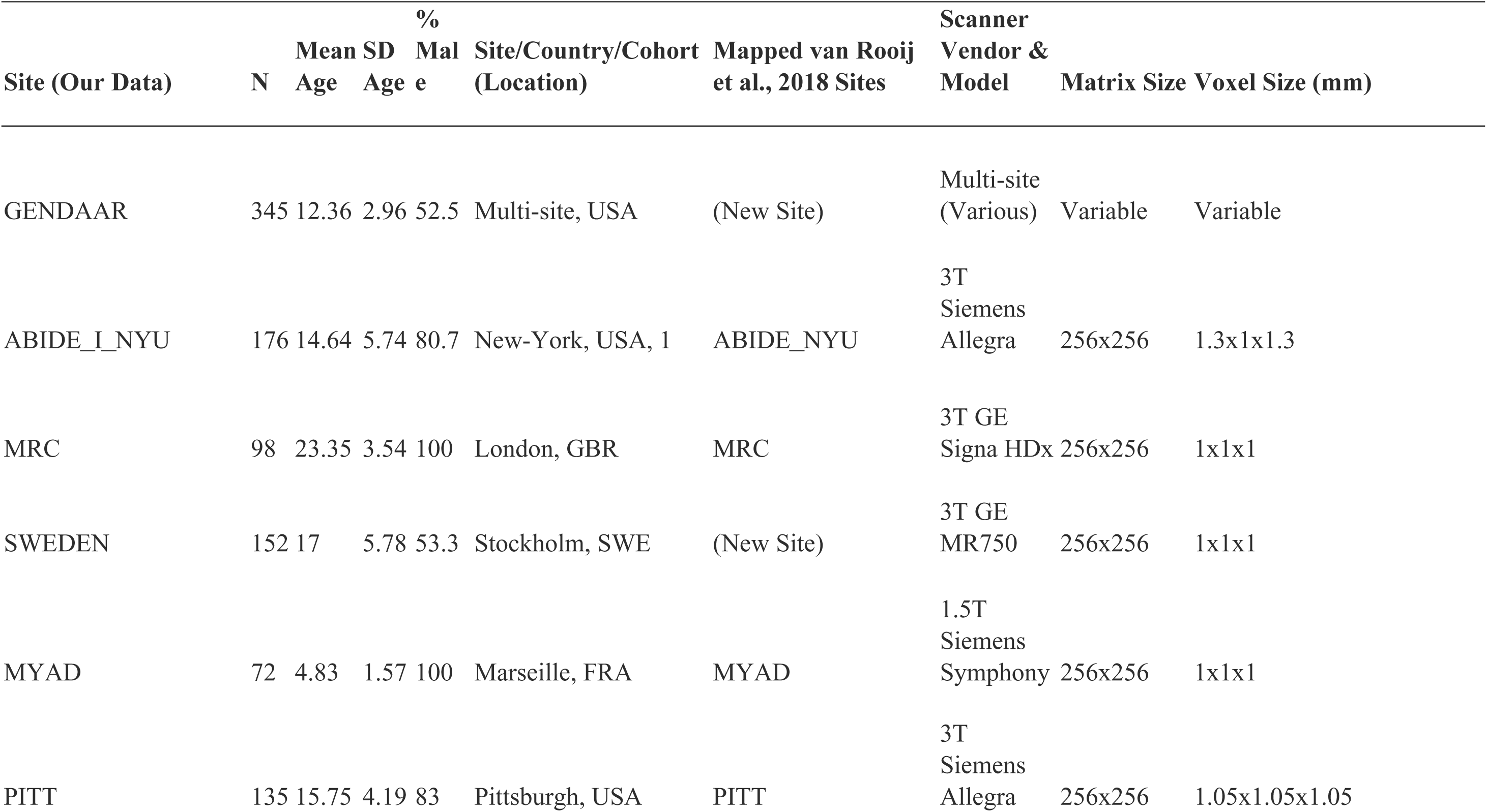
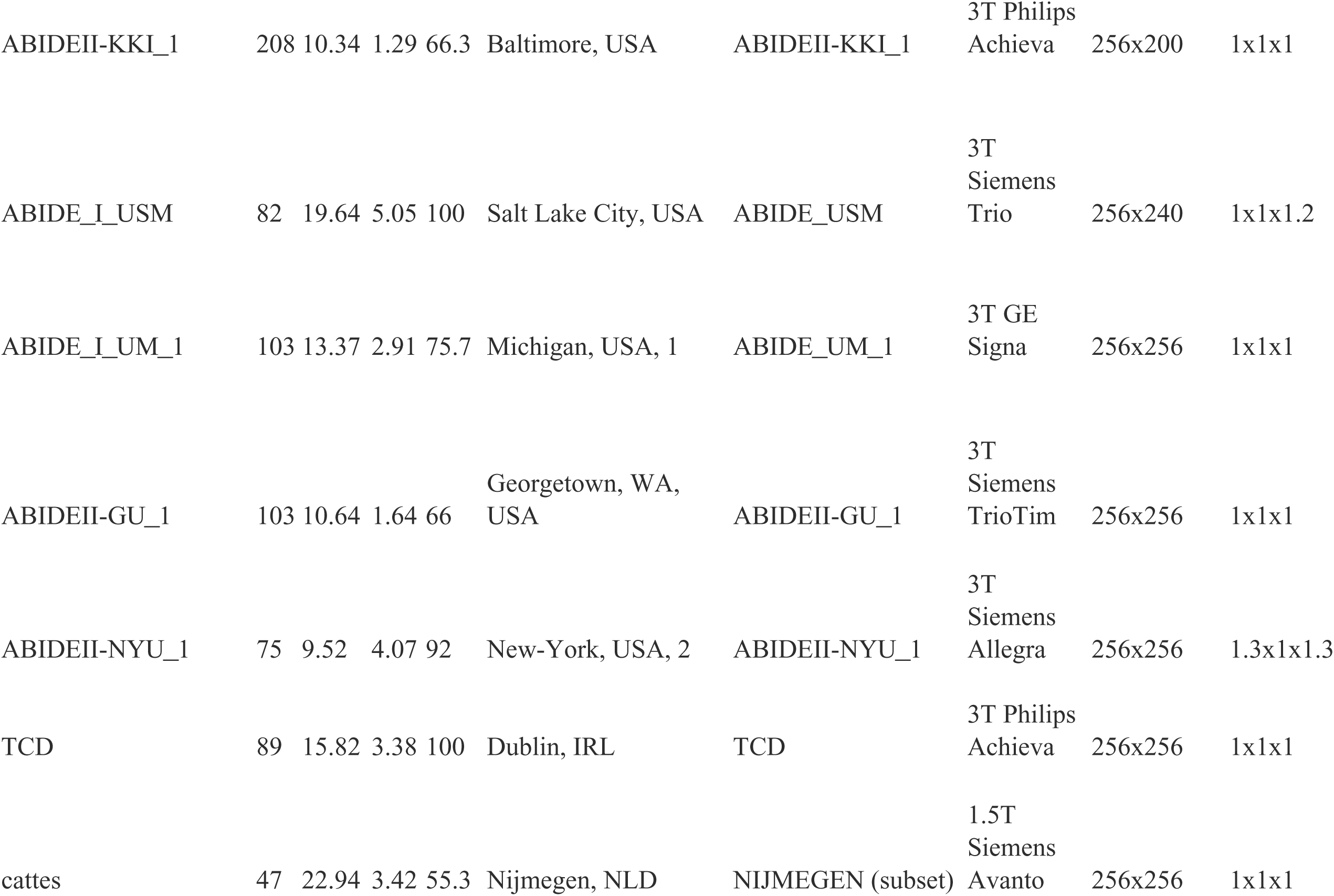
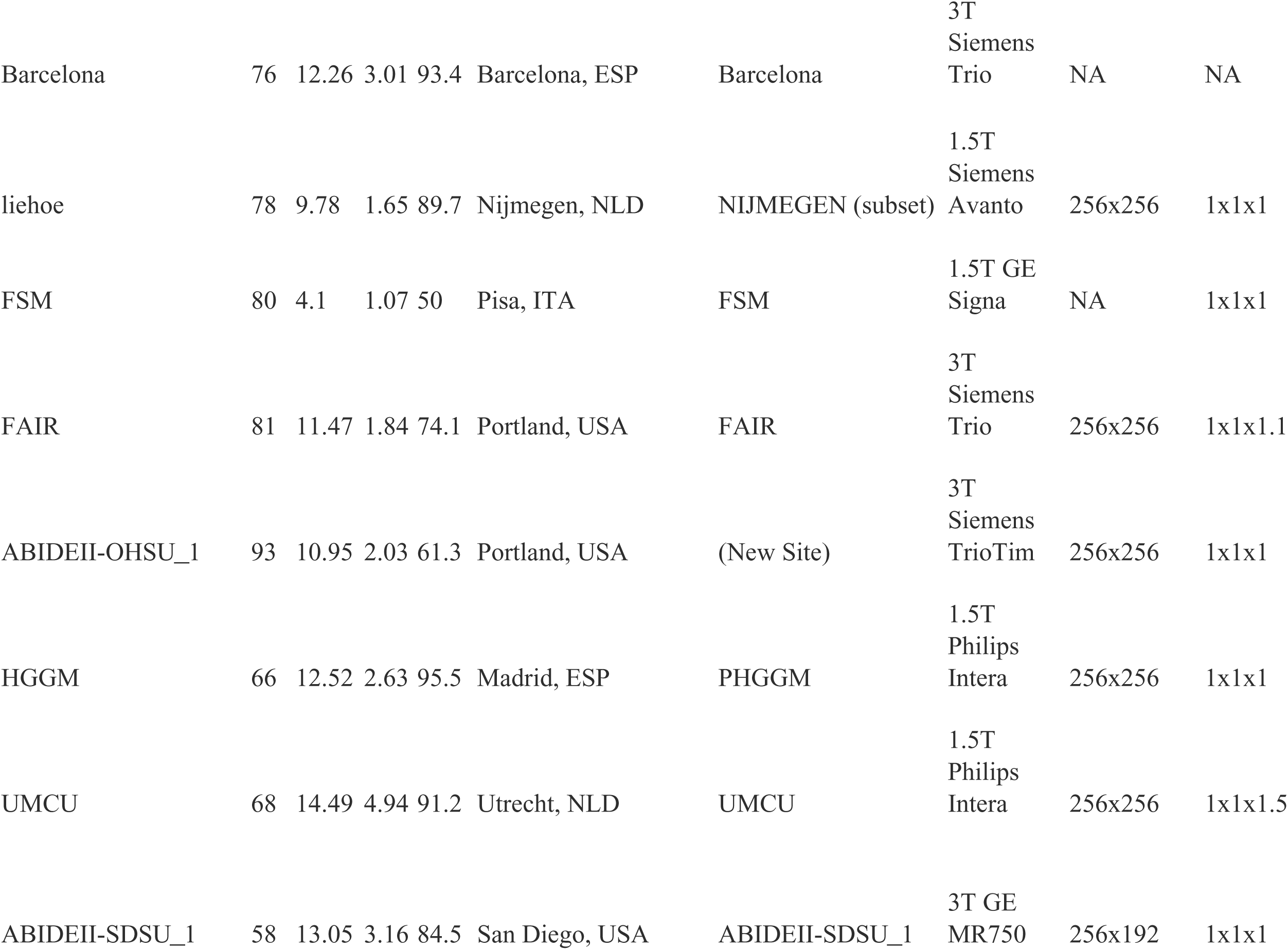
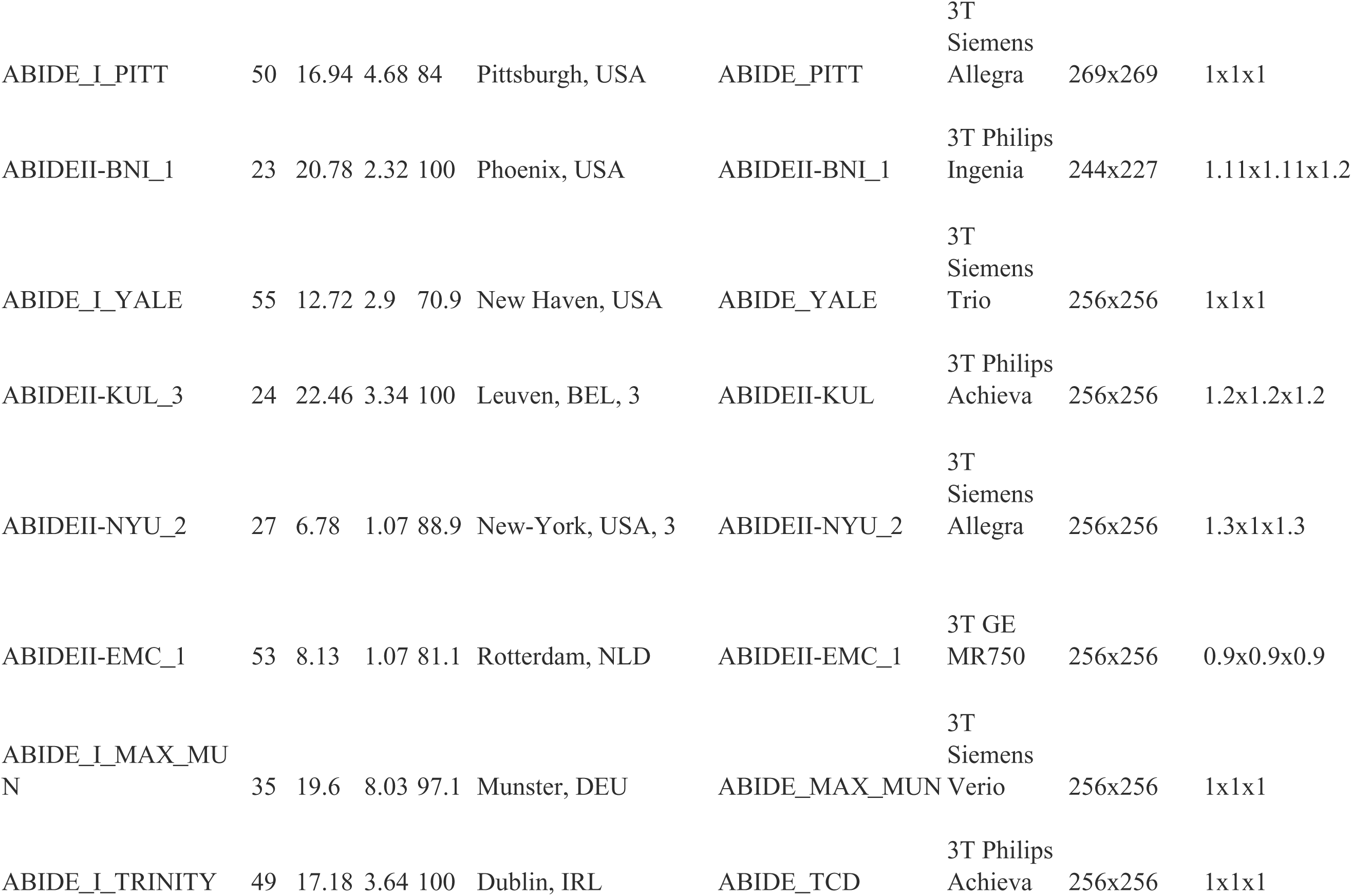
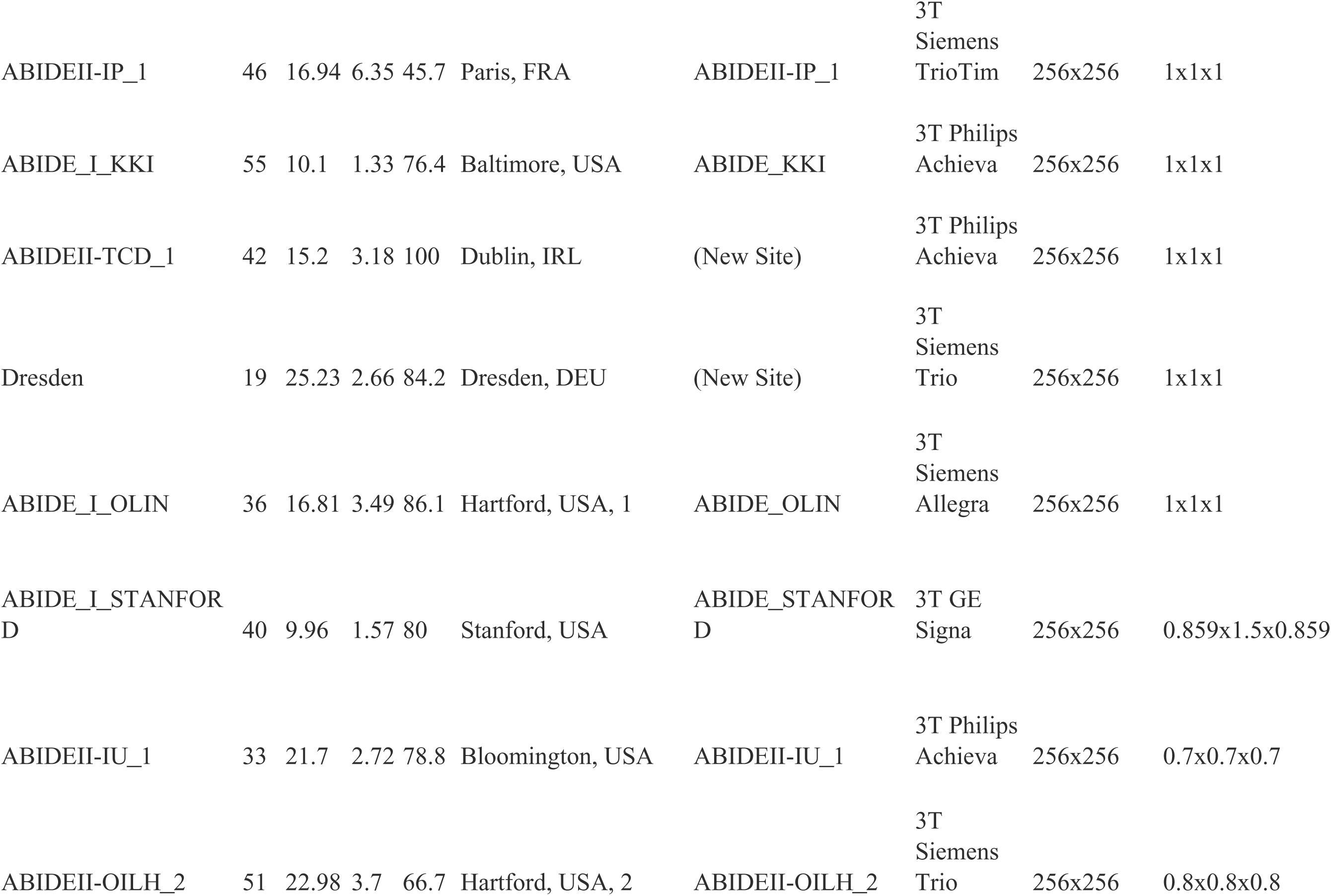
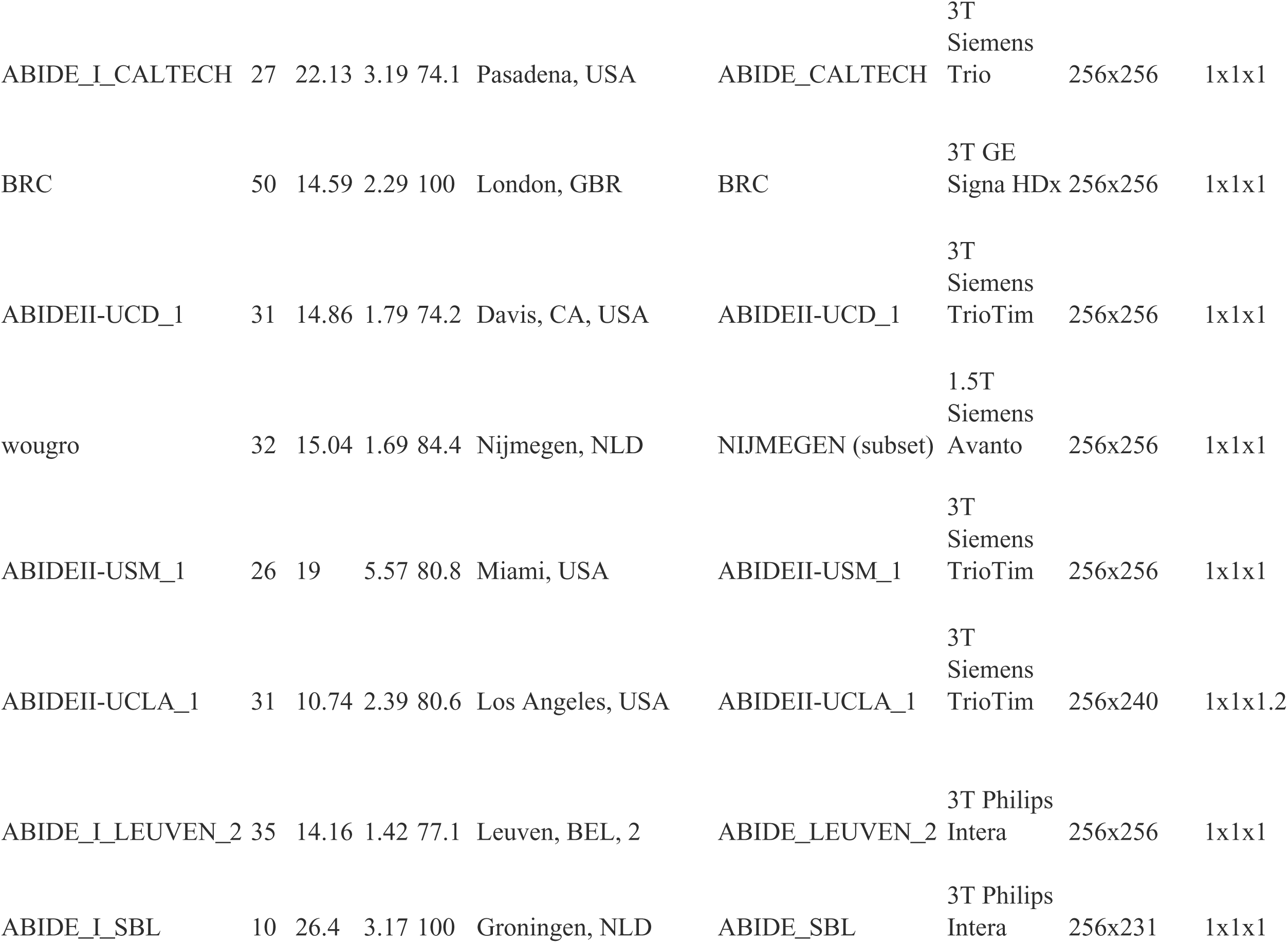
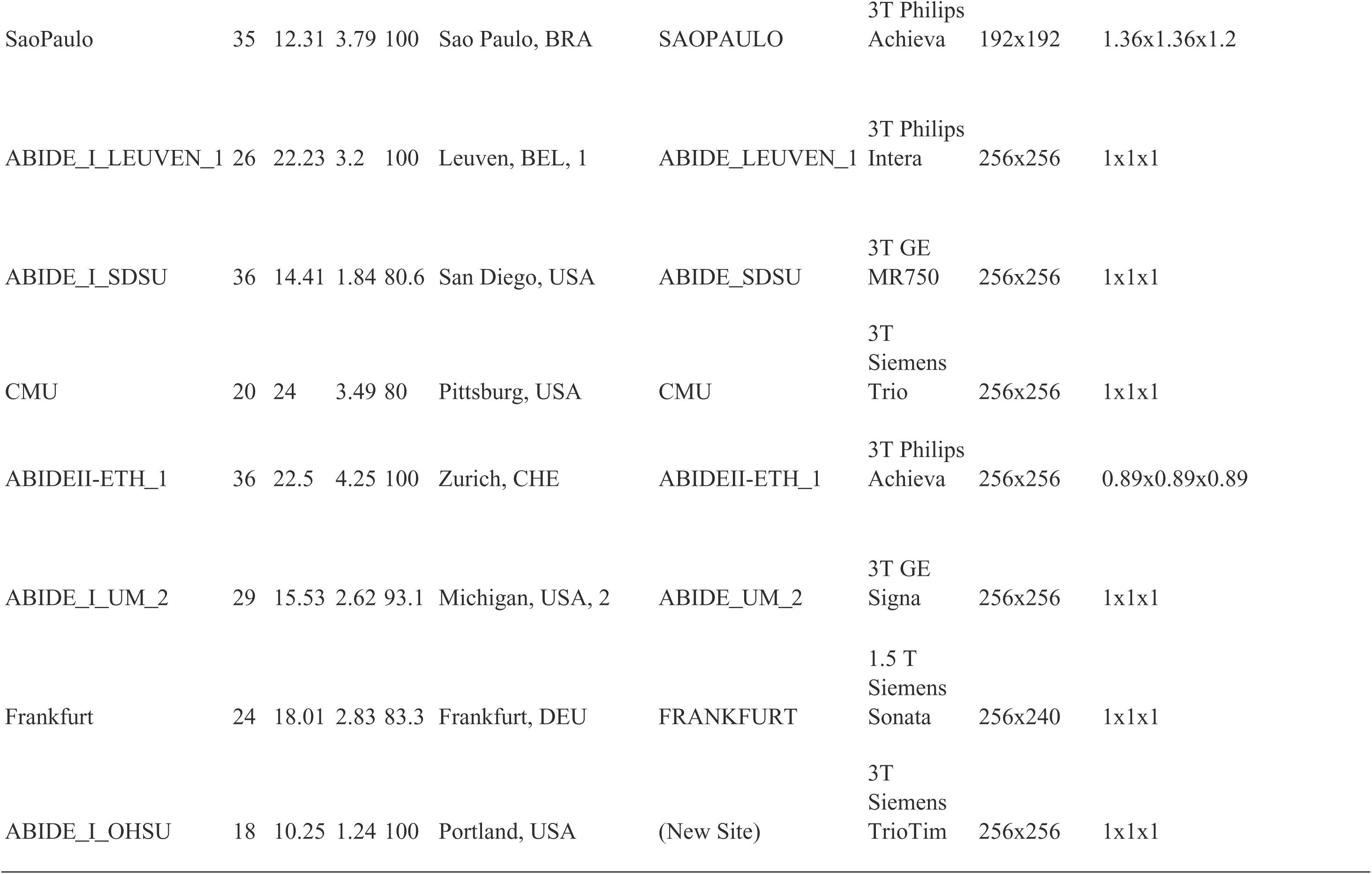
imaging acquisition details for the updated ENIGMA-ASD data.

**Supplementary Table 2.**
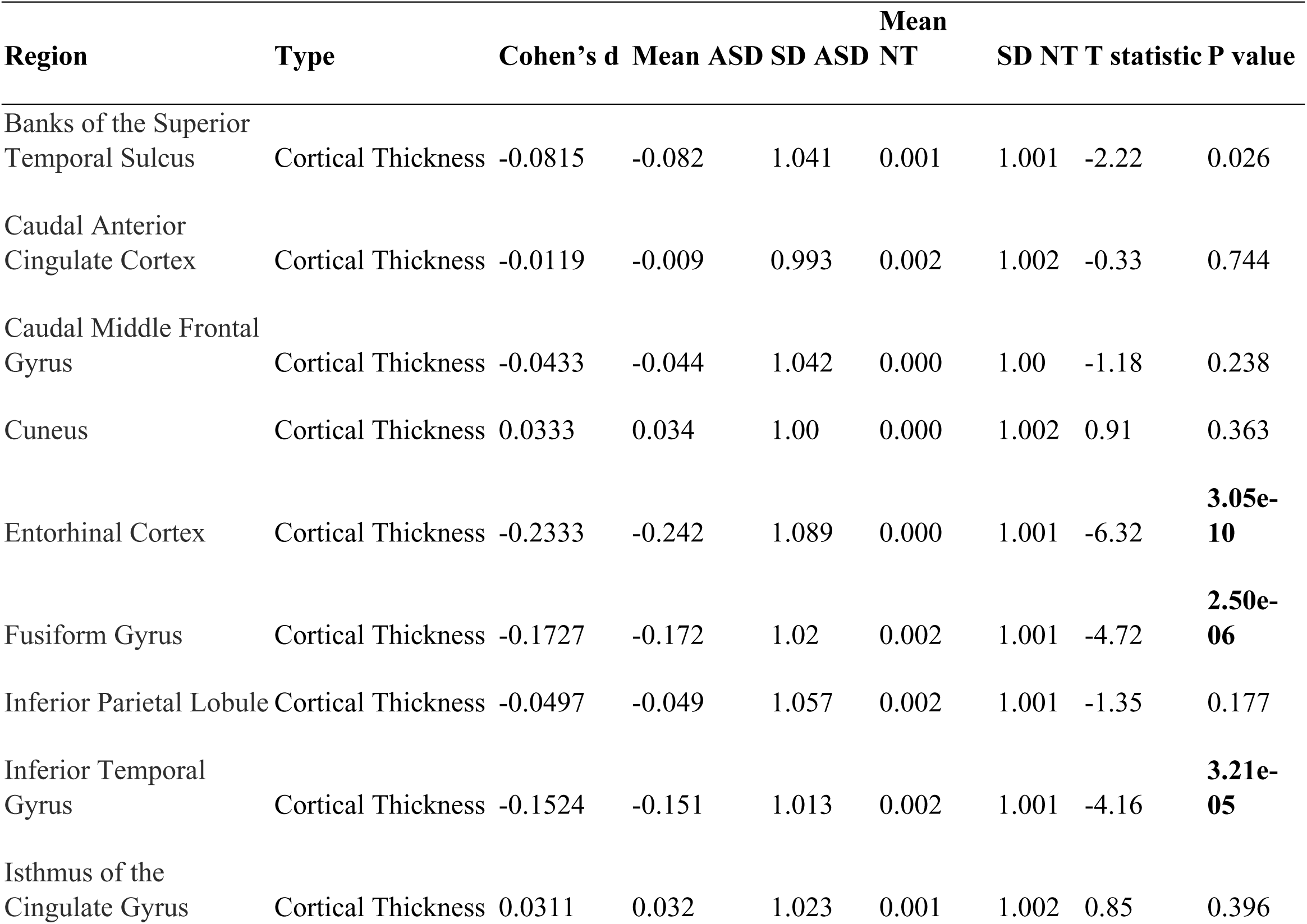
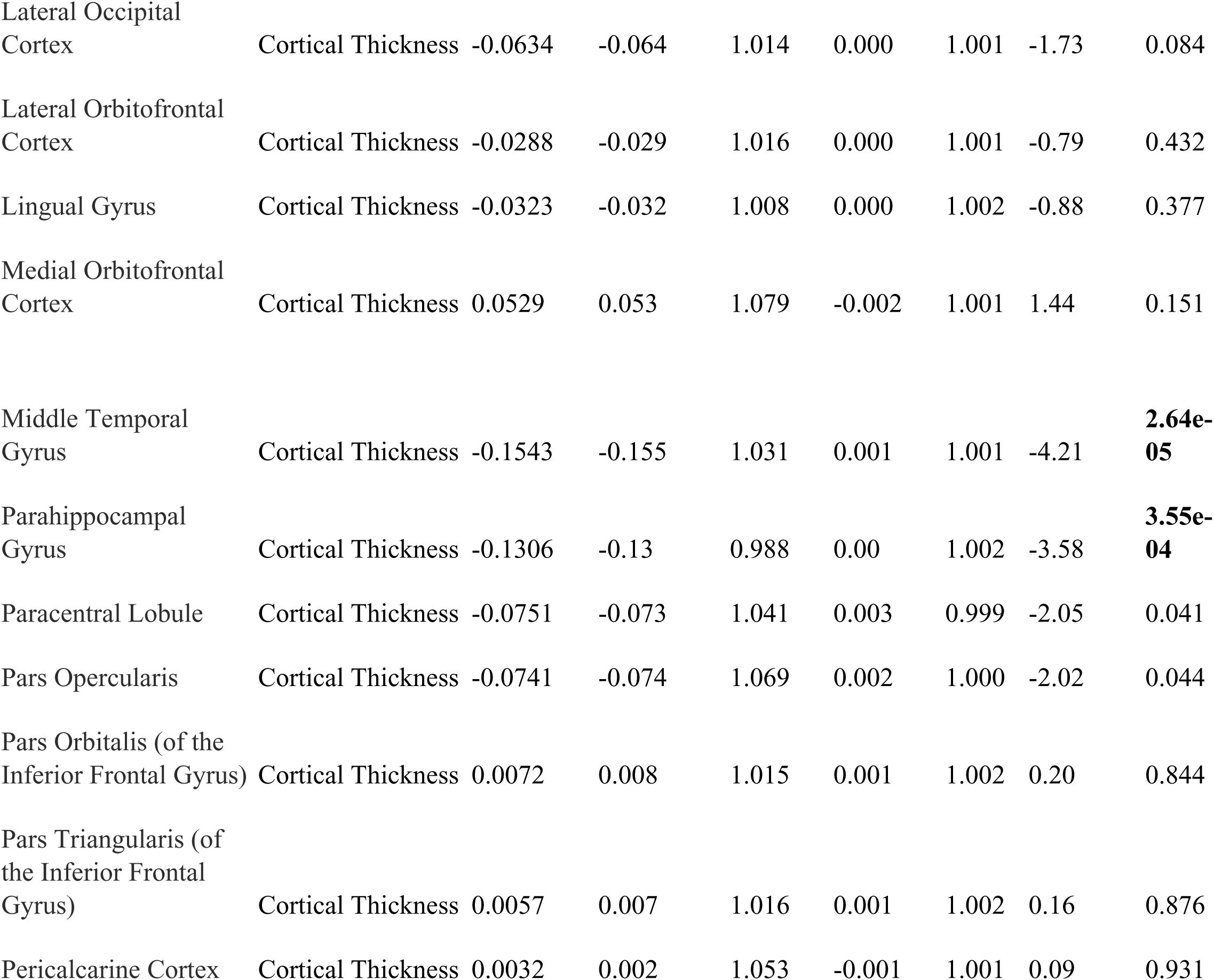
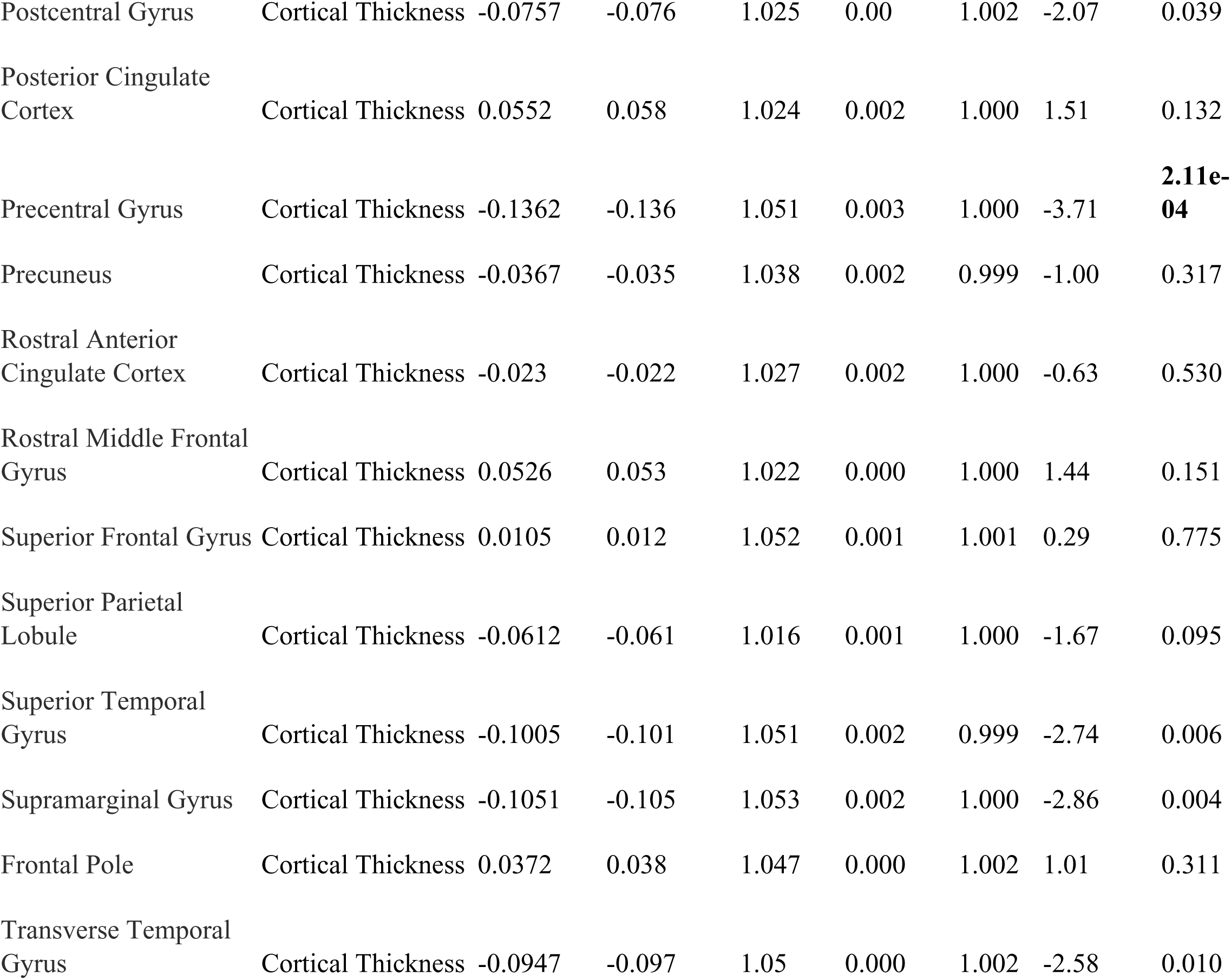
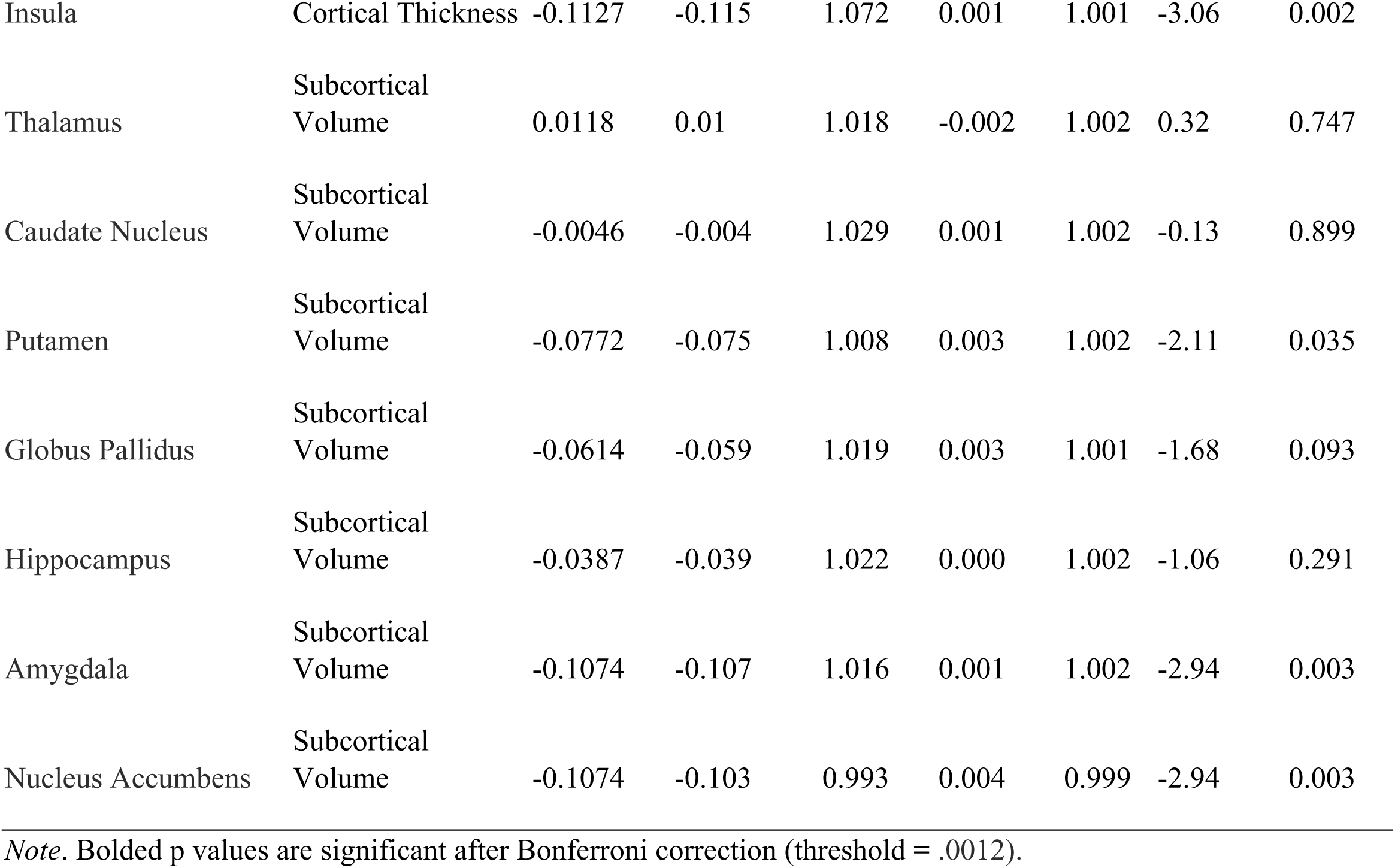
Standardized effect size of the neuroanatomical differences between individuals with autism and neurotypical controls in the update ENIGMA dataset.

**Supplementary Table 3.**
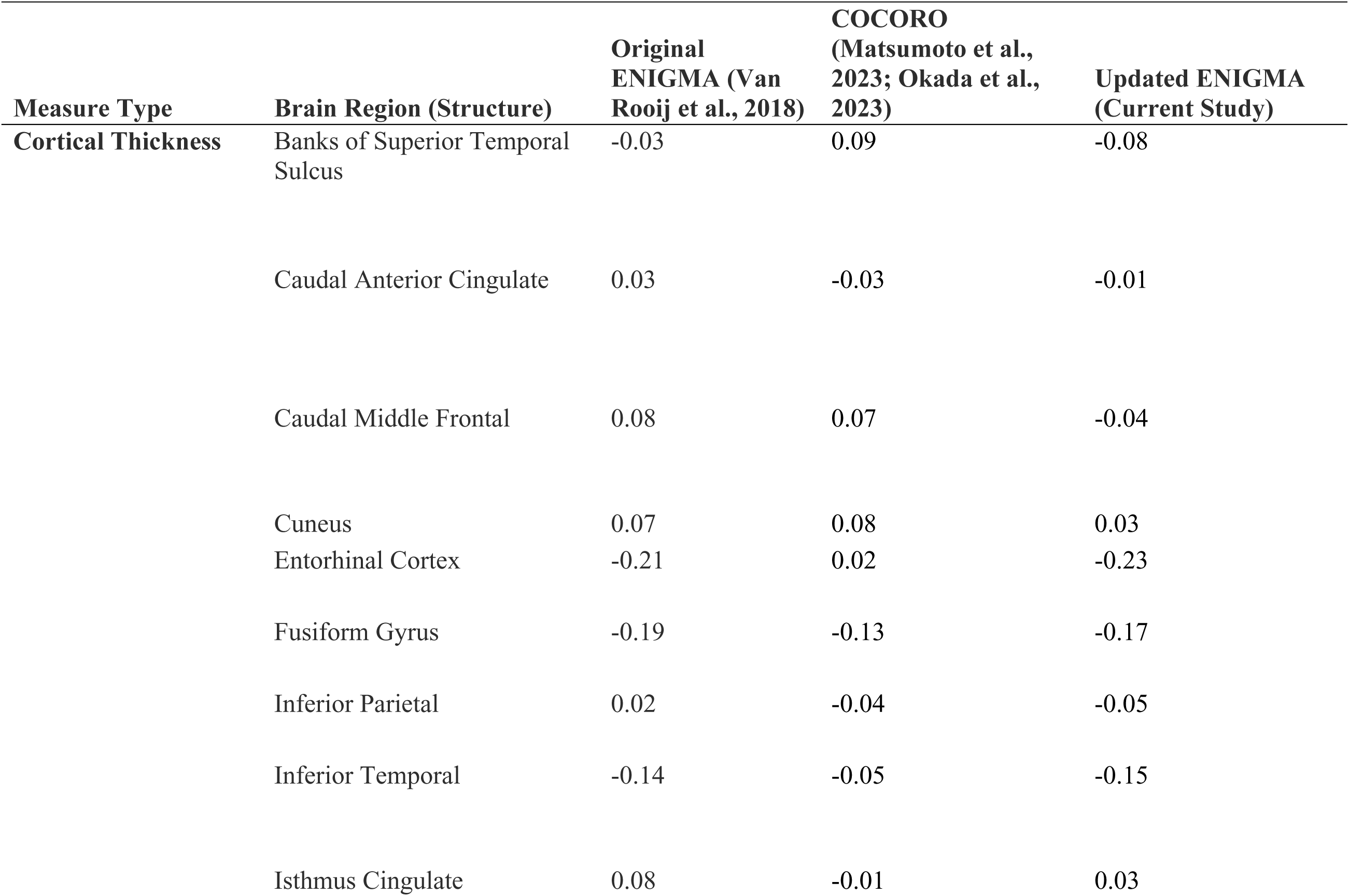
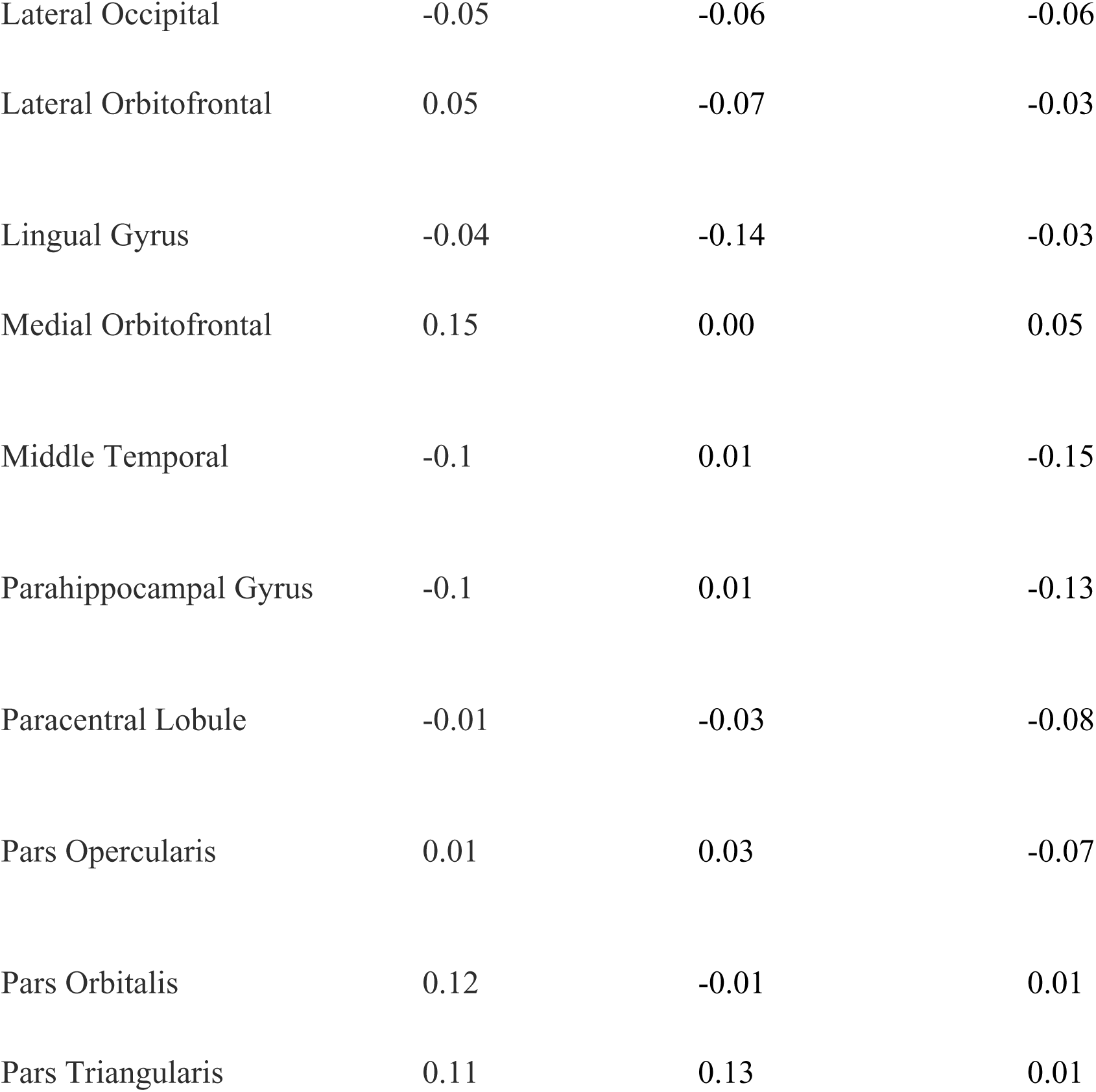
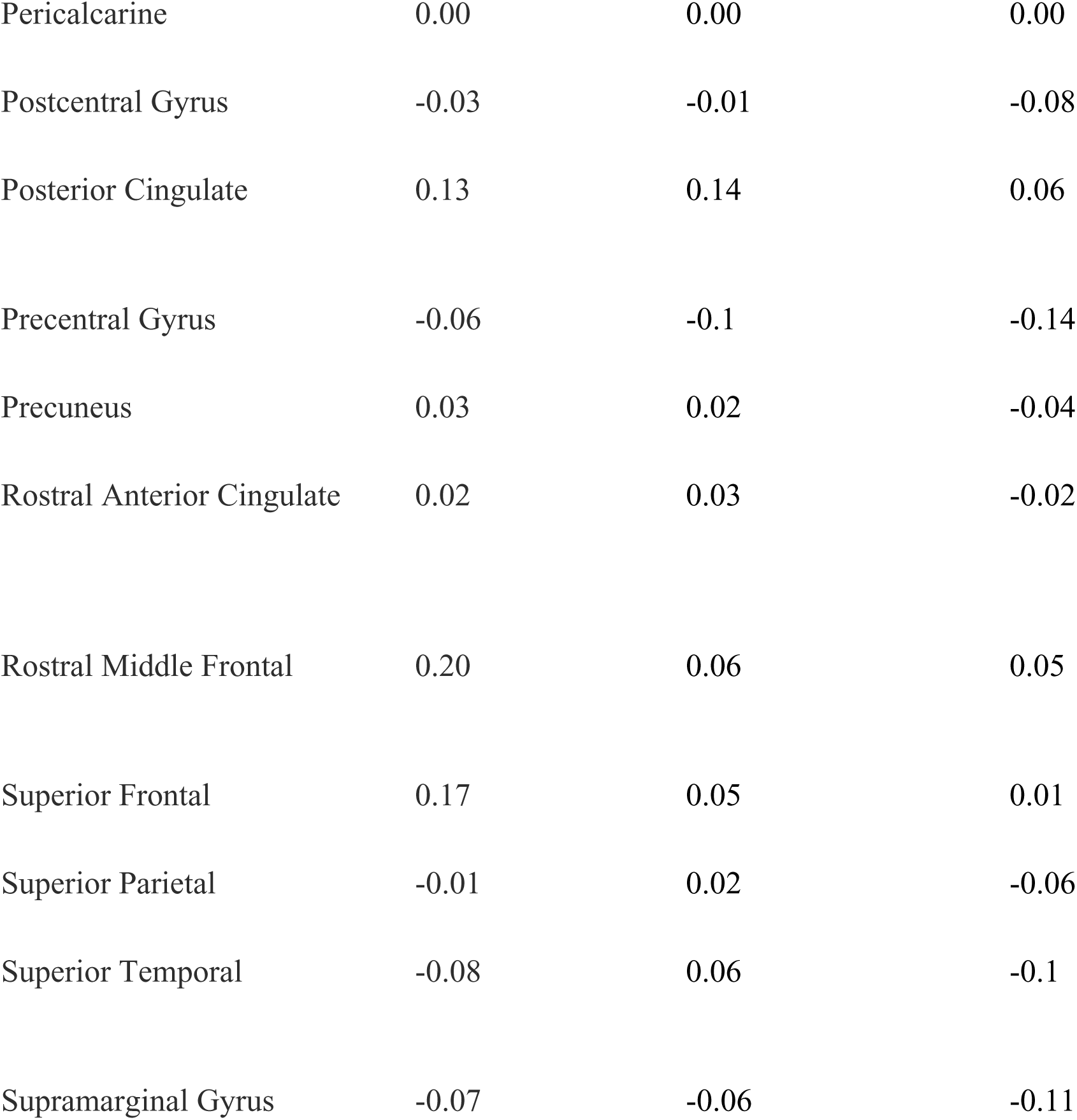
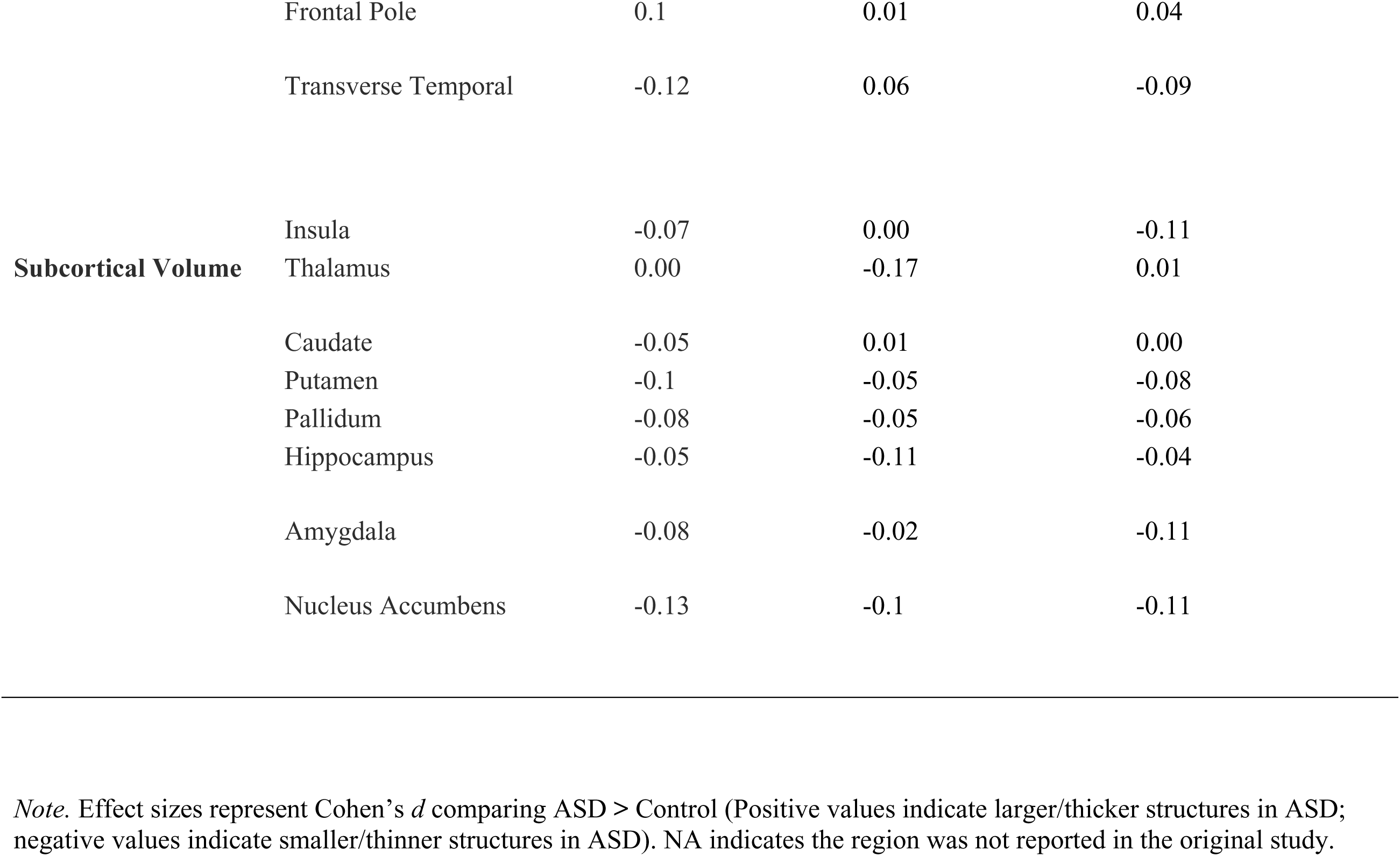
Comparison of Effect Sizes (Cohen’s *d*) Across Large-Scale Autism Neuroimaging Consortia.

**Supplementary Table 4.**
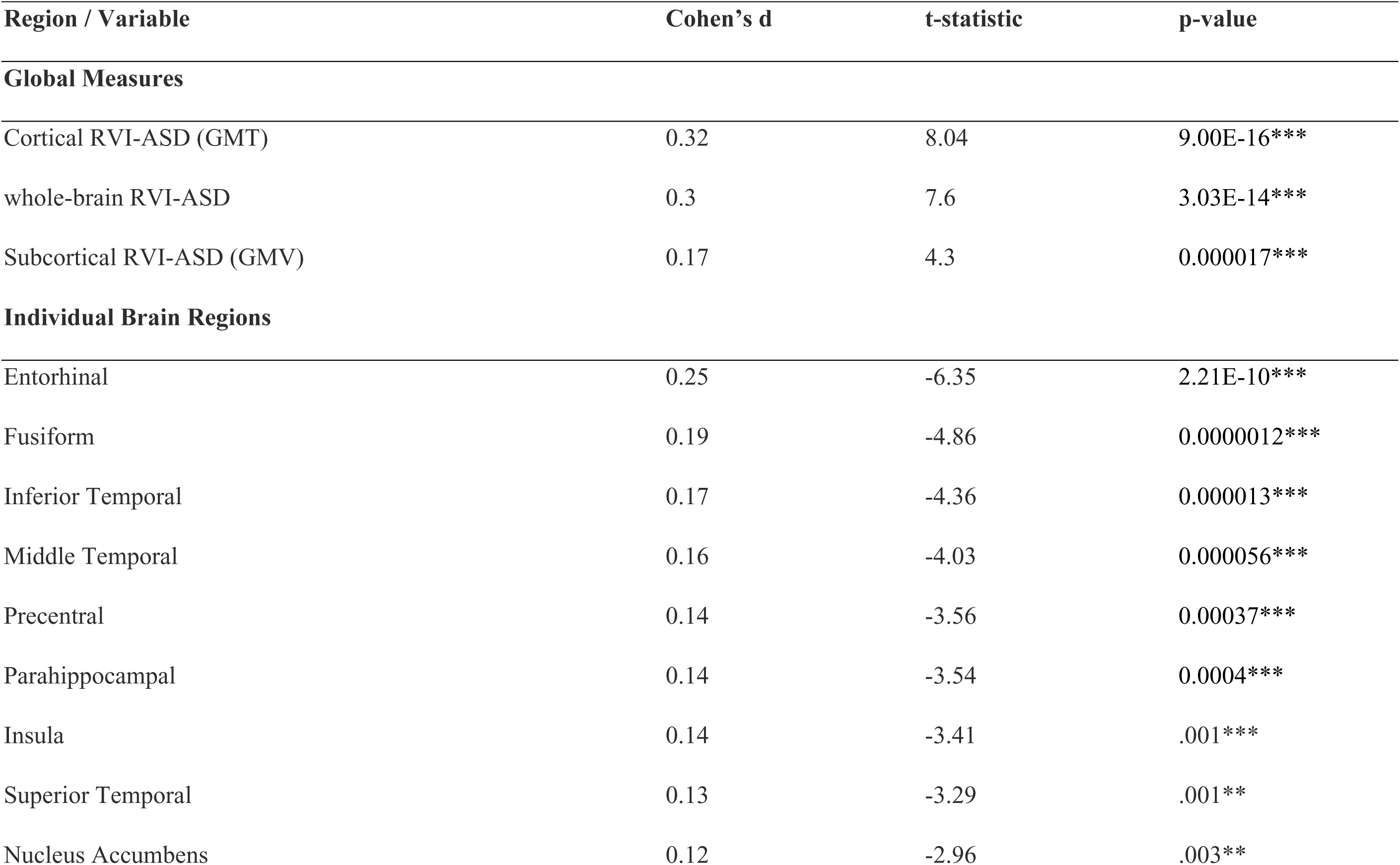
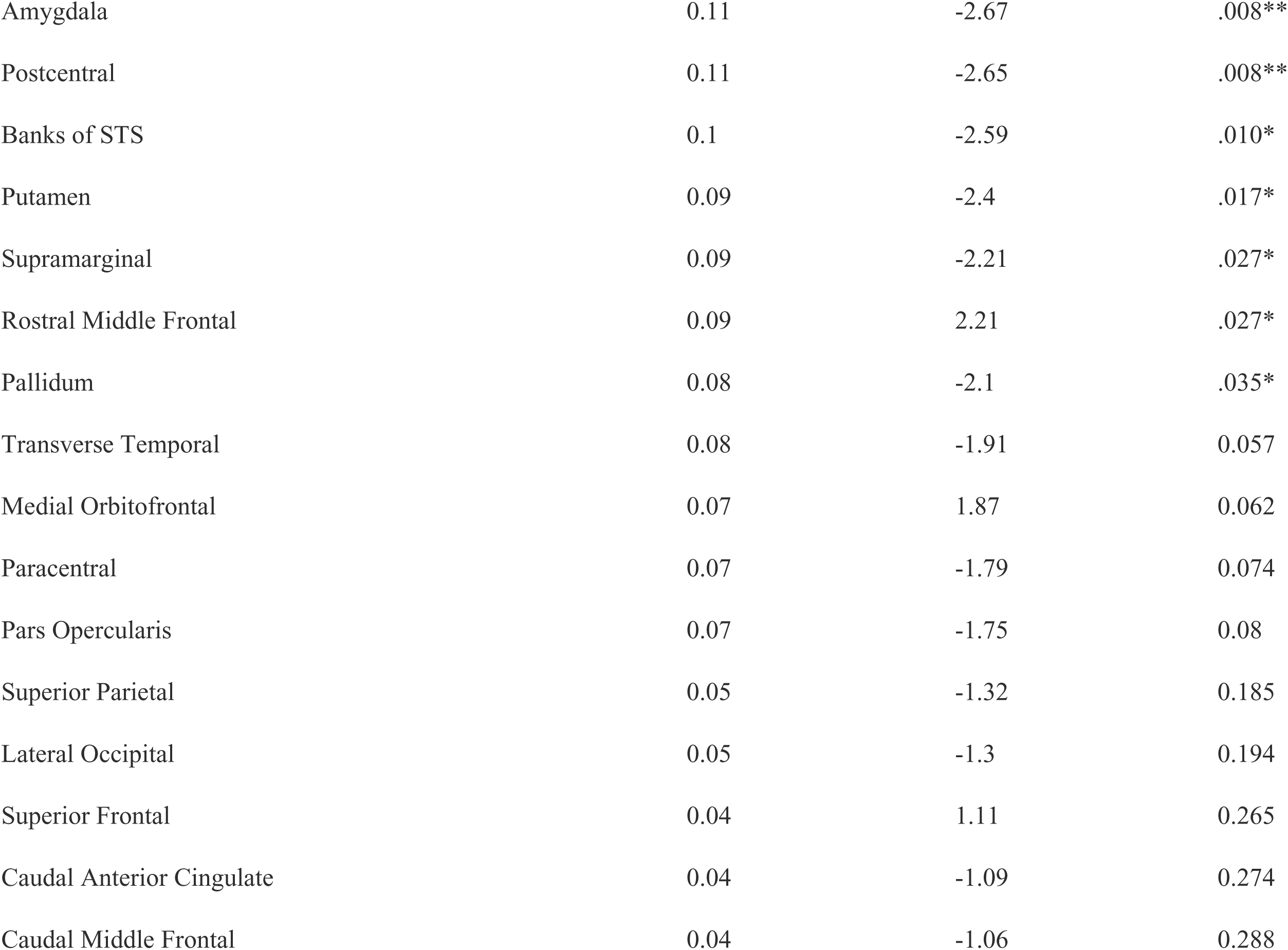
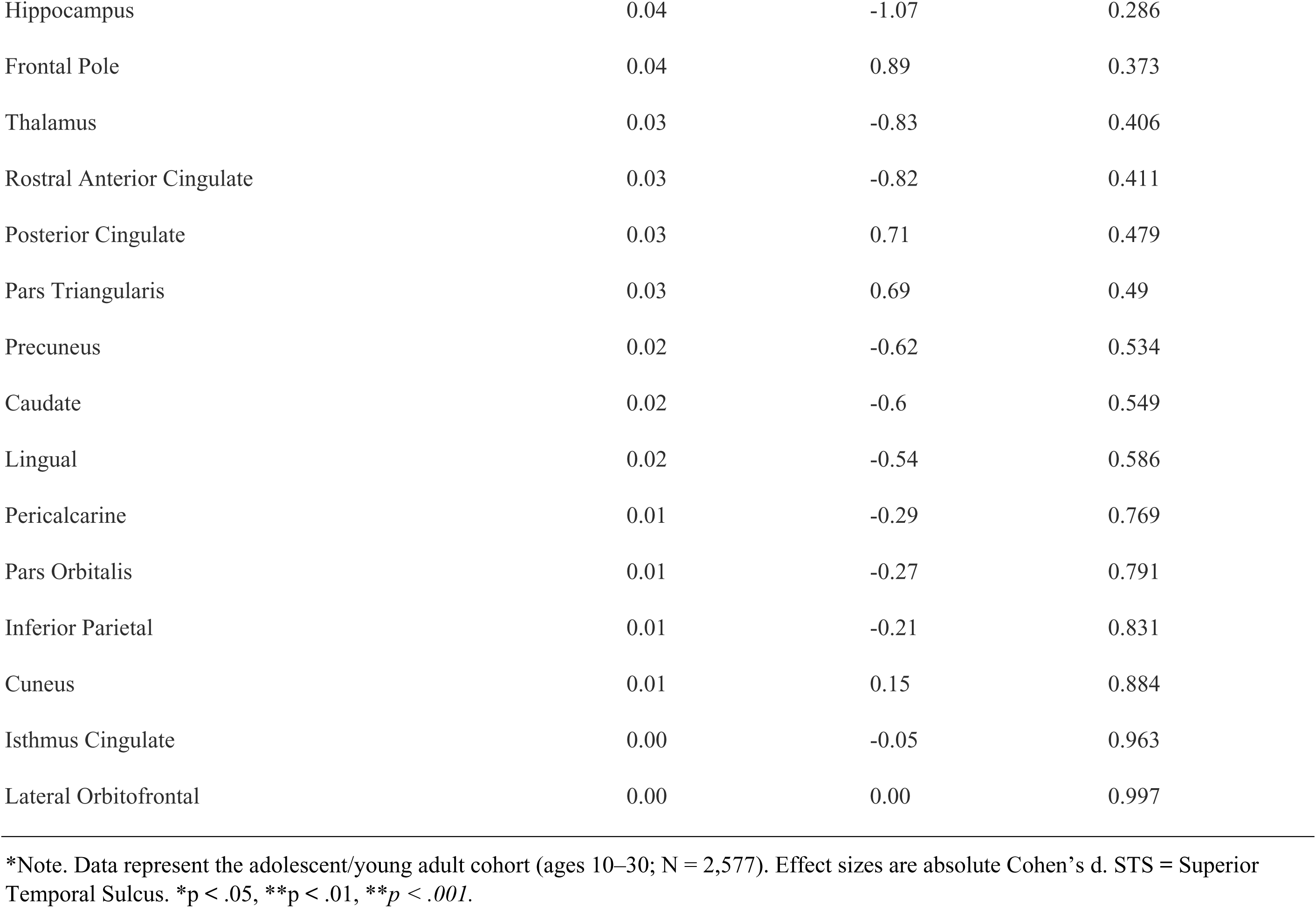
Effect Sizes for RVI and Brain Regions in Adolescents & Young Adults (Ages 10–30)

**Supplementary Table 5.** Effect Sizes for RVI and Brain Regions in Children (Under 10 years old)

| Region / Variable | Cohen's d | t-statistic | p-value |
| --- | --- | --- | --- |
| **Global Measures** |  |  |  |
| Cortical RVI-ASD (GMT) | 0.24 | 3.36 | .001** |
| whole-brain RVI-ASD | 0.20 | 2.9 | .004** |
| Subcortical RVI-ASD (GMV) | 0.11 | 1.57 | 0.116 |
| **Individual Brain Regions** |  |  |  |
| Entorhinal | 0.27 | -3.79 | 0.000162*** |
| Inferior Temporal | 0.19 | -2.64 | .008** |
| Middle Temporal | 0.18 | -2.62 | .009** |
| Fusiform | 0.18 | -2.51 | .012* |
| Precentral | 0.16 | -2.32 | .021* |
| Supramarginal | 0.15 | -2.17 | .031* |
| Parahippocampal | 0.15 | -2.09 | .037* |
| Thalamus | 0.14 | 2.04 | .042* |
| Transverse Temporal | 0.13 | -1.83 | 0.068 |
| Pericalcarine | 0.13 | 1.81 | 0.071 |
| Superior Frontal | 0.13 | -1.82 | 0.069 |
| Pars Triangularis | 0.11 | -1.59 | 0.112 |
| Pars Opercularis | 0.11 | -1.57 | 0.117 |
| Superior Temporal | 0.11 | -1.5 | 0.133 |
| Paracentral | 0.1 | -1.46 | 0.145 |
| Superior Parietal | 0.09 | -1.31 | 0.189 |
| Caudal Middle Frontal | 0.09 | -1.25 | 0.21 |
| Lateral Orbitofrontal | 0.09 | -1.23 | 0.217 |
| Pallidum | 0.08 | 1.17 | 0.241 |
| Nucleus Accumbens | 0.08 | -1.16 | 0.247 |
| Amygdala | 0.08 | -1.09 | 0.276 |
| Putamen | 0.07 | 0.99 | 0.324 |
| Insula | 0.07 | -0.94 | 0.348 |
| Cuneus | 0.06 | 0.91 | 0.366 |
| Caudate | 0.06 | 0.9 | 0.368 |
| Pars Orbitalis | 0.06 | -0.89 | 0.376 |
| Banks of STS | 0.06 | -0.85 | 0.396 |
| Inferior Parietal | 0.06 | -0.84 | 0.403 |
| Isthmus Cingulate | 0.06 | 0.8 | 0.423 |
| Rostral Anterior Cingulate | 0.05 | -0.75 | 0.451 |
| Precuneus | 0.05 | -0.7 | 0.485 |
| Lateral Occipital | 0.04 | -0.63 | 0.529 |
| Postcentral | 0.04 | -0.5 | 0.619 |
| Posterior Cingulate | 0.03 | 0.49 | 0.622 |
| Hippocampus | 0.03 | -0.48 | 0.63 |
| Lingual | 0.03 | -0.43 | 0.668 |
| Medial Orbitofrontal | 0.02 | 0.35 | 0.729 |
| Rostral Middle Frontal | 0.02 | -0.31 | 0.759 |
| Caudal Anterior Cingulate | 0.01 | 0.18 | 0.857 |
| Frontal Pole | 0 | -0.04 | 0.972 |

\*Note. Data represent the childhood cohort (under 10 years old; $N = 812$ ). Effect sizes are absolute Cohen's $d$ . STS = Superior Temporal Sulcus. \* $p < .05$ , \*\* $p < .01$ , \*\*\* $p < .001$ .

**Supplementary Table 6.** Correlations between Risk Factors and RVI Measures.

| Measure | RVI Baseline<br>(r) | RVI Baseline<br>(p) | RVI Second<br>(r) | RVI Second<br>(p) |
| --- | --- | --- | --- | --- |
| **Baseline Measures** |  |  |  |  |
| Pre-Perinatal | 0.022 | 0.144 | 0.008 | 0.345 |
| C-Section | 0.000 | 0.399 | -0.015 | 0.247 |
| Pregnancy Complications | 0.029 | 0.064 | 0.019 | 0.188 |
| Birth Complications | 0.020 | 0.171 | 0.007 | 0.364 |
| Pre-Perinatal (Composite) | 0.006 | 0.37 | -0.001 | 0.398 |
| Months Breastfed | 0.024 | 0.113 | 0.014 | 0.26 |
| PM2.5 Average | 0.001 | 0.397 | 0.001 | 0.397 |
| Lead Risk | -0.002 | 0.396 | -0.007 | 0.359 |
| Income-to-Needs | 0.017 | 0.216 | 0.025 | 0.108 |
| Financial Difficulty | -.037* | 0.021 | -0.031 | 0.056 |
| Neighbor Safety (P) | 0.015 | 0.246 | 0.018 | 0.208 |
| Neighbor Safety (Y) | 0.011 | 0.303 | 0.016 | 0.238 |
| Best Friend | 0.004 | 0.388 | 0.003 | 0.391 |
| Regular Friend | -0.018 | 0.202 | -0.004 | 0.389 |
| Bullying | -0.003 | 0.391 | -0.011 | 0.312 |
| NLE (P) | 0.001 | 0.398 | 0.004 | 0.388 |
| NLE (Y) | -0.01 | 0.323 | -0.027 | 0.089 |
| Active Days | 0.011 | 0.309 | 0.008 | 0.354 |
| Screen Use | -0.006 | 0.37 | 0.00 | 0.399 |
| Sleep Disturbance Total | -0.011 | 0.311 | -0.013 | 0.283 |
| Family Conflict (P) | -0.023 | 0.133 | -0.026 | 0.101 |
| Family Conflict (Y) | 0.002 | 0.395 | 0.002 | 0.395 |
| School Environment | 0.002 | 0.394 | 0.001 | 0.397 |
| School Pos. Involvement | -0.013 | 0.277 | -0.01 | 0.326 |
| School Disengagement | 0.009 | 0.335 | 0.012 | 0.3 |
| CBCL Anx/Dep | 0.006 | 0.373 | 0.001 | 0.398 |
| CBCL Withdrawn/Dep | -0.013 | 0.273 | -0.015 | 0.256 |
| CBCL Social Problems | 0.004 | 0.384 | 0.007 | 0.36 |
| CBCL DSM5 Depression | 0.008 | 0.348 | -0.001 | 0.398 |
| CBCL DSM5 Anxiety | 0.009 | 0.331 | 0.004 | 0.385 |
| CBCL OCD | 0.006 | 0.370 | 0.003 | 0.393 |
| CBCL DSM5 ADHD | -0.011 | 0.307 | -0.013 | 0.276 |
| PGBI Score | -0.01 | 0.321 | -0.001 | 0.398 |
| **2-Year Measures** |  |  |  |  |
| Neighborhood Crime | 0.023 | 0.132 | 0.019 | 0.192 |
| PM2.5 Average | 0.001 | 0.397 | 0.001 | 0.397 |
| Lead Risk | -0.002 | 0.396 | -0.007 | 0.359 |
| Income-to-Needs | 0.029 | 0.071 | .033* | 0.043 |
| Financial Difficulty | -.041* | 0.011 | -.035* | 0.028 |
| Neighbor Safety (P) | 0.016 | 0.239 | 0.022 | 0.151 |
| Neighbor Safety (Y) | 0.015 | 0.25 | 0.008 | 0.346 |
| Best Friend | 0.003 | 0.391 | 0.011 | 0.304 |
| Regular Friend | 0.008 | 0.349 | 0.024 | 0.117 |
| Bullying | 0.002 | 0.397 | -0.002 | 0.397 |
| NLE (P) | -0.026 | 0.092 | -0.017 | 0.214 |
| NLE (Y) | 0.012 | 0.299 | 0.017 | 0.218 |
| Active Days | 0.00 | 0.399 | 0.006 | 0.368 |
| Screen Use | -0.024 | 0.123 | -0.016 | 0.232 |
| Sleep Disturbance Total | 0.00 | 0.399 | -0.002 | 0.397 |
| Family Conflict (P) | -0.006 | 0.373 | -0.006 | 0.37 |
| Family Conflict (Y) | -0.025 | 0.112 | -0.025 | 0.11 |
| School Environment | 0.001 | 0.398 | -0.01 | 0.327 |
| School Pos. Involvement | -0.009 | 0.331 | -0.015 | 0.251 |
| School Disengagement | -0.001 | 0.398 | 0.01 | 0.33 |
| CBCL Anx/Dep | 0.03 | 0.058 | .036* | 0.027 |
| CBCL Withdrawn/Dep | 0.008 | 0.351 | 0.014 | 0.272 |
| CBCL Social Problems | 0.007 | 0.362 | 0.002 | 0.396 |
| CBCL DSM5 Depression | 0.018 | 0.206 | 0.021 | 0.157 |
| CBCL DSM5 Anxiety | 0.022 | 0.142 | 0.028 | 0.075 |
| CBCL OCD | 0.022 | 0.142 | 0.024 | 0.118 |
| CBCL DSM5 ADHD | -0.019 | 0.192 | -0.019 | 0.188 |
| PGBI Score | -0.009 | 0.336 | -0.002 | 0.395 |
| **Polygenic & Clinical Scores** |  |  |  |  |
| MDD_L | -.034* | 0.037 | -.032* | 0.049 |
| BD_L | 0.013 | 0.279 | 0.027 | 0.082 |
| SSD_L | -0.014 | 0.265 | -0.004 | 0.387 |
| BDPRS | -0.009 | 0.342 | 0.002 | 0.394 |
| SSDPRS | -0.002 | 0.394 | 0.00 | 0.399 |
| MDDPRS | 0.00 | 0.399 | -0.016 | 0.226 |
| **Social Responsiveness Scale (SRS)** |  |  |  |  |
| SRS Difficulty with Routine (ssrs_24) | -0.021 | 0.188 | -0.015 | 0.35 |
| SRS Unusual Communication (ssrs_39) | -0.017 | 0.29 | -0.02 | 0.193 |
| SRS Social Cues (ssrs_18) | -0.018 | 0.254 | -0.018 | 0.242 |
| SRS Conversation Flow (ssrs_35) | -0.016 | 0.308 | -0.017 | 0.285 |
| SRS Difficulty Making Friends (ssrs_37) | -0.016 | 0.318 | -0.019 | 0.225 |
| SRS Social Withdrawal (ssrs_6) | -0.014 | 0.377 | -0.021 | 0.179 |

**Supplementary Table 7.**
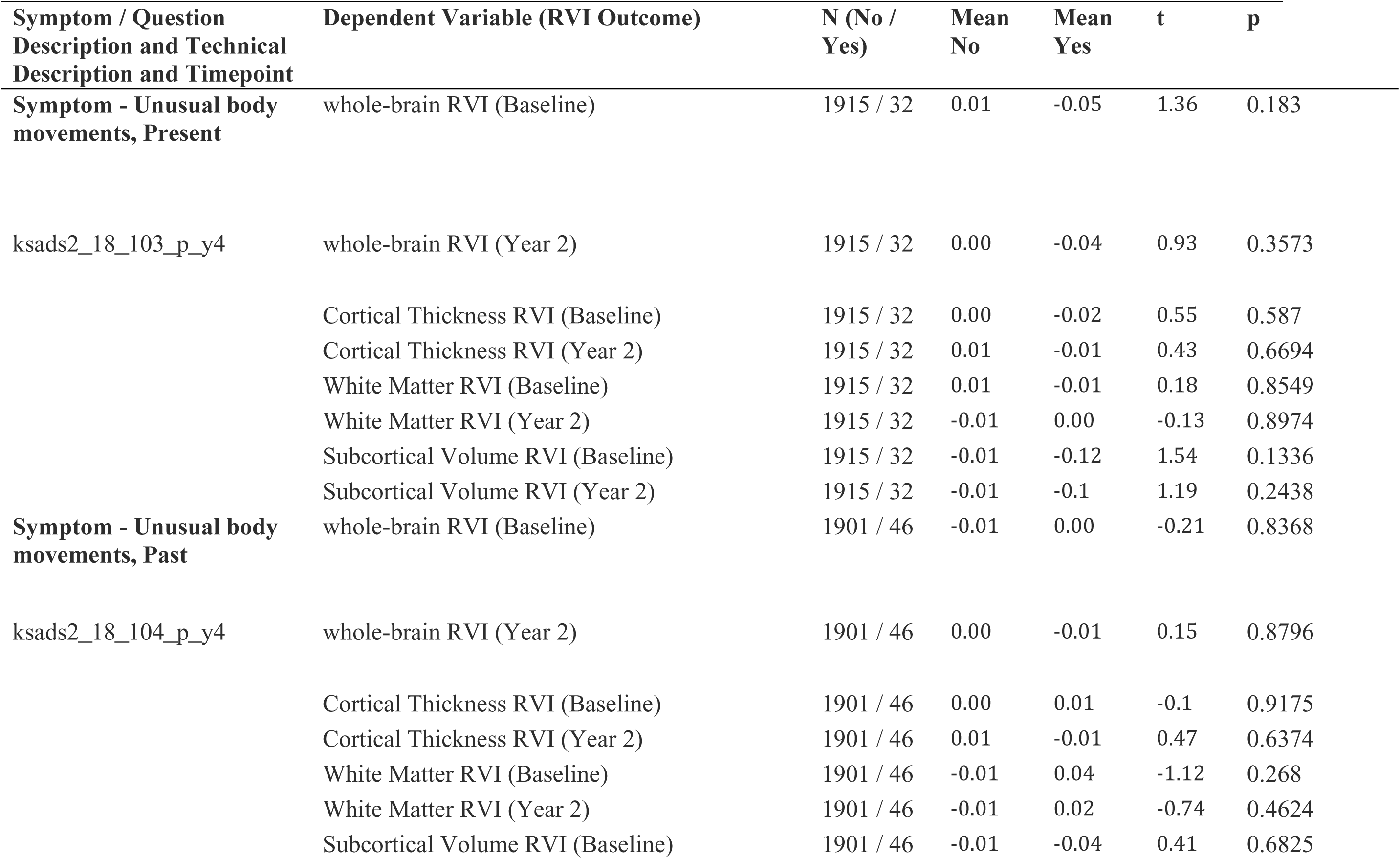
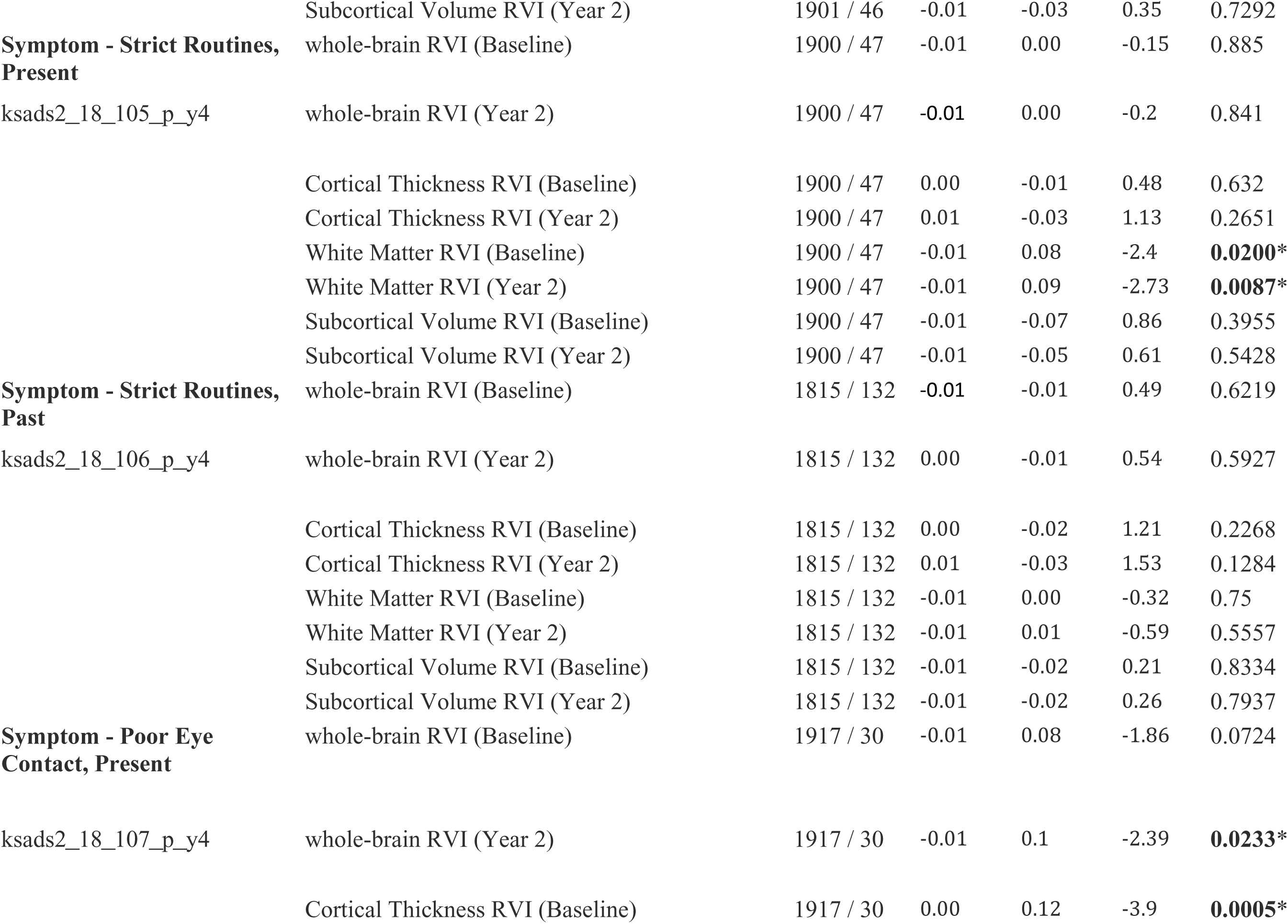
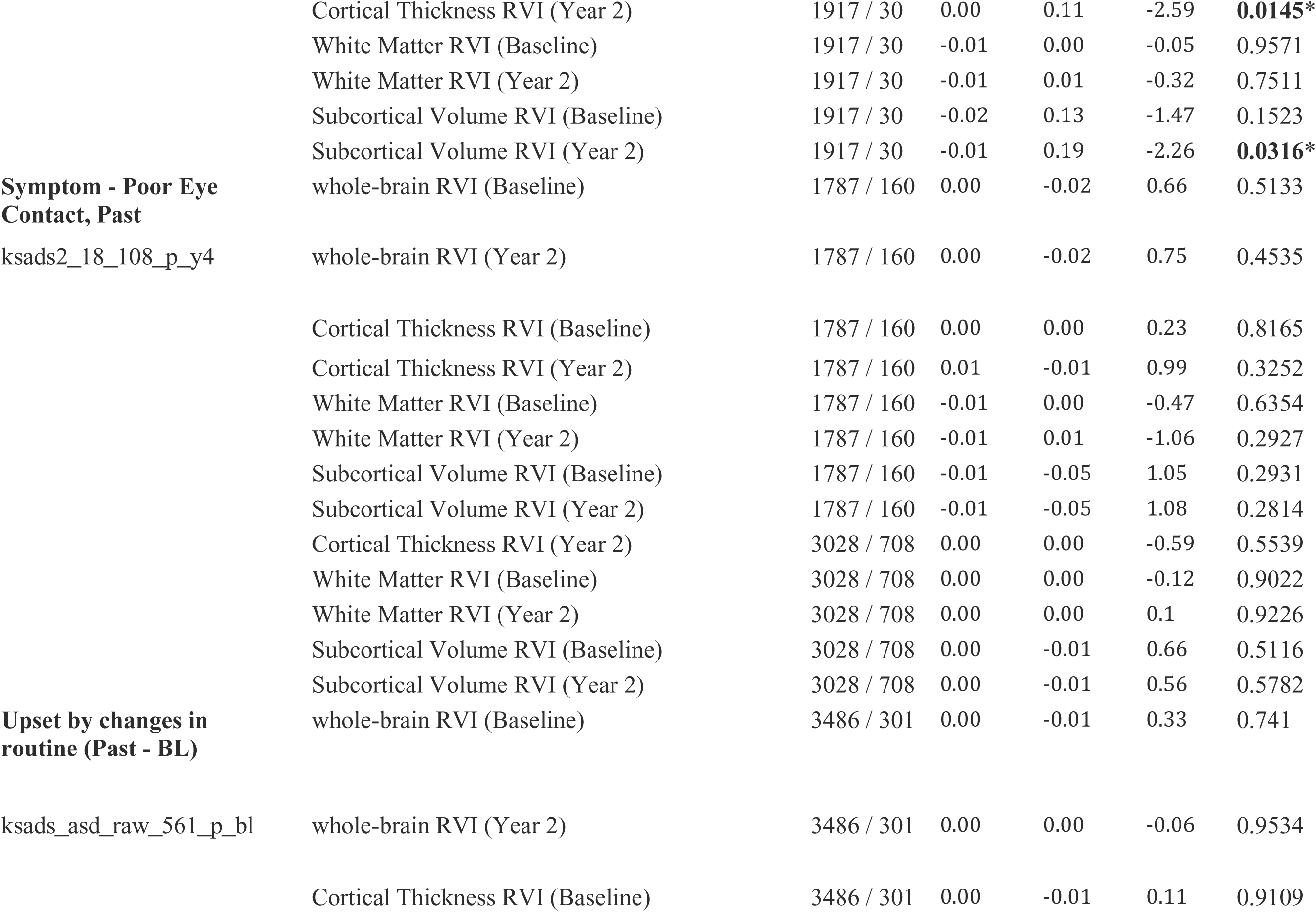
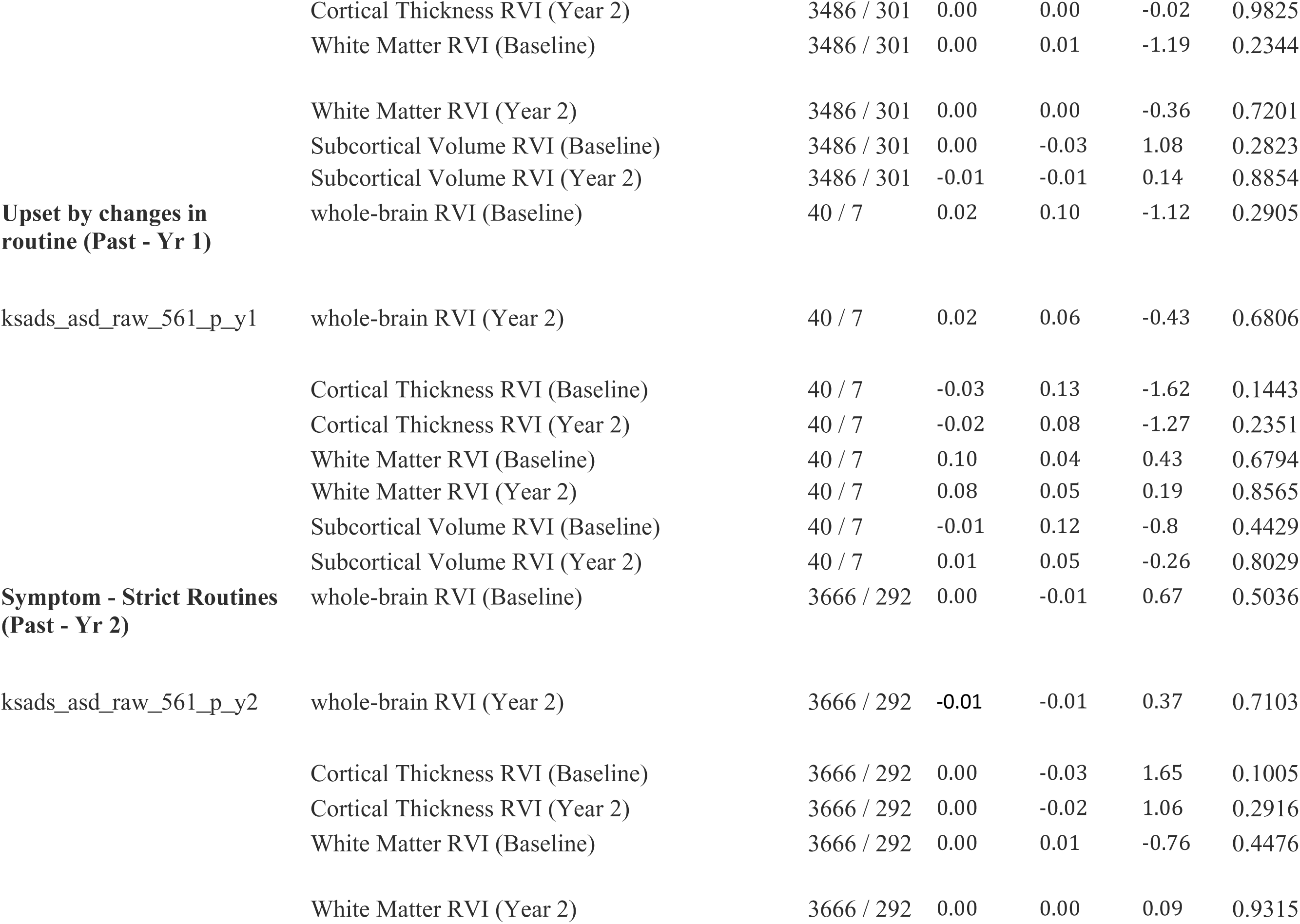
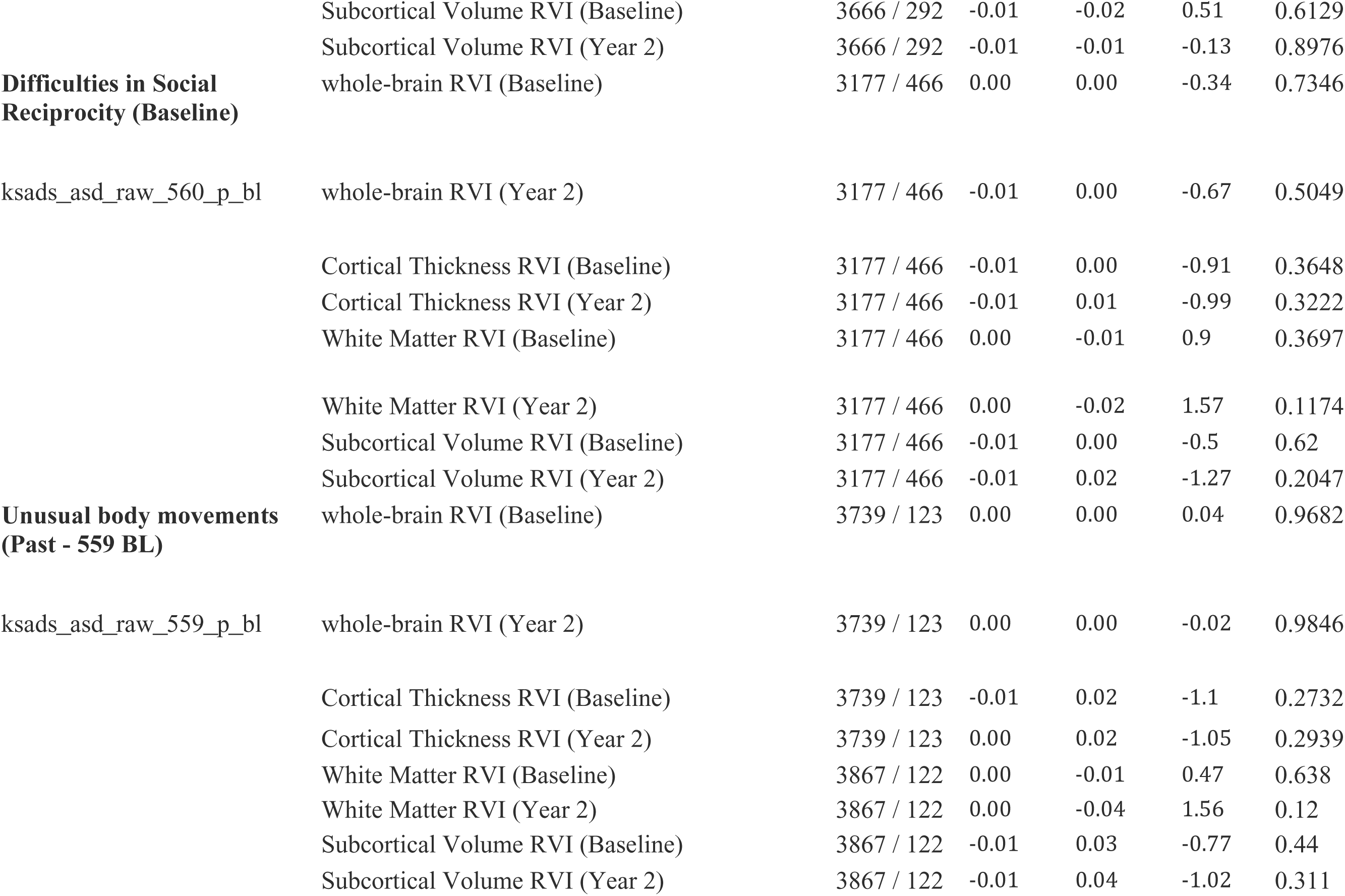
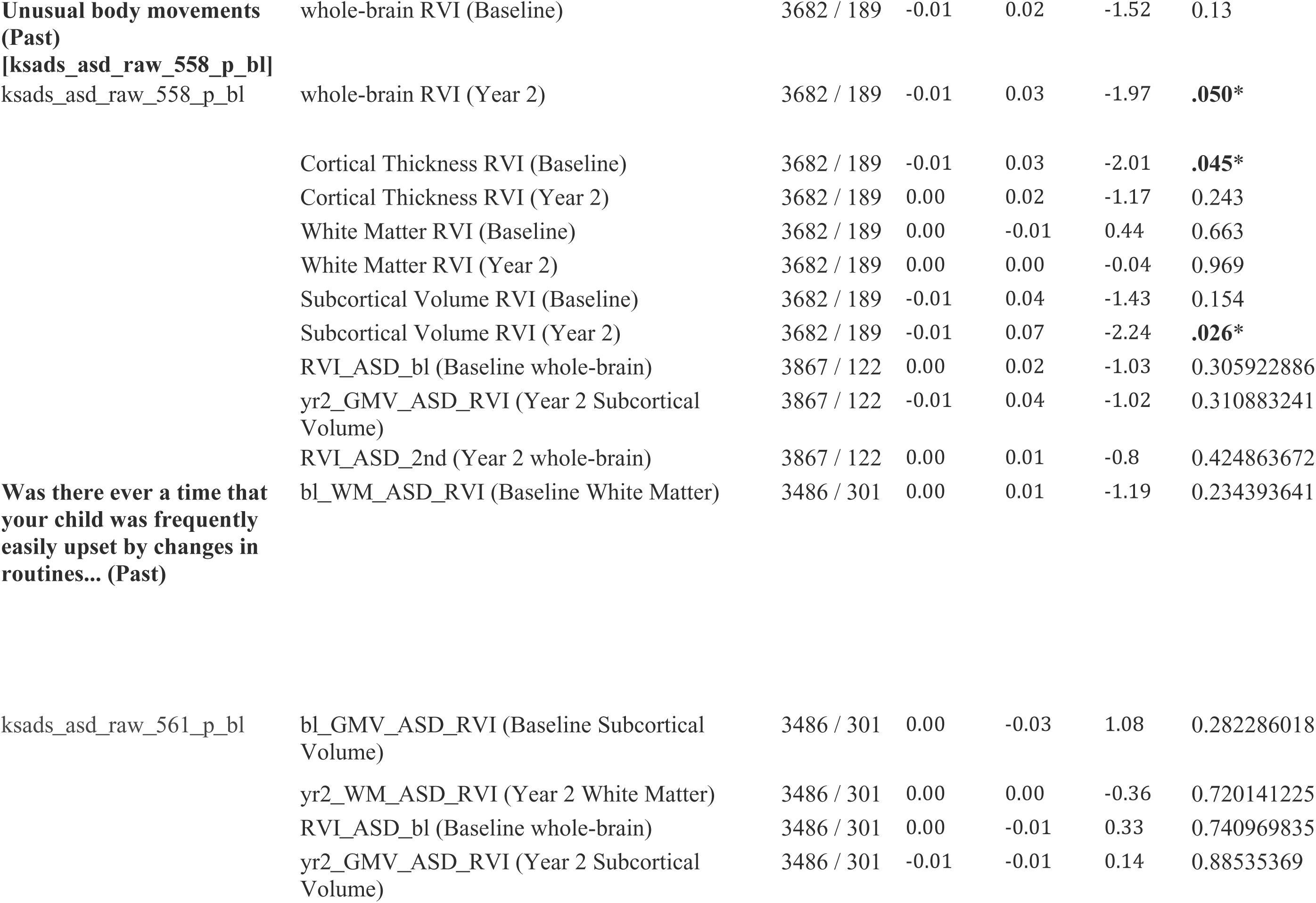
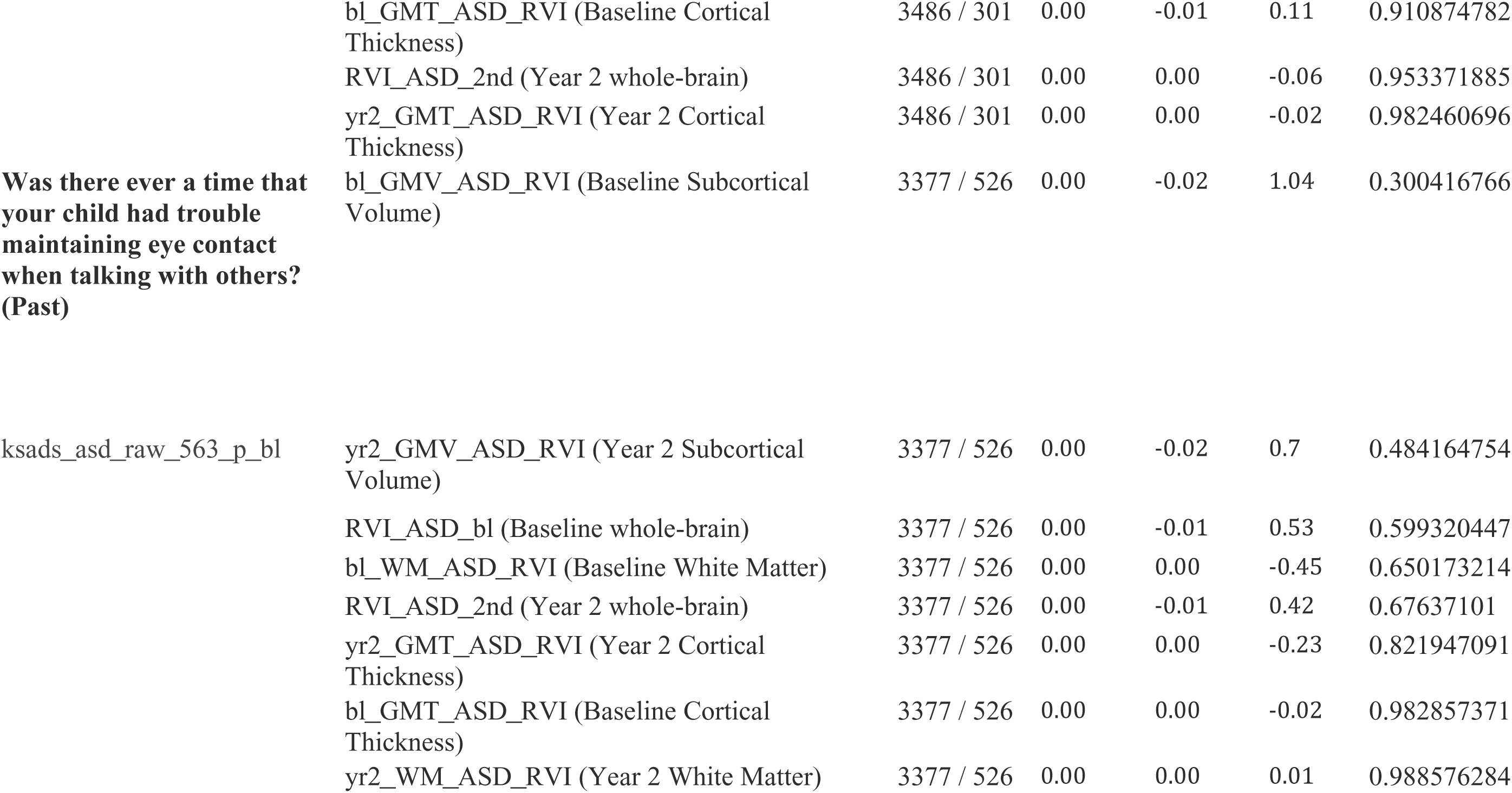
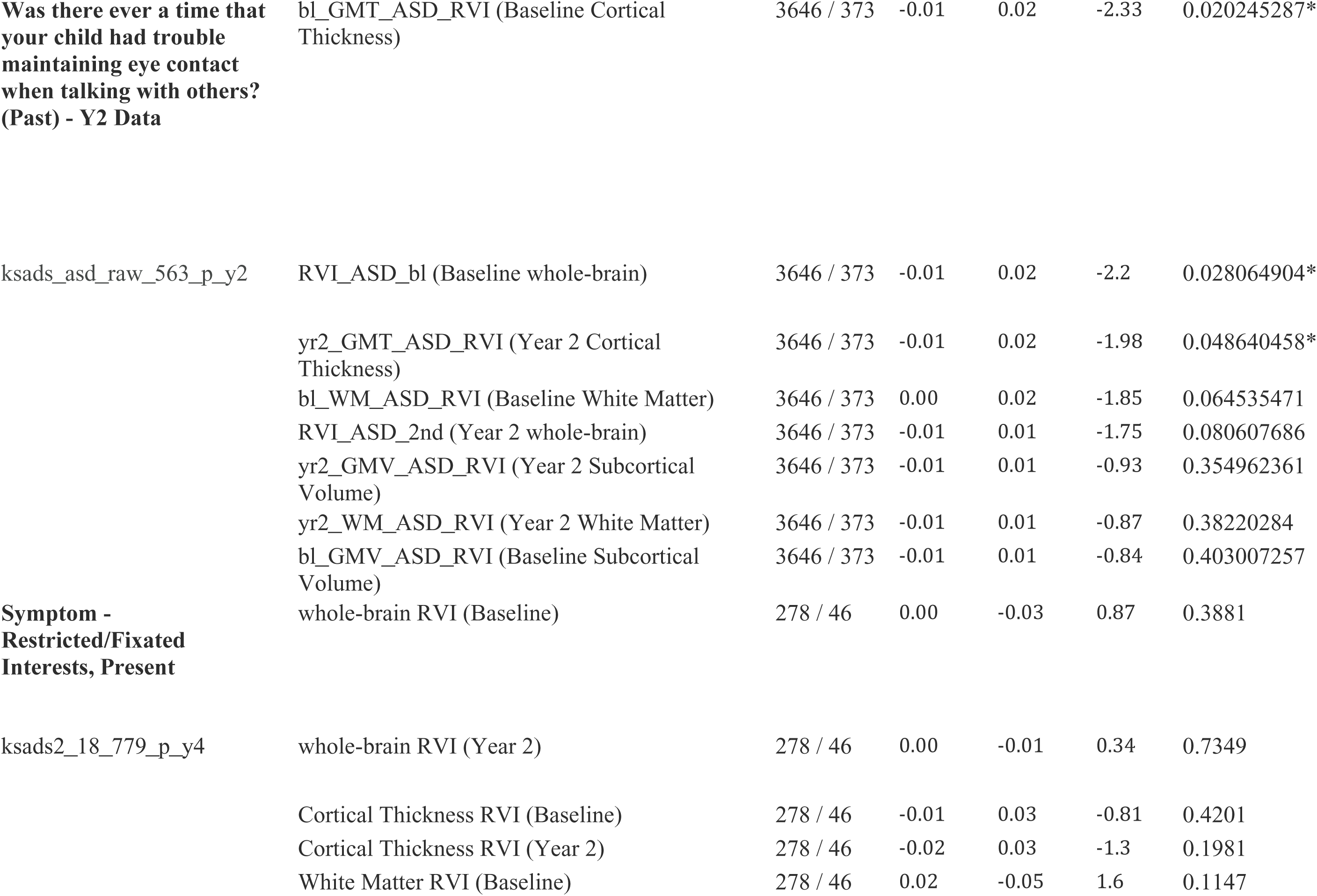
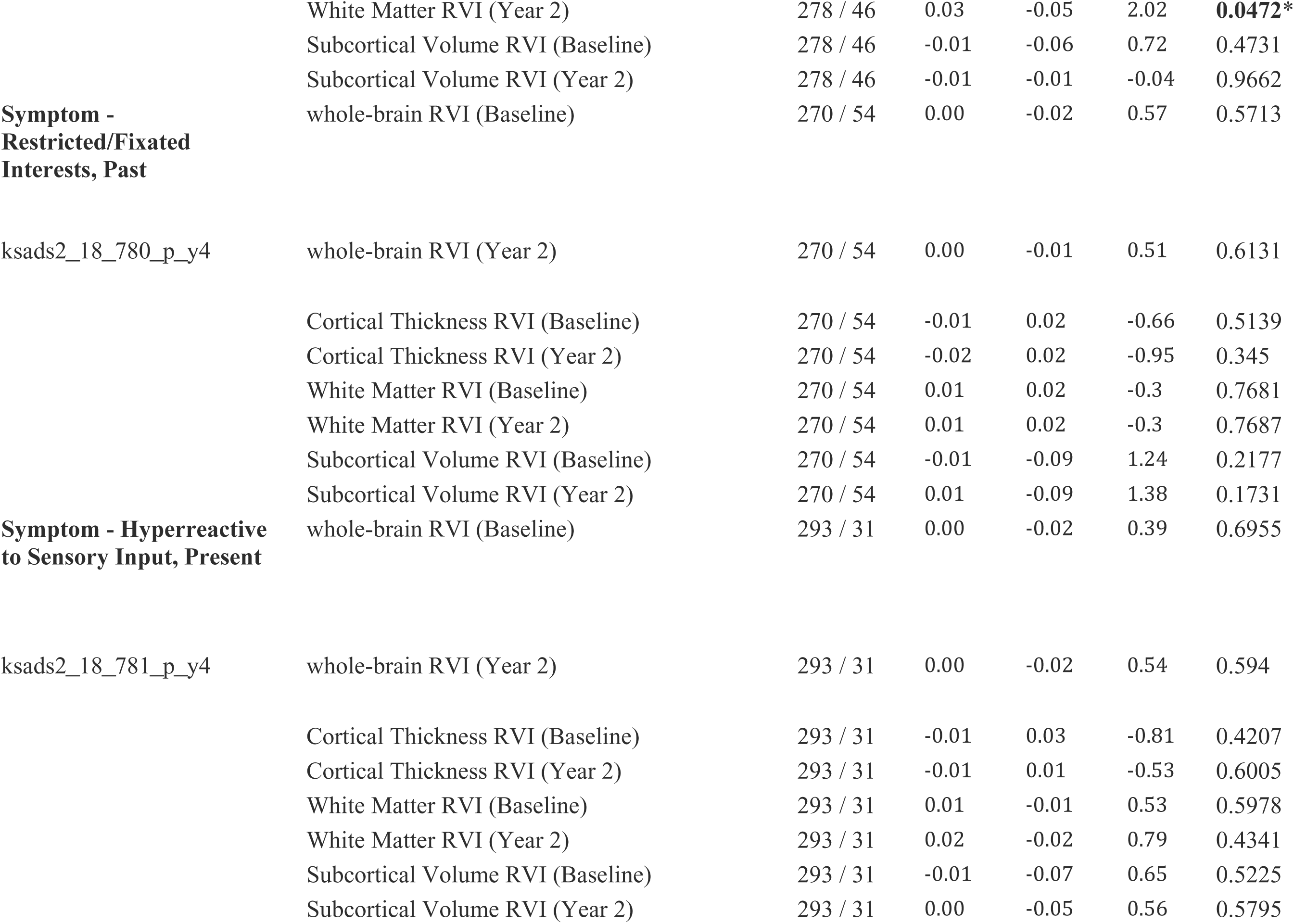
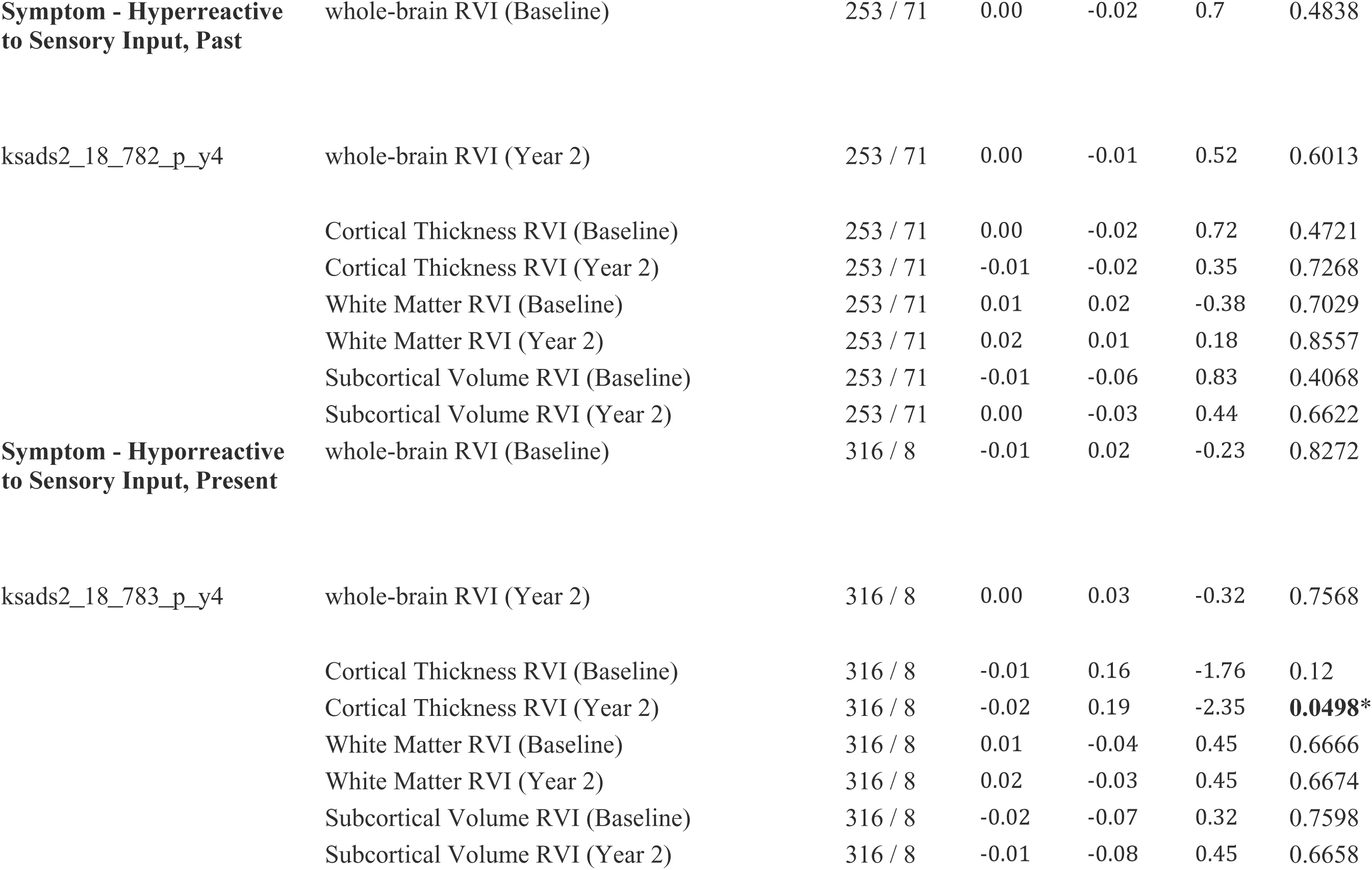
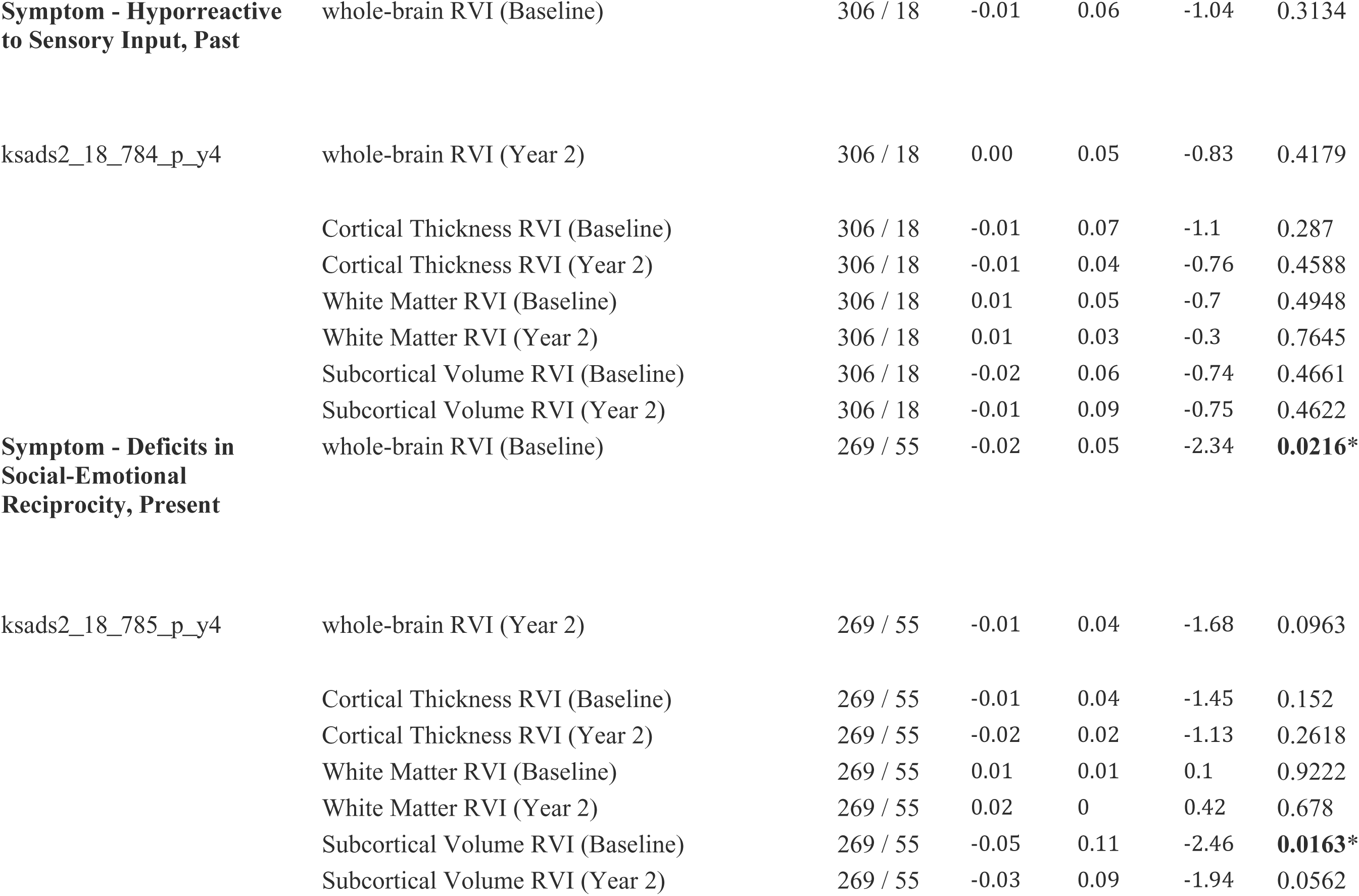
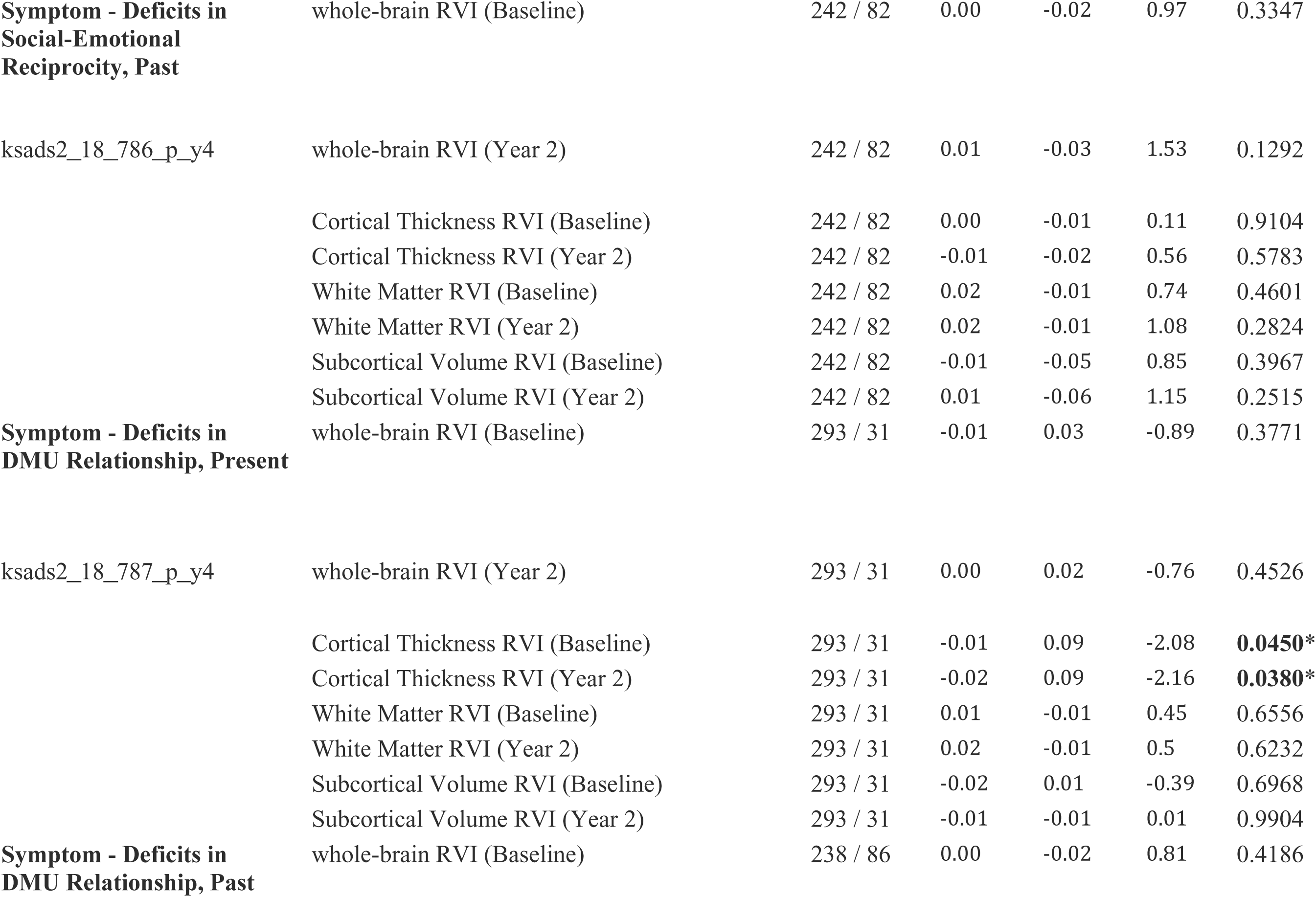
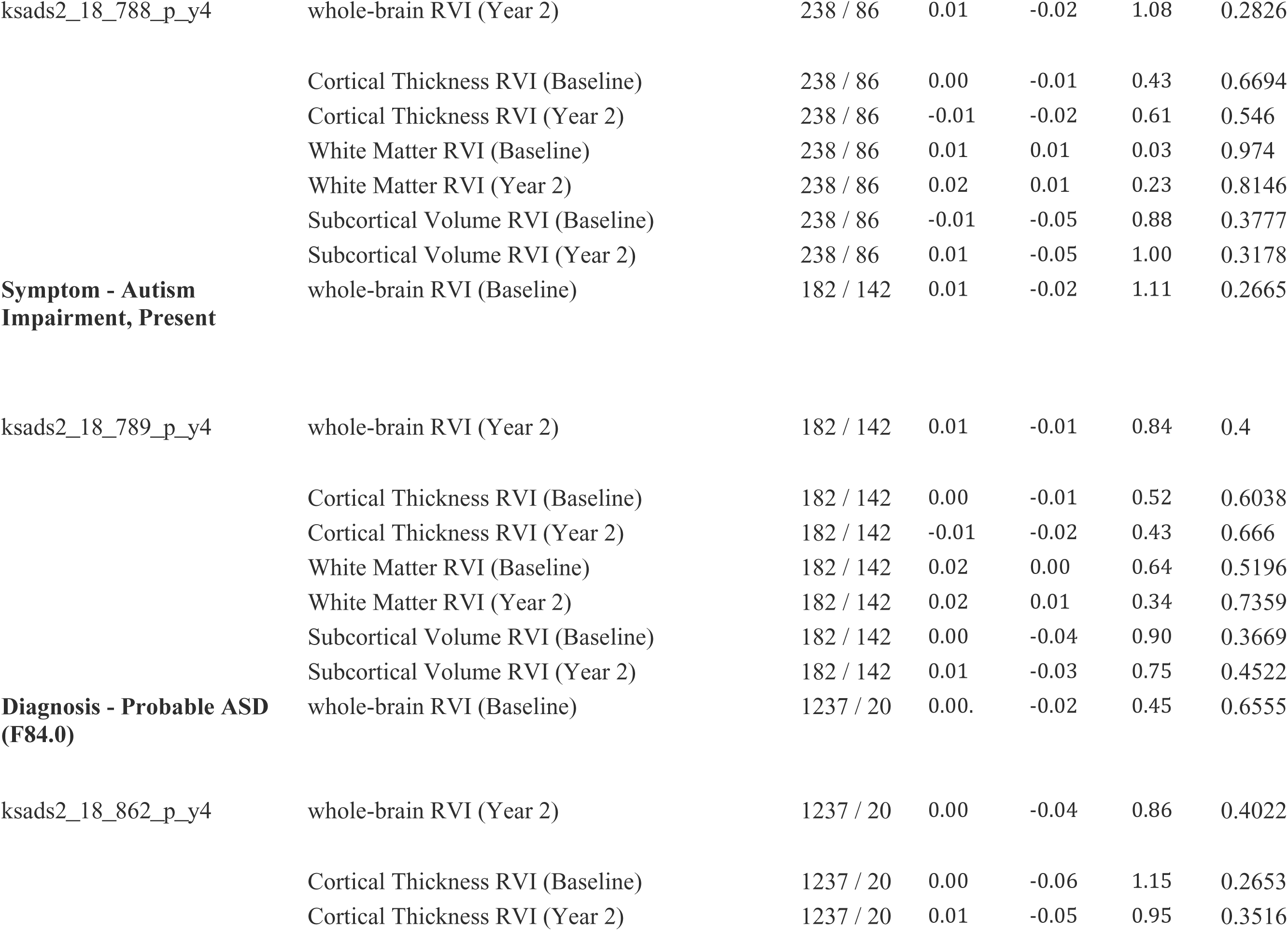
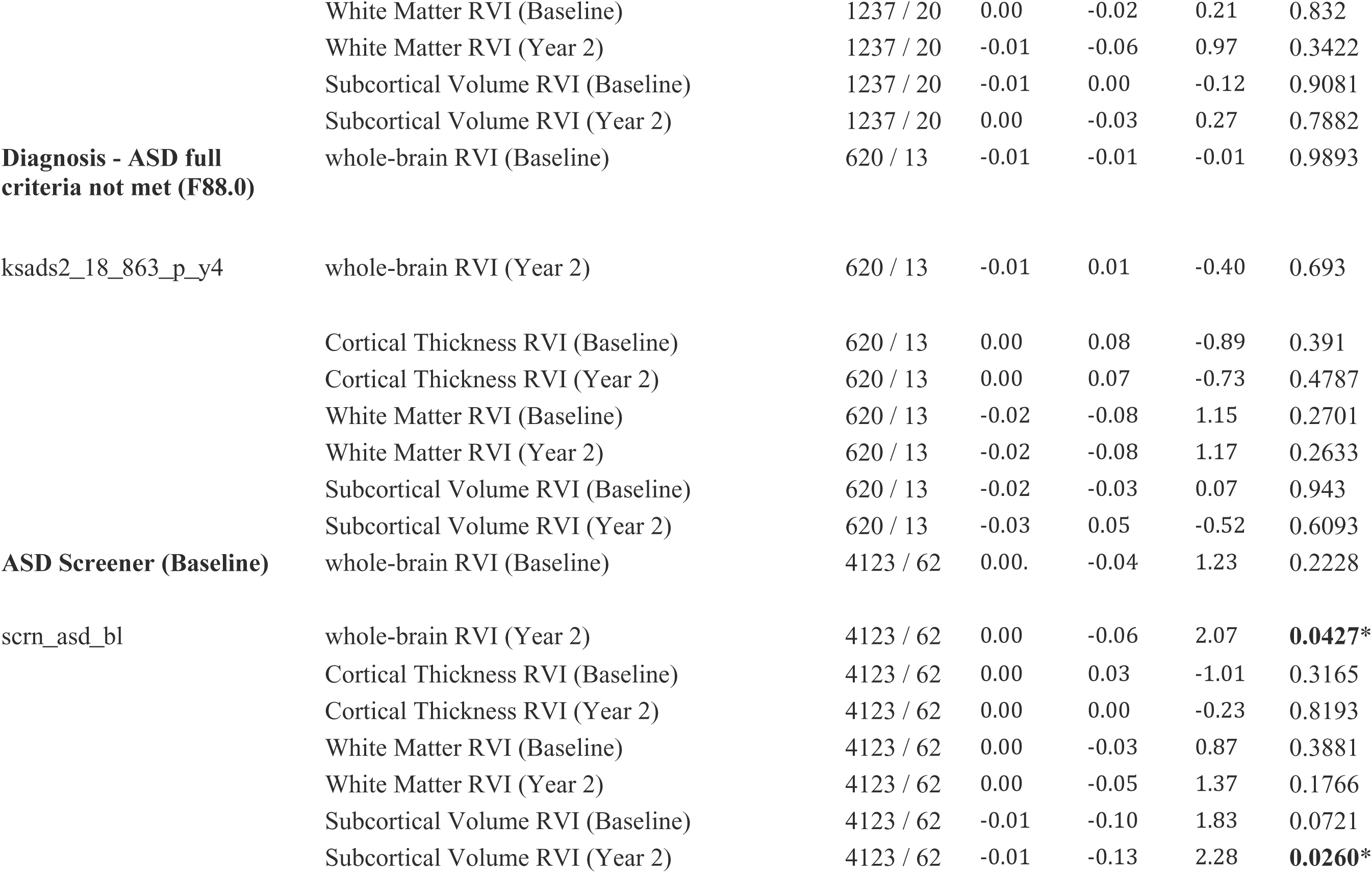
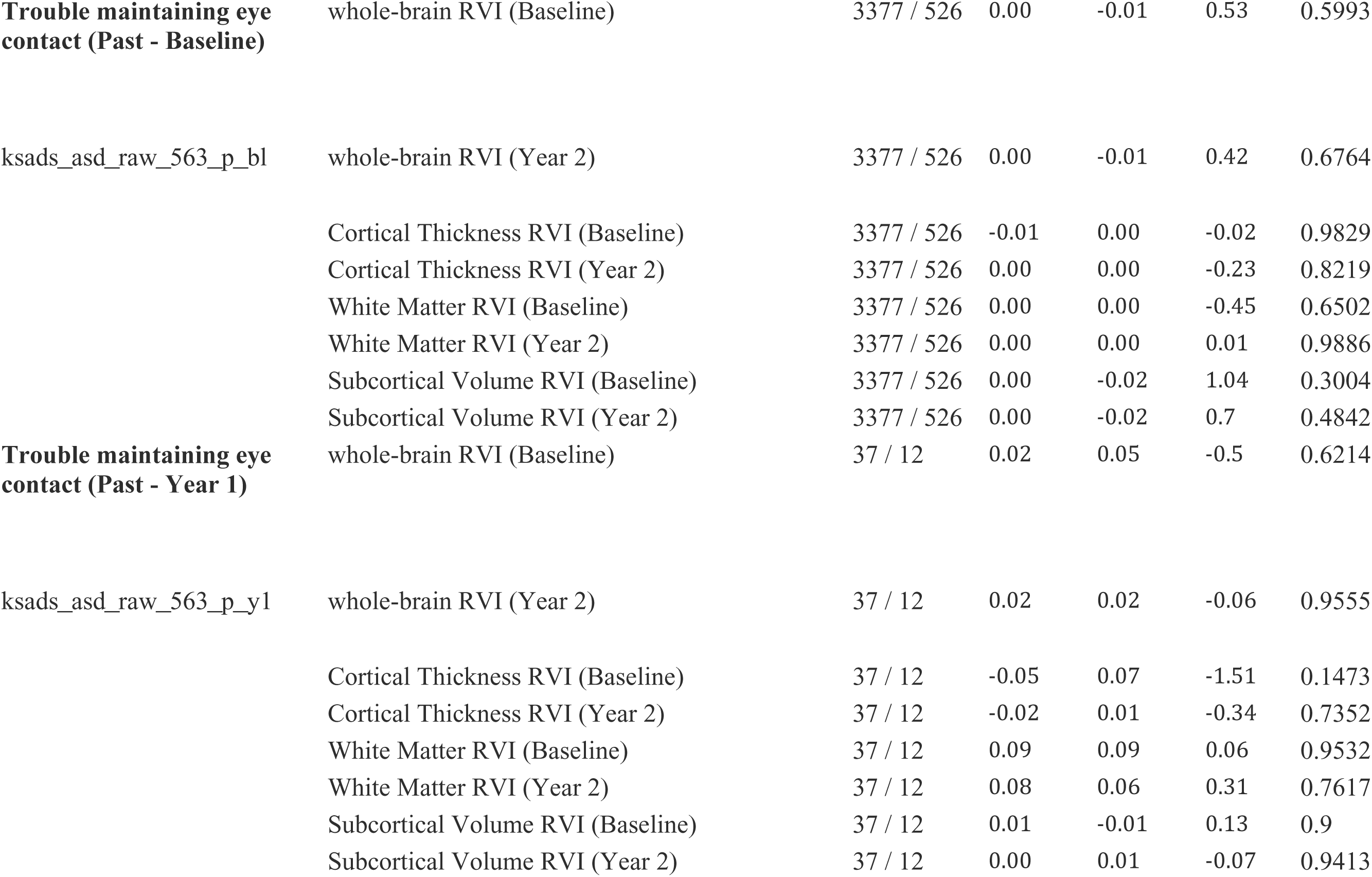
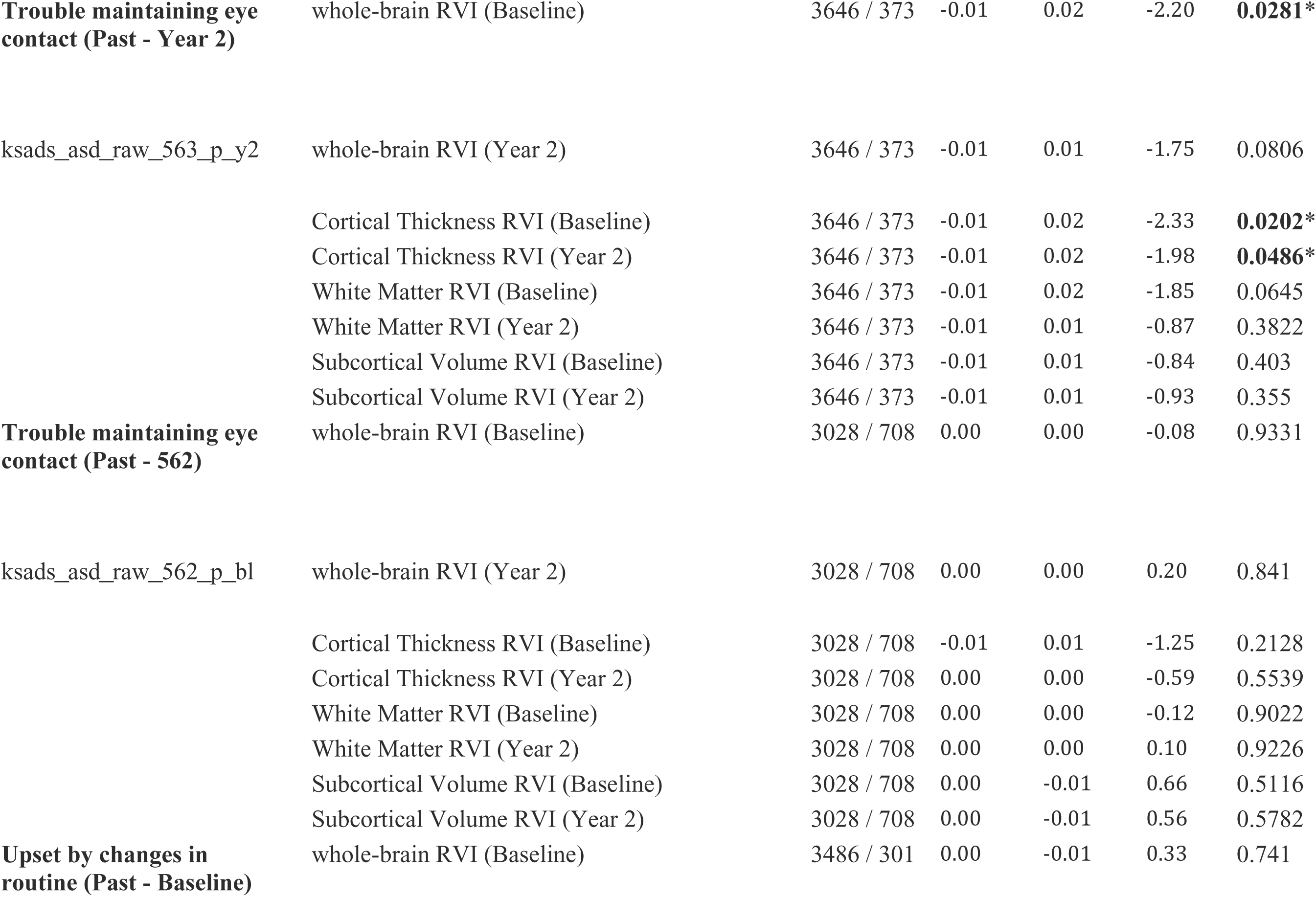
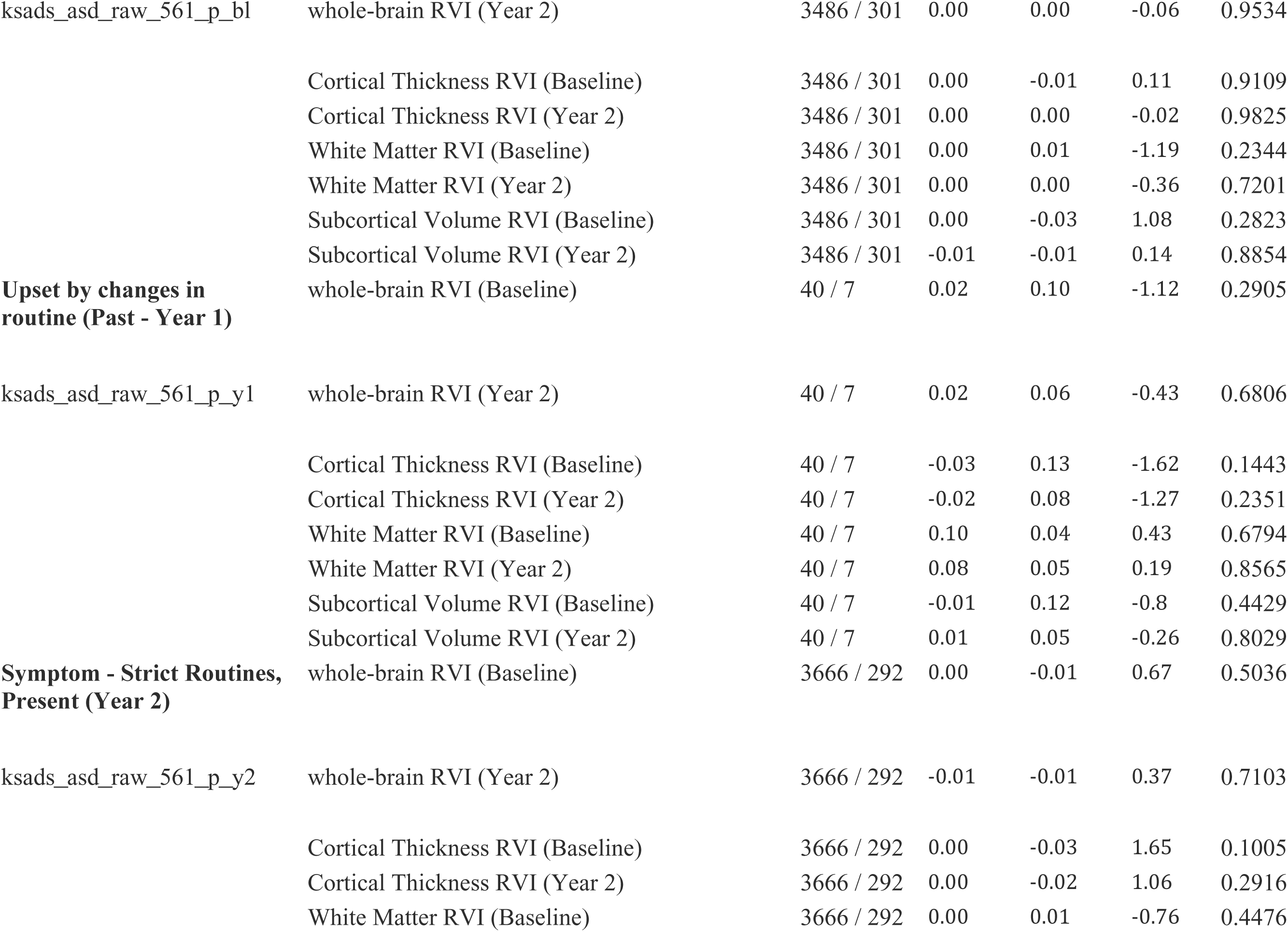
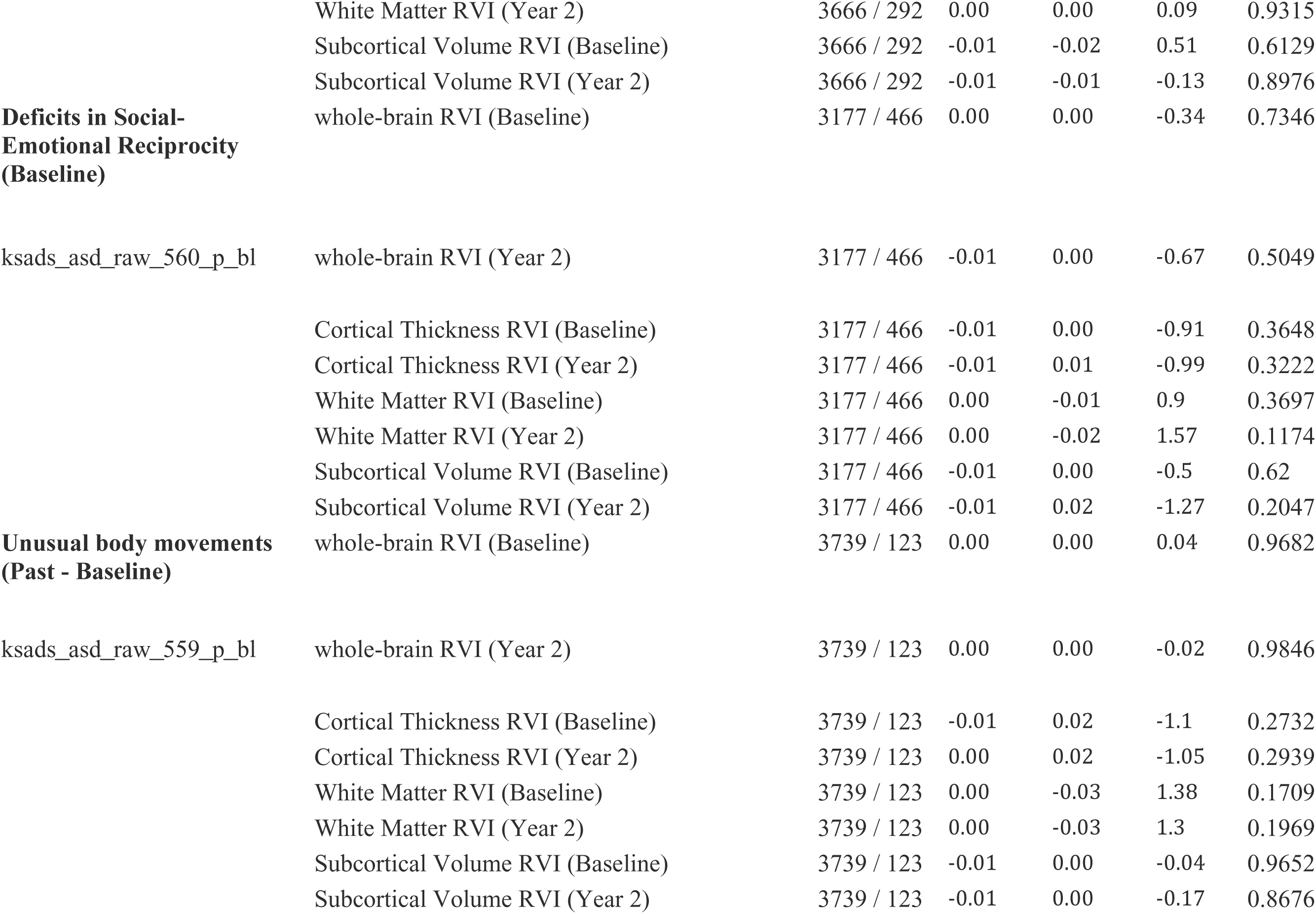
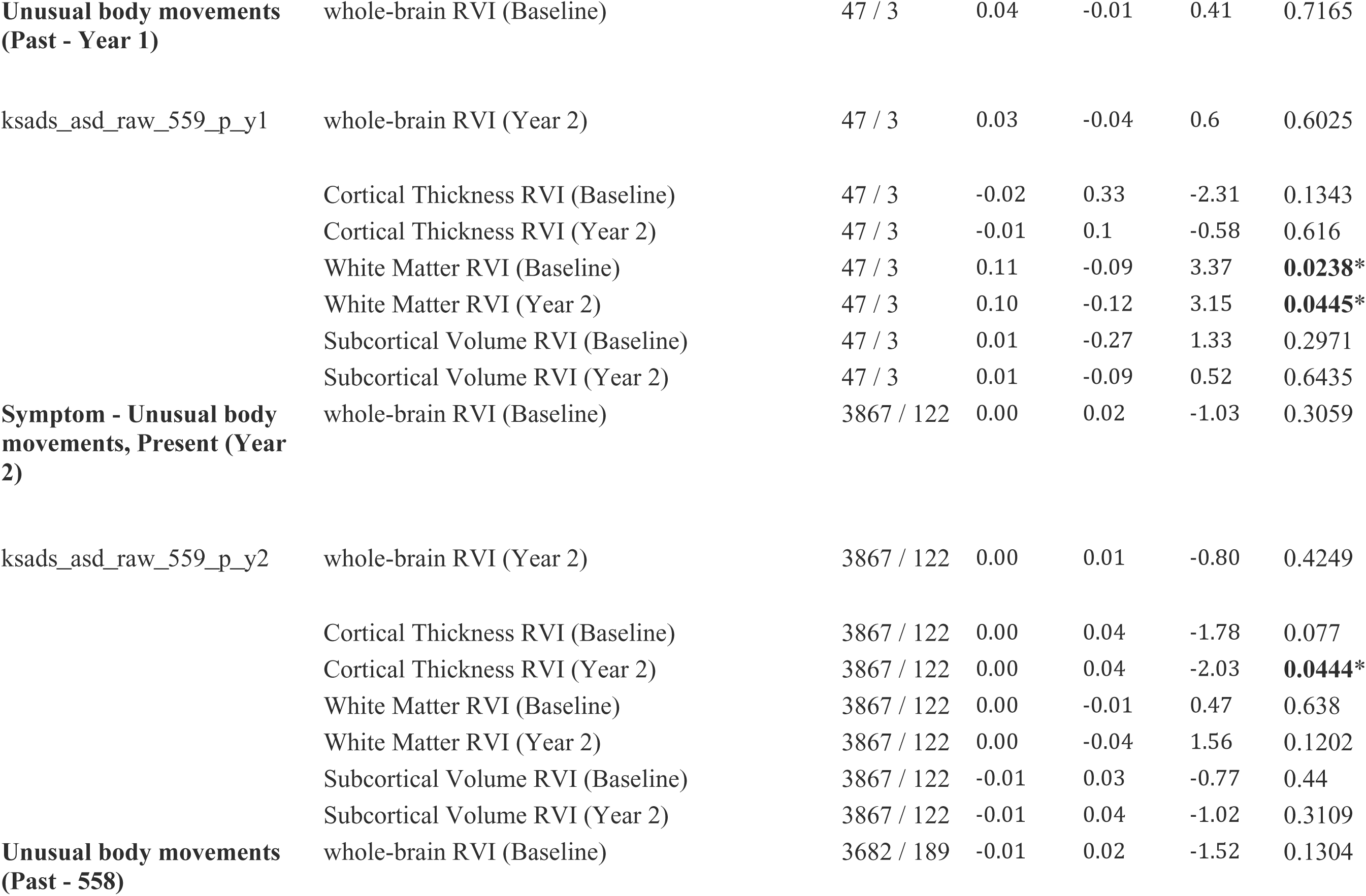
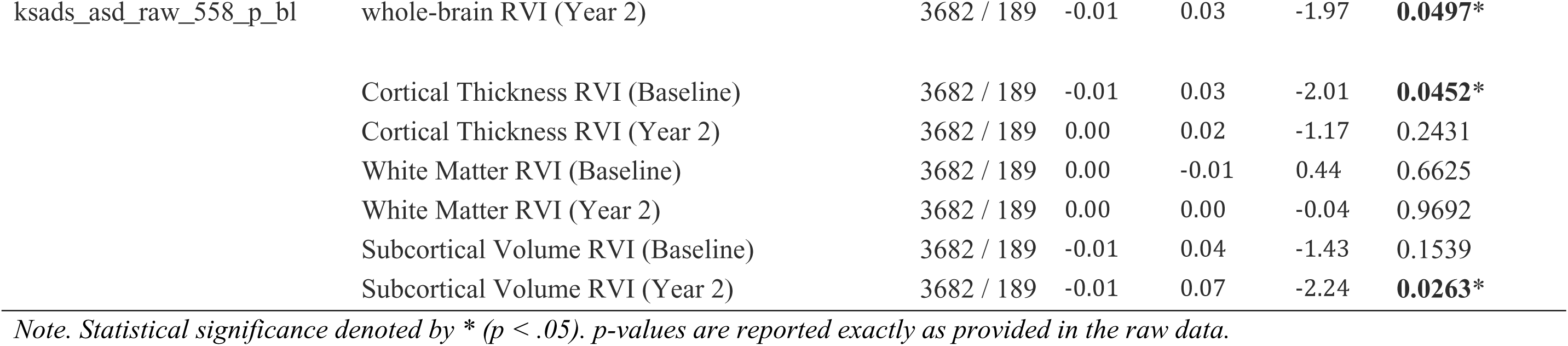
Group Differences (T-Tests) for Dichotomous KSADS Symptom Variables and RVI-ASD Variables.

**Supplementary Table 8.**
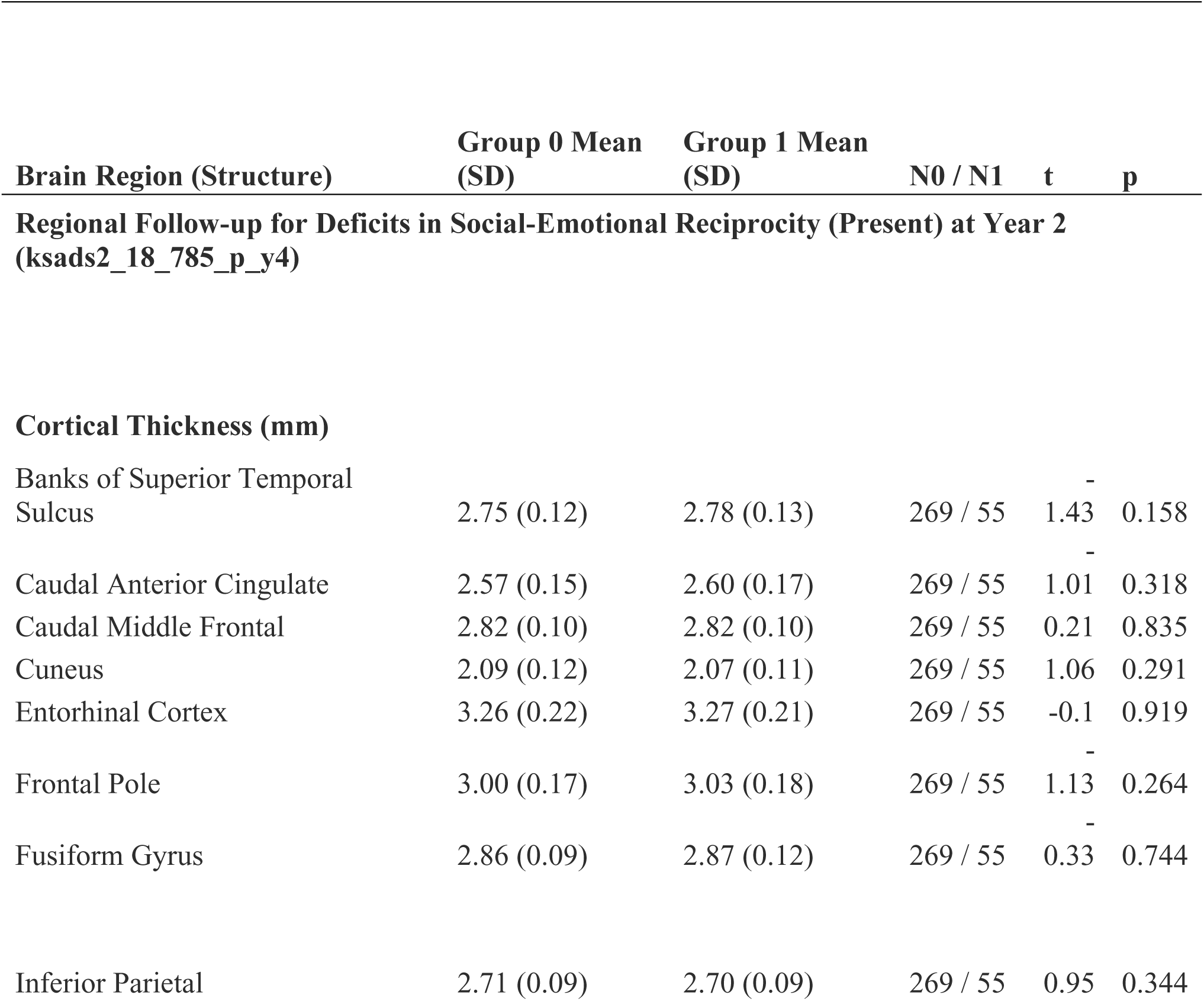
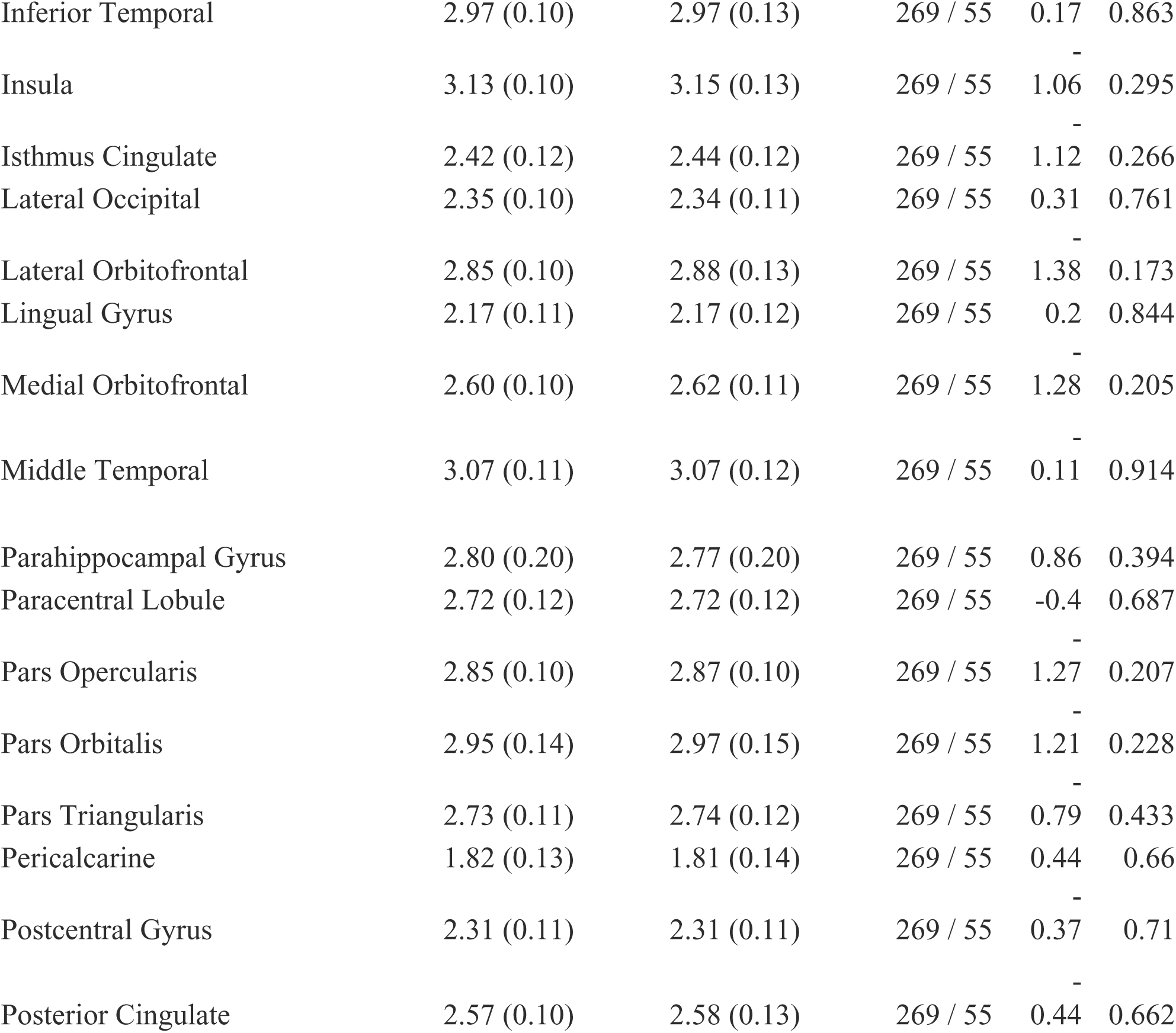
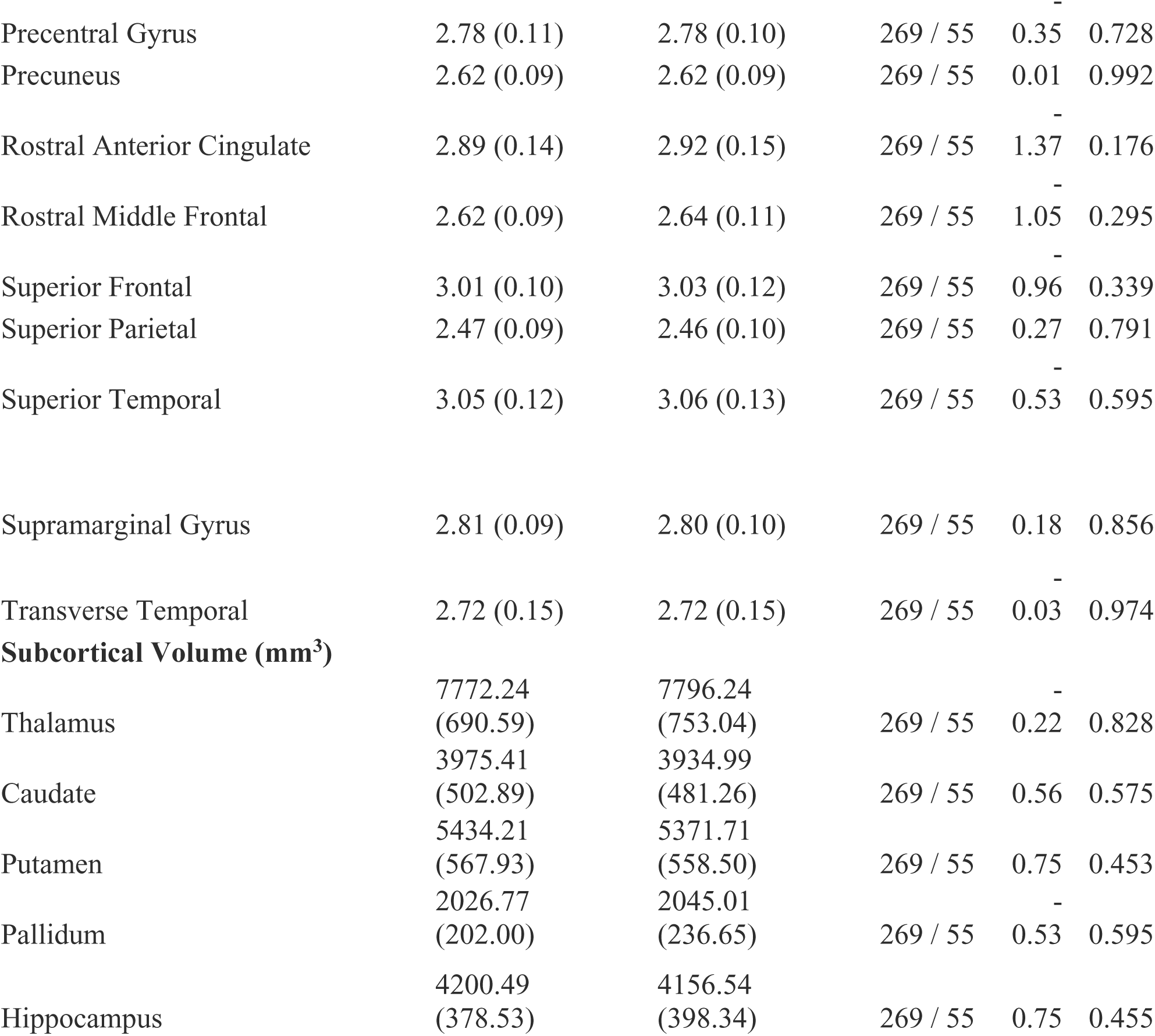
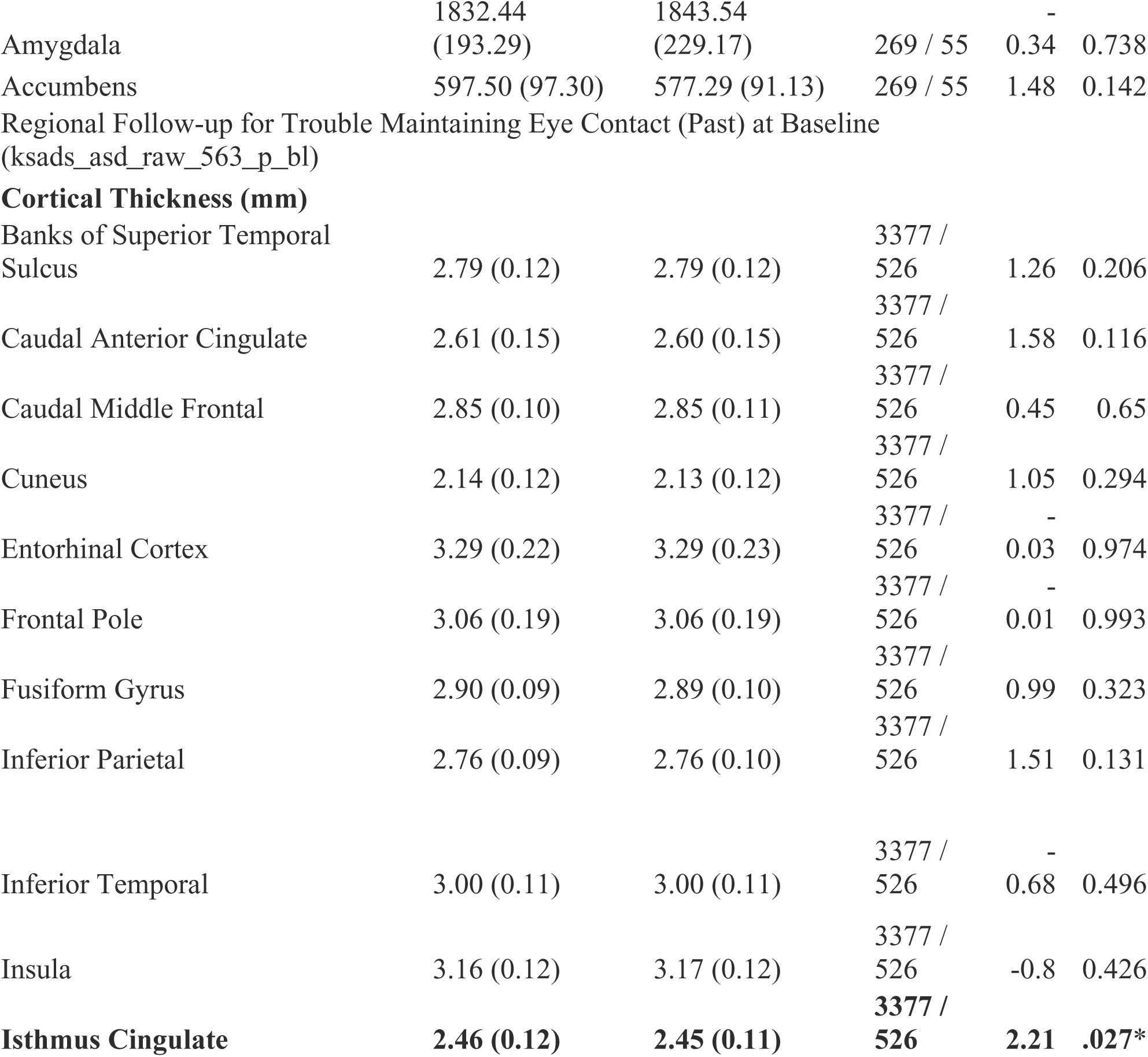
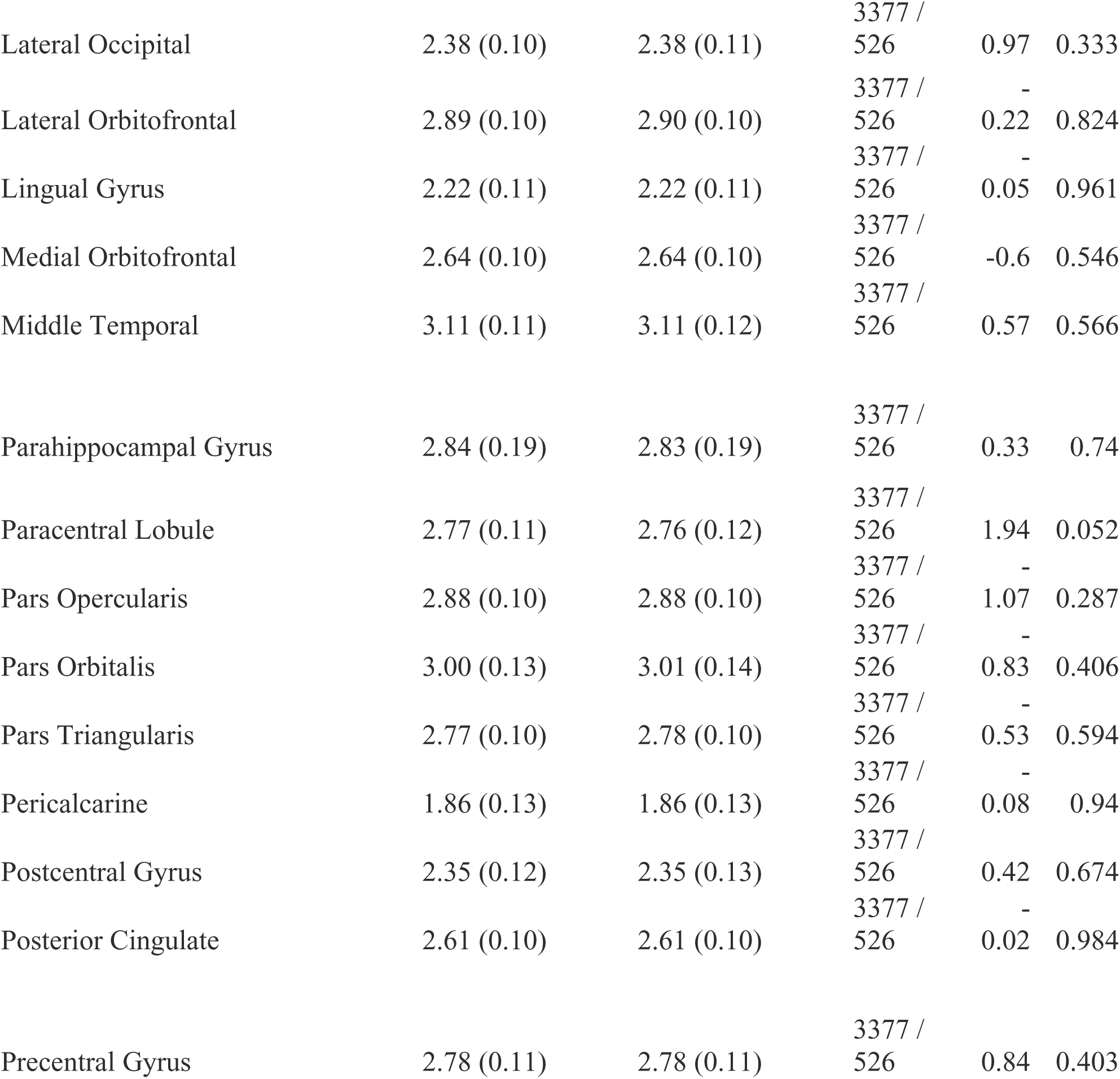
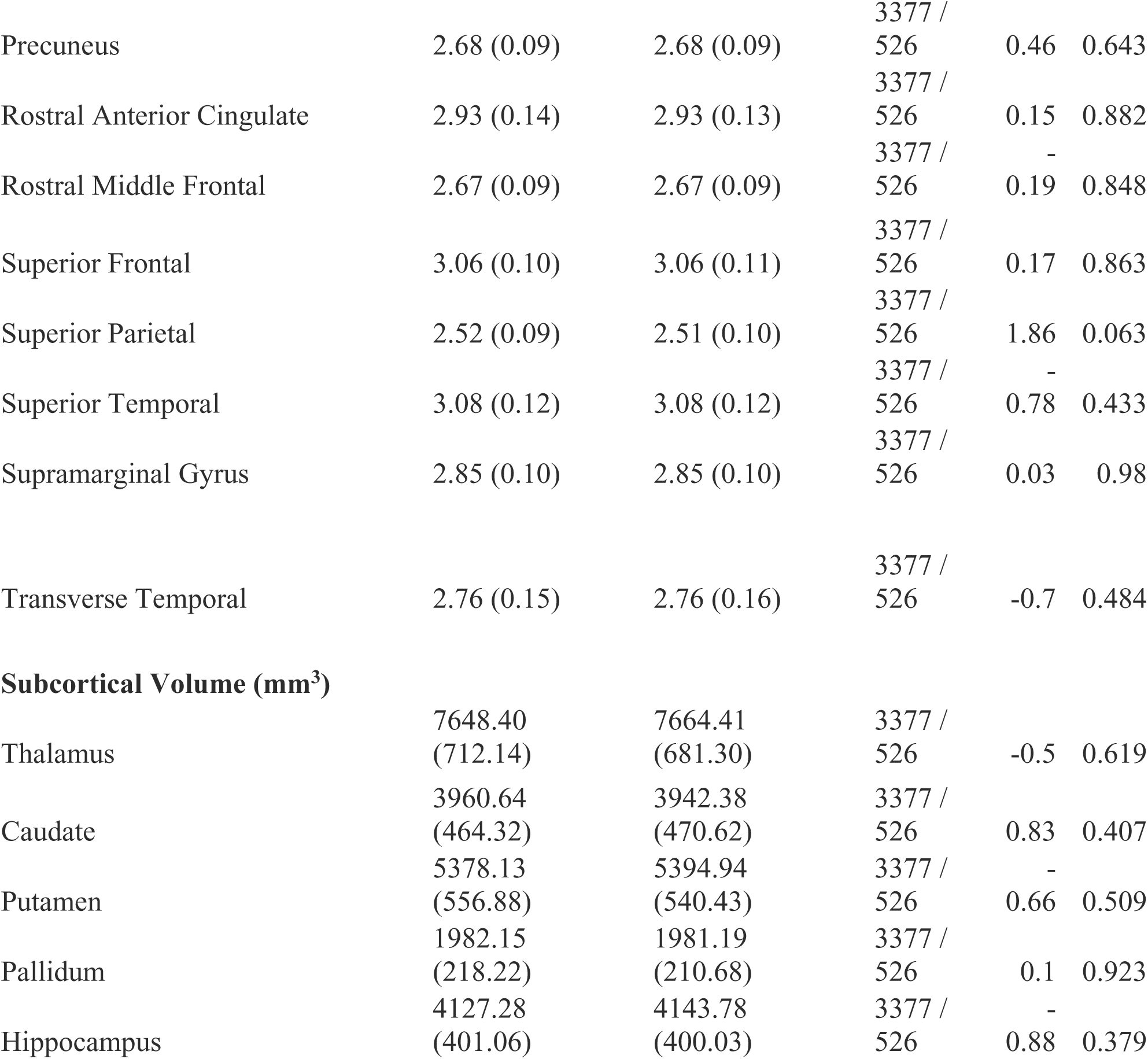
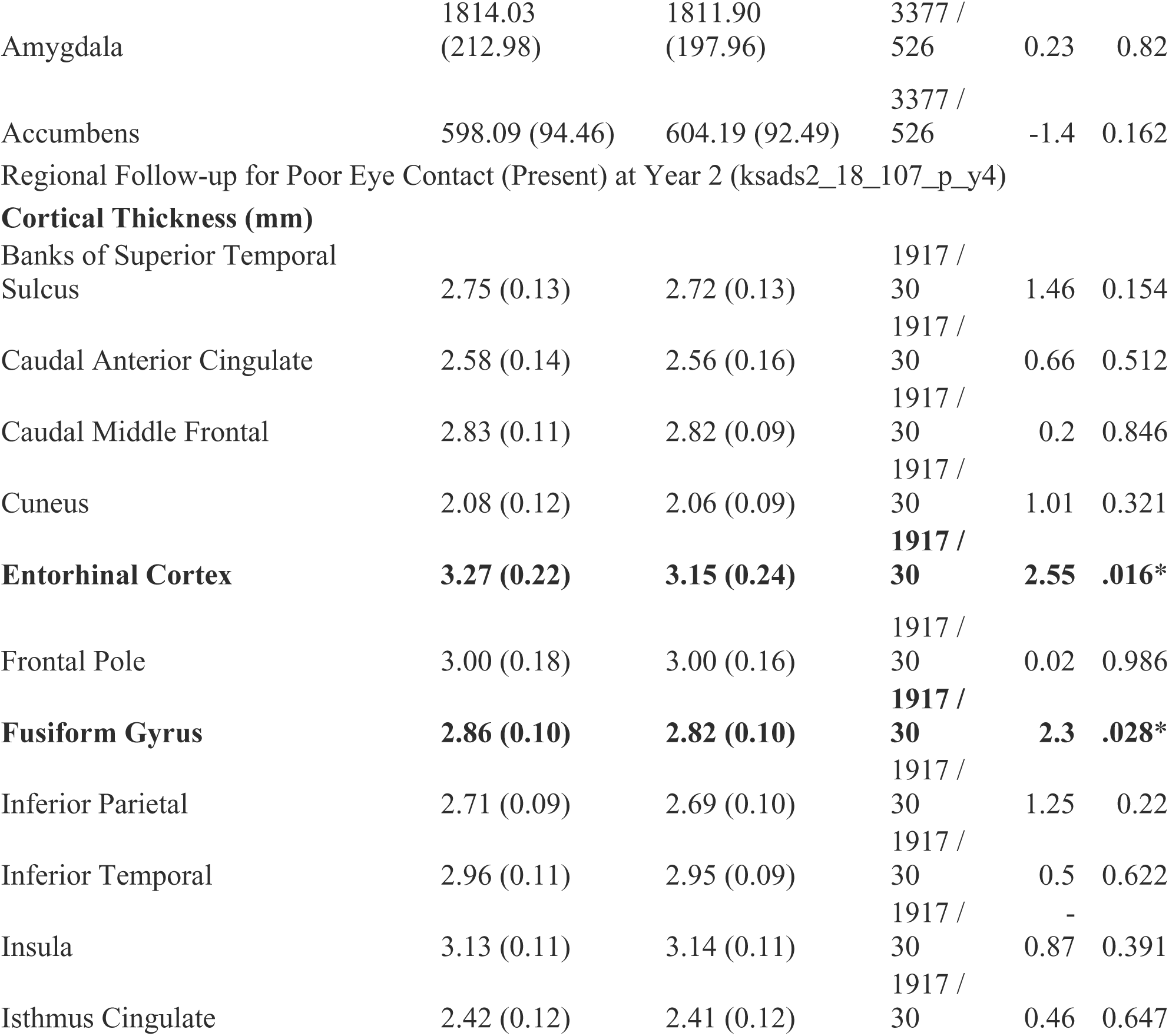
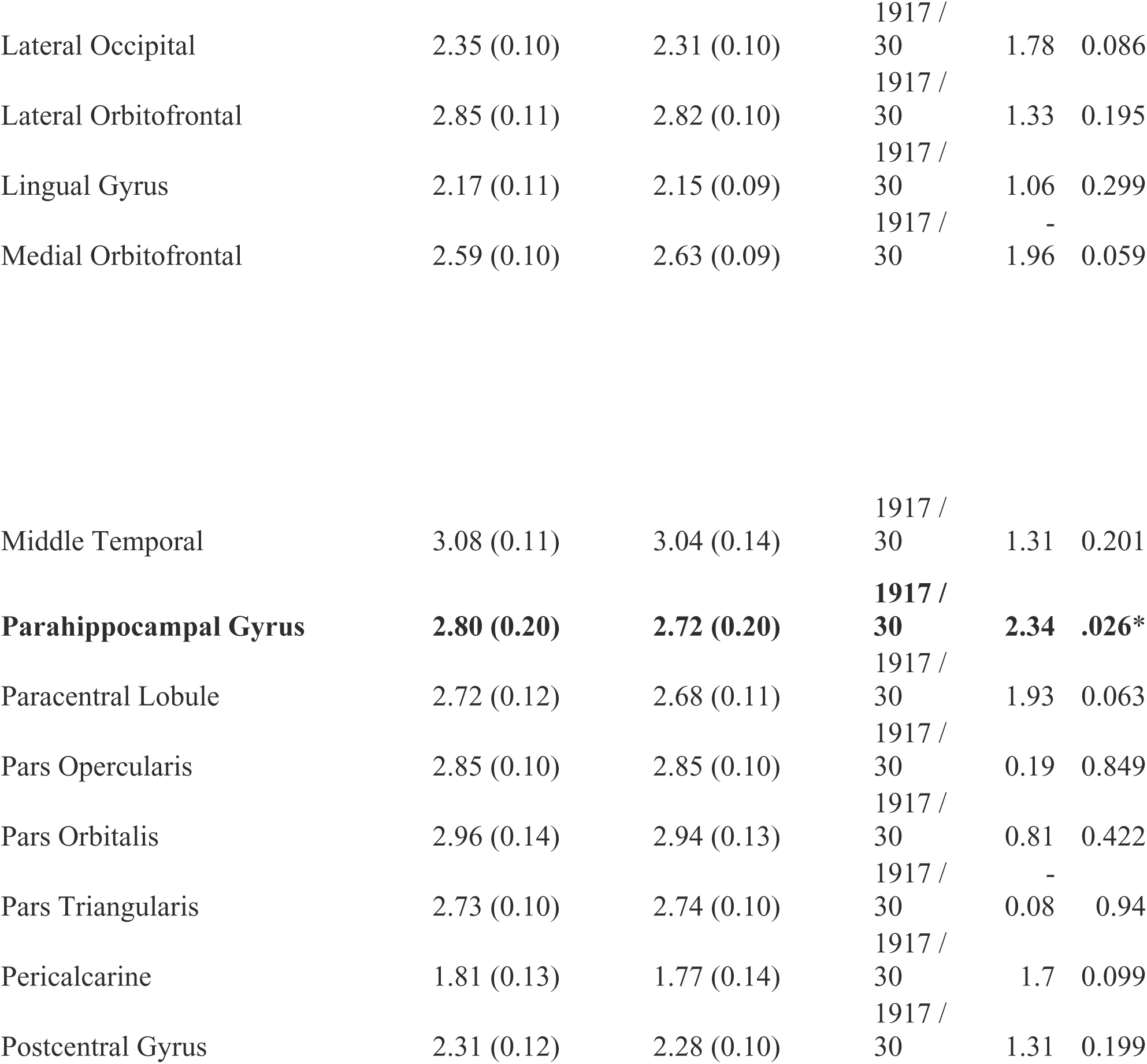
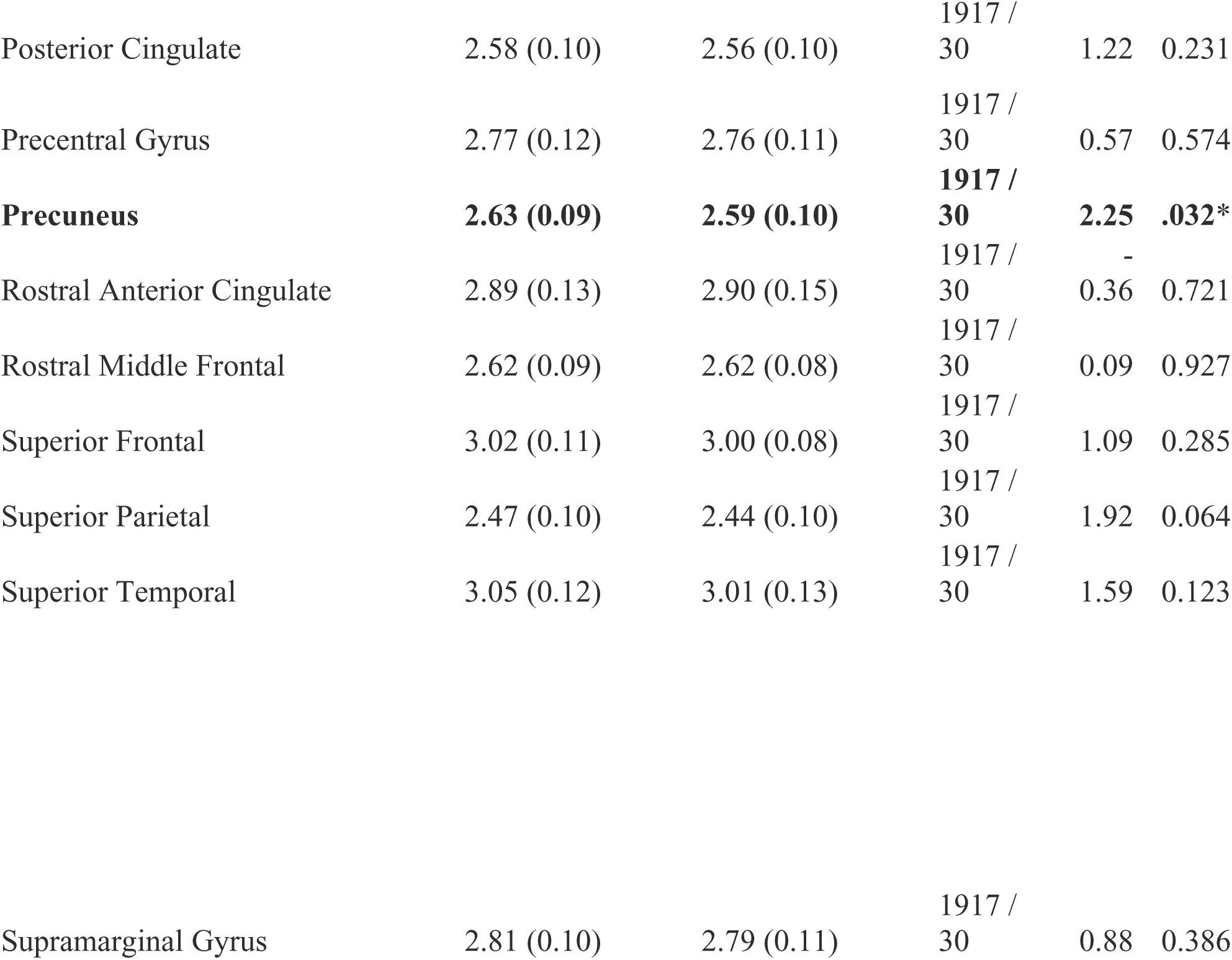
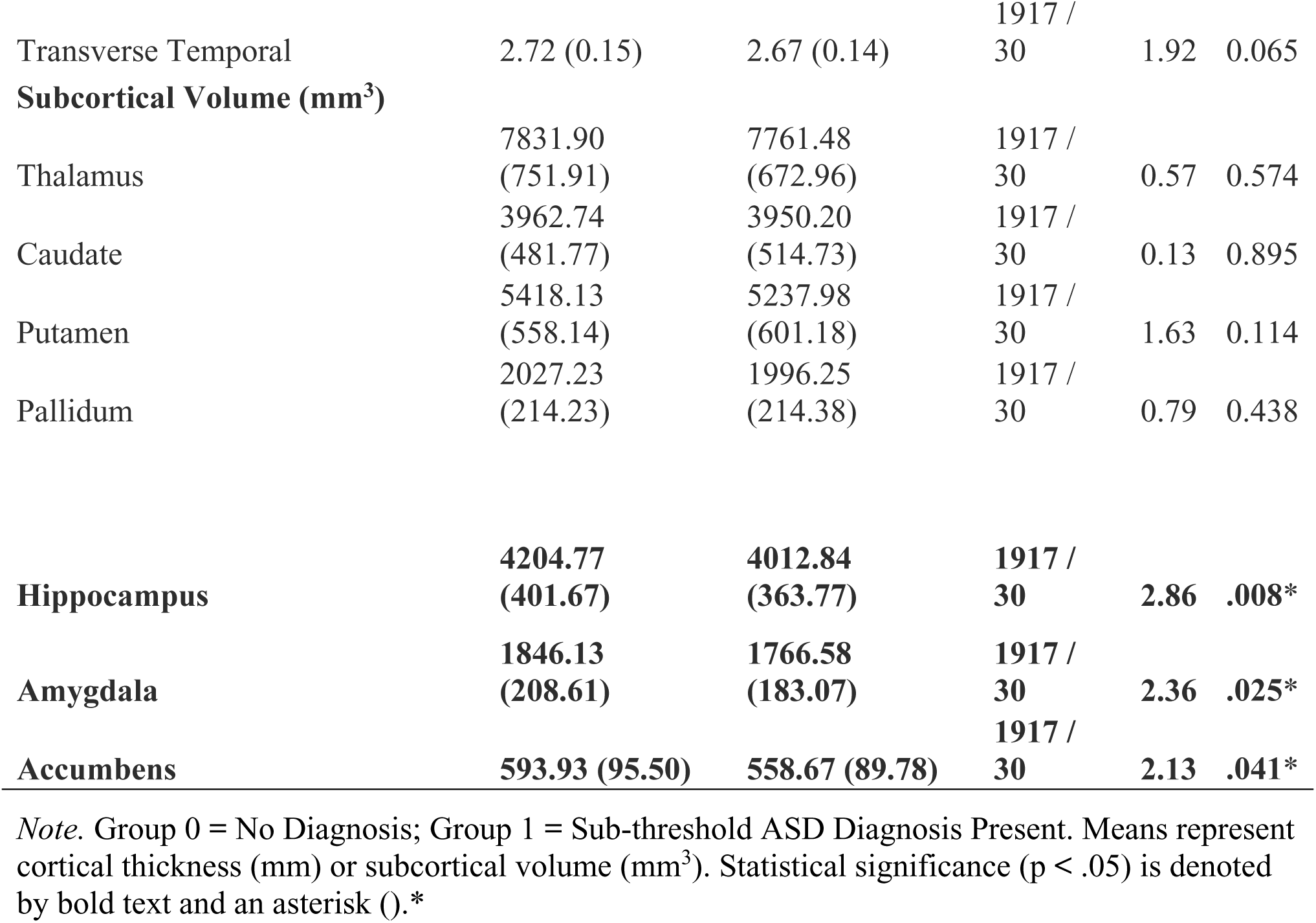
Complete Bilateral Regional Drivers of Significant whole-brain RVI-ASD Associations.

**Supplementary Table 9.**
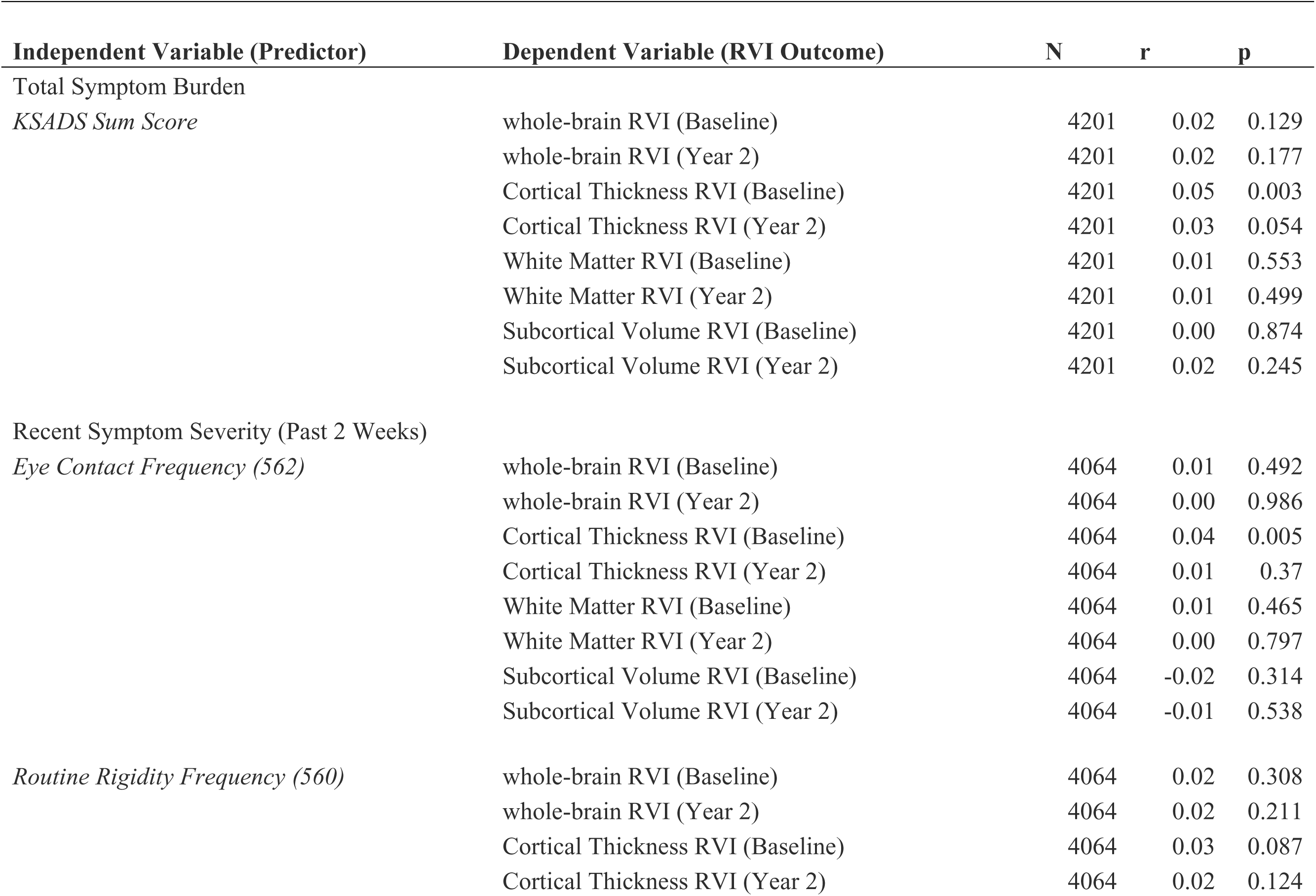
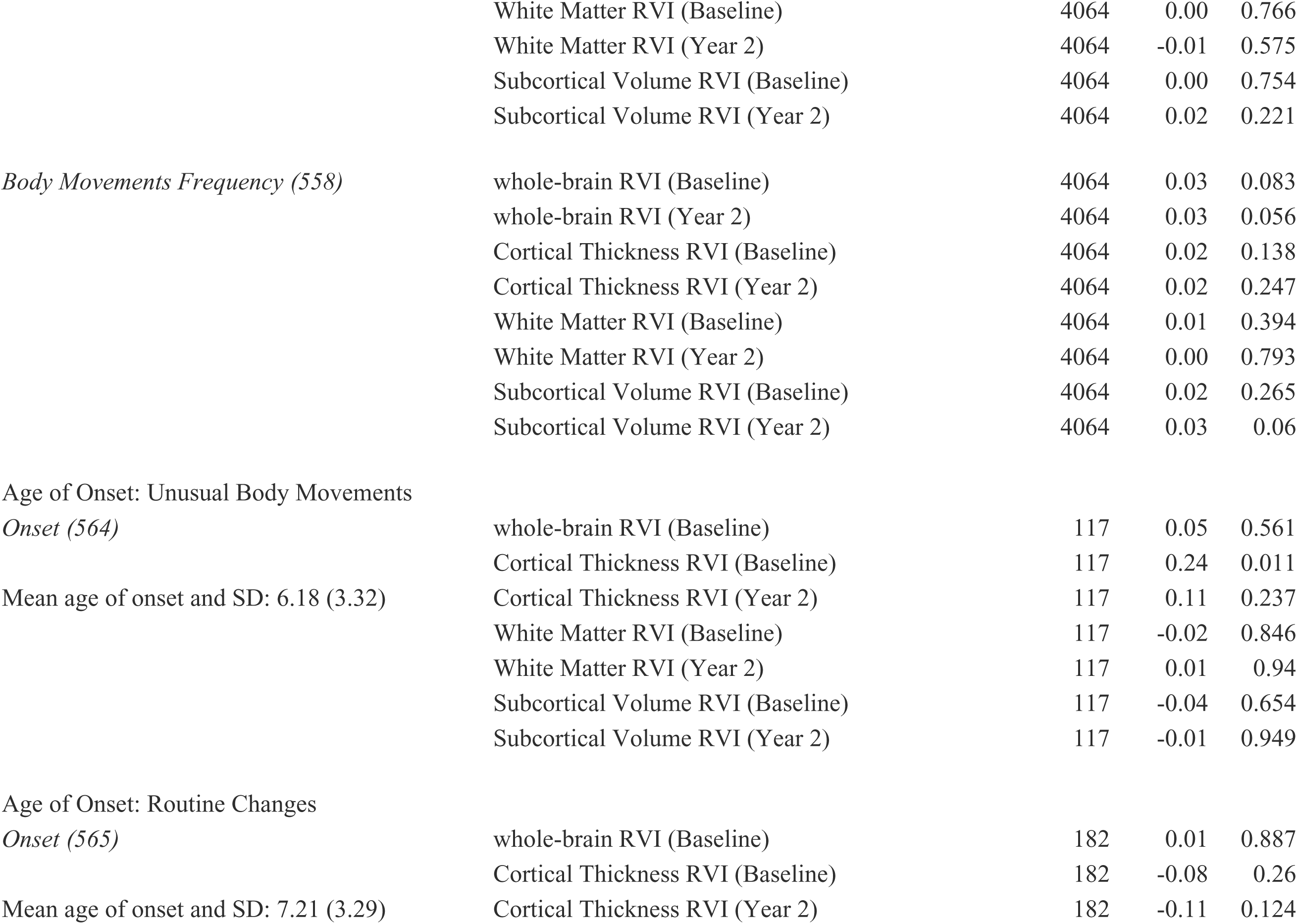
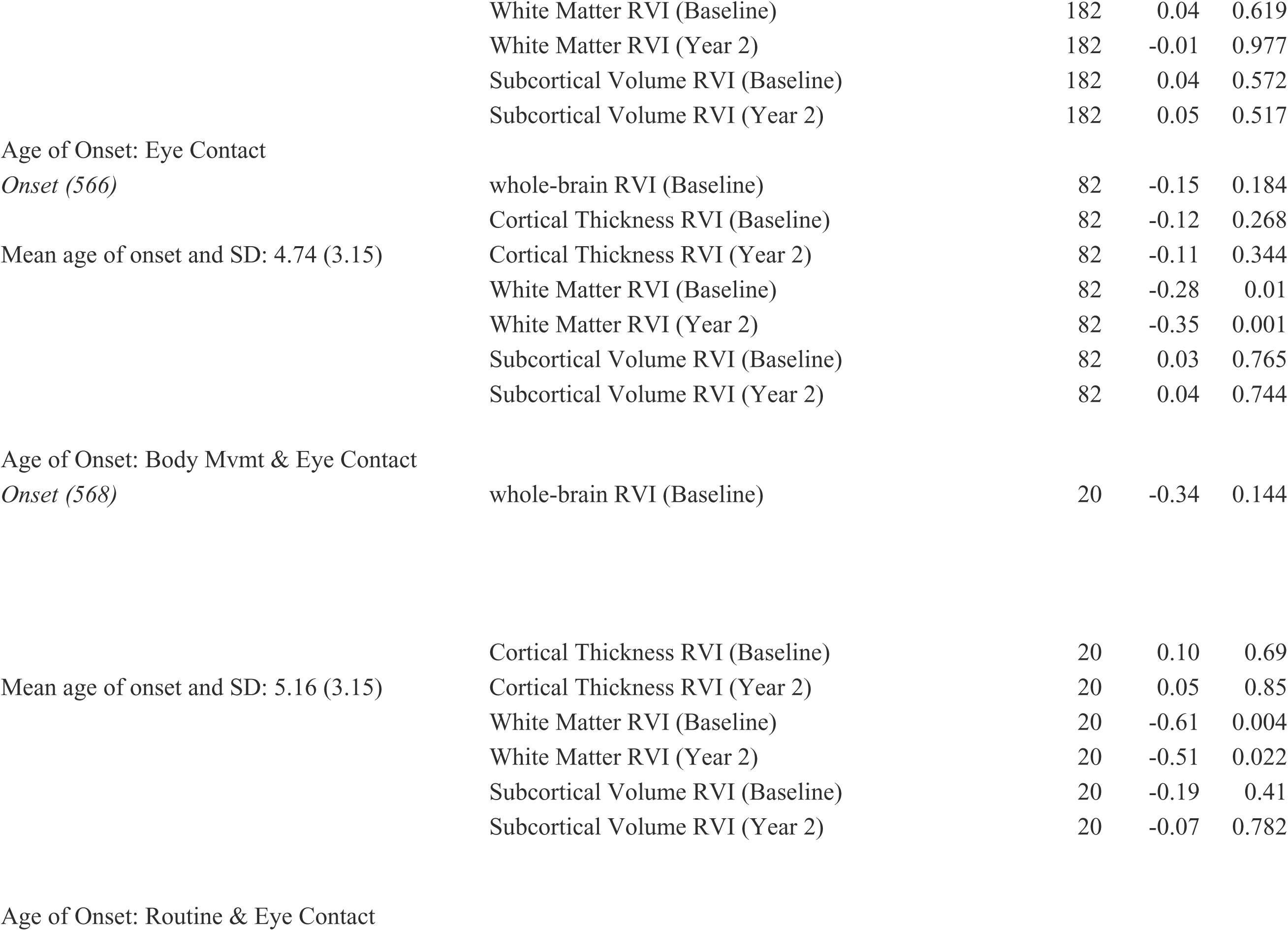
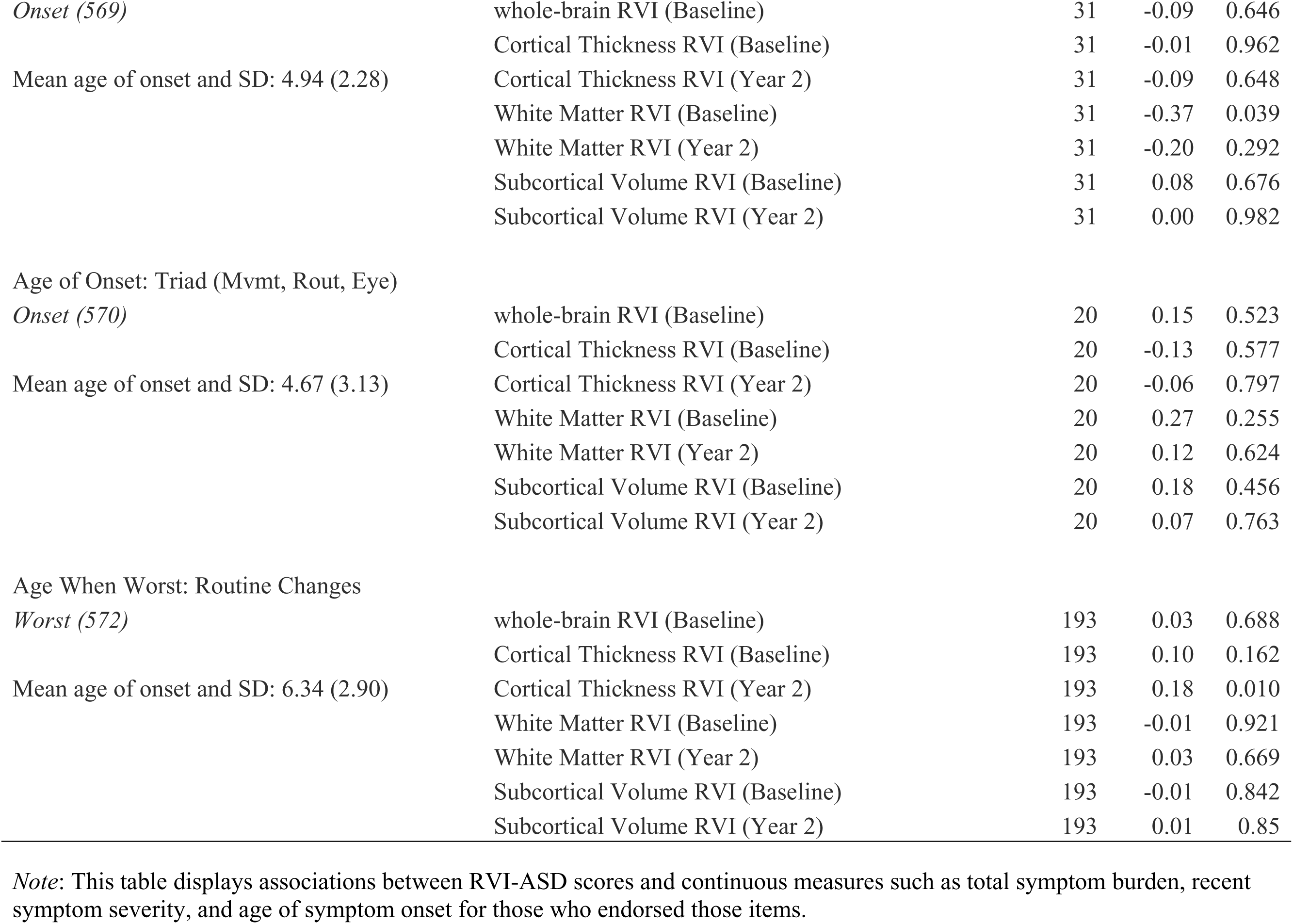
Correlations with Continuous KSADS Variables (Age of Onset & Symptom Sum Scores) and RVI-ASD Variables.

## Supplementary Materials: Clinical and Behavioral Measures

### 1. Kiddie Schedule for Affective Disorders and Schizophrenia (KSADS-COMP)

Parents reported on their child’s current and lifetime mental health using the computerized Kiddie Schedule for Affective Disorders and Schizophrenia for DSM-5 (KSADS-COMP). As noted in the main text, the ABCD study protocol excluded participants with severe autism unable to comply with scanning procedures (Kaufman et al., 1997; Townsend et al., 2020). Consequently, our analysis prioritized specific symptom dimensions over categorical diagnostic labels to capture phenotypic variance within this population-based sample. We examined the presence (past or current) of core autism phenotypes, and symptom burden and developmental trajectories. Due to the conditional branching logic of the KSADS-COMP, follow-up questions regarding Age of Onset, Symptom Severity, and specific Symptom Codes were only administered to the subset of parents who endorsed the primary screening items. Consequently, analyses involving both continuous variables and dichotomous group comparisons were restricted to the subsample of participants possessing the specific symptom (sample sizes ranging from N=3 to N=373; see Supplemental Appendix). We report the sample size (N) for all sub-analyses to contextualize the magnitude of correlation coefficients (r) and difference statistics (t) relative to their statistical significance values (p). This ensures that large effect sizes driven by small subsamples are interpreted with appropriate context.

#### A. Diagnostic Categories

We analyzed categorical diagnostic variables to identify participants meeting criteria for autism spectrum disorder or related neurodevelopmental conditions.

- **Confirmed Autism Spectrum Disorder (F84.0):** ksads2_18_861
- **Probable ASD:** ksads2_18_862
- **Other Specified Neurodevelopmental Disorder (F88.0):** ksads2_18_863
- **Administrative Label (F88.0):** ksads_18_903 (Note: This variable indicates “full criteria not assessed” and is treated as an administrative label rather than a confirmed clinical diagnosis).

#### B. Dichotomous Symptom Variables

We examined the presence (past or current) of core autism phenotypes derived from two distinct sources within the KSADS supplement:

### 1. Symptom Codes (Binary: 0=Absent, 1=Present)

Specific binary symptom codes were analyzed to capture the two core domains of ASD:

○ **Social Communication Deficits:**

▪ Poor eye contact: ksads2_18_107 (Present) / ksads2_18_108 (Past)
▪ Deficits in social-emotional reciprocity: ksads2_18_785 (Present) / ksads2_18_786 (Past)
▪ Deficits in developing/maintaining relationships: ksads2_18_787 (Present) / ksads2_18_788 (Past)
○ **Restricted/Repetitive Behaviors:**

▪ Unusual body movements: ksads2_18_103 (Present) / ksads2_18_104 (Past)
▪ Strict routines: ksads2_18_105 (Present) / ksads2_18_106 (Past)
▪ Restricted interests: ksads2_18_779 (Present) / ksads2_18_780 (Past)
▪ Sensory reactivity (Hyper/Hypo): ksads2_18_781–784 (Present/Past)

### 2. Lifetime History Items (Raw Interview Data)

We analyzed raw interview items regarding the lifetime presence of specific traits, independent of the final symptom code assignment. These items capture sub-threshold or historical presentations:

○ *Eye Contact:* “Was there ever a time your child had trouble maintaining eye contact?” (ksads_asd_raw_563_p)
○ *Routines:* “Was there ever a time your child was frequently easily upset by changes in routines?” (ksads_asd_raw_561_p)
○ *Body Movements:* “Was there ever a time your child frequently did unusual body movements?” (ksads_asd_raw_559_p)

#### C. Derived Continuous Measures

To characterize symptom burden and developmental trajectories, we calculated three types of continuous metrics:

### 1. Symptom Burden (Sum Score)

A cumulative **KSADS Sum Score** (ksads_sumscore) was generated by summing positive responses across all screened autism items (Symptom Codes and Lifetime History items). This provides a single continuous measure of total ASD-related behavioral burden.

### 2. Recent Symptom Severity (0–3 Scale)

We analyzed the continuous frequency/severity of core symptoms reported in the “past two weeks.” Items were rated on a 4-point scale (0=Not at all; 1=Rarely; 2=Several days; 3=More than half the days; 4=Nearly every day):

○ Trouble maintaining eye contact: ksads_asd_raw_562_p
○ Upset by routine changes: ksads_asd_raw_560_p
○ Unusual body movements: ksads_asd_raw_558_p

### 3. Age of Onset

To investigate developmental gradients, we calculated the precise age of symptom onset using parent-reported dates (Age = Symptom Onset Date - Demographic Birthdate). We extracted onset dates for specific domains and combined presentations:

○ **Single Domain Onset:**

▪ Eye contact deficits: ksads_asd_raw_566_p
▪ Routine rigidity: ksads_asd_raw_565_p
▪ Unusual body movements: ksads_asd_raw_564_p
○ **Combined/Comorbid Onset:**

▪ Movements + Routine rigidity: ksads_asd_raw_567_p
▪ Movements + Eye contact deficits: ksads_asd_raw_568_p
▪ Rigidity + Eye contact deficits: ksads_asd_raw_569_p
▪ **Triad (All three):** ksads_asd_raw_570_p

### 2. Child Behavior Checklist (CBCL)

The CBCL is a standardized parent-report questionnaire used to assess emotional and behavioral problems in children. It provides empirically based syndrome scales (Achenbach, 2009).

- **Variables Used:**

○ **Syndrome Scales:** Anxious/Depressed, Withdrawn/Depressed, Social Problems, Attention Problems, Aggressive Behavior.
○ **DSM-Oriented Scales:** Depressive Problems, Anxiety Problems, Somatic Problems, ADHD Problems, Oppositional Defiant Problems, Conduct Problems.
○ **Scoring:** Parents rate items on a 3-point scale (0 = Not True, 1 = Somewhat or Sometimes True, 2 = Very True or Often True). Raw scores were converted to T-scores for analysis.

### 3. Social Responsiveness Scale (SRS)

The SRS is a parent-report questionnaire that identifies the presence and severity of social impairment associated with Autism Spectrum Disorder (Constantino, 2013).

- **Variables Used:**

○ **Total Score:** A composite measure of social impairment.
○ **Subscales:** Social Awareness, Social Cognition, Social Communication, Social Motivation, and Restricted Interests/Repetitive Behavior.
○ **Specific Items:** Difficulty with routines, unusual communication, social cues, conversation flow, difficulty making friends, and social withdrawal.
○ **Scoring:** Items are rated on a 4-point Likert scale (1 = Not True, 2 = Sometimes True, 3 = Often True, 4 = Almost Always True).

### 4. Demographic and Environmental Variables

#### A. Socioeconomic Status (SES) Measures

- **Income-to-Needs Ratio (INR)**

○ **Source:** Parent Demographics Survey (Barch et al., 2021).
○ **Scoring:** This variable is a standardized ratio calculated by dividing the **Total Household Income** by the **Federal Poverty Level (FPL)** corresponding to the specific household size (based on the U.S. Department of Health and Human Services poverty guidelines for the year of data collection).
○ **Calculation Note:** Since income in the ABCD study is reported in bins (e.g., “50,000 through 74,999”), the median value of the endorsed bin (e.g., 62,500) was used as the numerator.
○ **Interpretation:** A ratio of 1.0 indicates the family is exactly at the poverty line. Values < 1.0 indicate living in poverty, while values > 1.0 indicate income above the poverty line (e.g., 3.0 = 300% of the poverty level).
- **Financial Adversity (Financial Difficulty)**

○ **Source:** Parent Demographics Survey (Barch et al., 2021).
○ **Scoring:** A cumulative **sum score (Range: 0–7)** representing the total count of distinct financial hardships experienced by the family in the past 12 months.
○ **Items Assessed:** Parents answered “Yes” (1) or “No” (0) to seven specific hardships:

1. Inability to afford food.
2. Utilities turned off due to non-payment.
3. Inability to pay full rent or mortgage.
4. Eviction from home.
5. Phone service disconnected due to non-payment.
6. Inability to afford a doctor/hospital visit.
7. Inability to afford a dentist visit.
○ **Interpretation:** Higher scores indicate greater financial adversity and instability.

#### B. Perinatal and Developmental History

- **Birth Complications Composite**

○ **Source:** Developmental History Questionnaire (dhx0; Barch et al., 2021).
○ **Scoring:** A **composite sum score** quantifying the total number of adverse events reported during pregnancy or delivery.
○ **Items Included:** Premature birth (gestational age < 37 weeks), need for oxygen/ventilator at birth, jaundice requiring treatment, incubation/NICU stay > 2 days, and specific pregnancy complications (e.g., pre-eclampsia, placenta previa).
○ **Interpretation:** Higher scores reflect a more biologically adverse perinatal environment.
- **Delivery Method (C-Section)**

○ **Source:** Developmental History Questionnaire (Barch et al., 2021).
○ **Scoring:** A binary variable derived from the item asking “How was the child delivered?”
○ **Coding:**0.00= Vaginal Delivery; 1 = Cesarean Section (C-section).

#### D. Environmental Exposures

- **Lead Exposure Risk**

○ **Source:** ABCD Residential History Derived Scores (Barch et al., 2021). (reshist_addr1_lead_risk).
○ **Method:** This variable is a geocoded proxy for lead exposure risk based on the participant’s primary residential address. It utilizes **housing age data** from the American Community Survey (ACS) at the census tract level.
○ **Scoring:** The score represents the estimated percentage of housing units in the census tract built before 1980 (a primary risk factor for lead paint).
○ **Interpretation:** Higher scores indicate an older housing stock in the neighborhood, serving as a proxy for higher potential lead exposure risk.
- **Neighborhood Safety**

○ **Source:** Parent and Youth Neighborhood Safety Questionnaire (Barch et al., 2021).
○ **Scoring:** A **mean composite score** derived from three items assessing perceived safety: (1) “I feel safe walking in my neighborhood,” (2) “My neighborhood is safe from crime,” and (3) “There is a lot of violence in my neighborhood” (reverse-coded).
○ **Scale:** Items are rated on a 1–5 Likert scale (Strongly Disagree to Strongly Agree).
○ **Interpretation:** Higher scores indicate higher perceived neighborhood safety.

#### E. Psychosocial Stress

- **Negative Life Events (PLE)**

○ **Source:** ABCD Life Events Scale (PLE) - Youth Report (Tiet et al., 2001).
○ **Scoring:** A **sum score** reflecting the total count of traumatic or stressful life events the child has *ever* experienced.
○ **Items Assessed:** The scale includes events such as natural disasters, serious accidents, witnessing violence, divorce/separation of parents, or death of a close family member.
○ **Interpretation:** Higher scores indicate a higher cumulative load of traumatic stress exposure.

### 5. Cognitive Measures (NIH Toolbox)

Cognitive performance was assessed using the NIH Toolbox Cognition Battery (NIHTB-CB), a computerized assessment suite administered on an iPad (Luciana et al., 2018; Weintraub et al., 2013). The battery yields seven individual subtest scores, which are aggregated into three composite measures of cognitive functioning: Fluid Cognition, Crystallized Cognition, and Total Cognition.

#### A. Fluid Cognition Domain (New Learning & Processing)

This domain measures the capacity for new learning, processing speed, and problem-solving in novel situations. It consists of five specific tasks:

1. **Flanker Inhibitory Control and Attention Test:** Measures attention and inhibitory control. Participants must indicate the orientation of a central arrow while ignoring flanking arrows that may be congruent or incongruent.
2. **Dimensional Change Card Sort (DCCS) Test:** Measures cognitive flexibility (set shifting). Participants match target cards by shape or color, switching rules frequently.
3. **Picture Sequence Memory Test:** Measures episodic memory. Participants must recall the sequence of pictured activities presented as a story.
4. **List Sorting Working Memory Test:** Measures working memory. Participants recall and sequence a series of visually and orally presented stimuli (animals and foods) by size.
5. **Pattern Comparison Processing Speed Test:** Measures processing speed. Participants rapidly identify whether two side-by-side images are identical or different.

#### B. Crystallized Cognition Domain (Accumulated Knowledge)

This domain measures accumulated knowledge and verbal ability, which are generally more dependent on education and cultural experience. It consists of two tasks:

1. **Picture Vocabulary Test:** Measures receptive vocabulary. Participants select which of four images best matches a spoken word.
2. **Oral Reading Recognition Test:** Measures reading decoding and language. Participants read and pronounce single words of increasing difficulty.

#### C. Composite Scores & Scoring

- **Scoring Method:** Raw scores from each subtest were converted into **Age-Corrected Standard Scores** (Mean = 100, SD = 15) based on the NIH Toolbox normative sample. This correction accounts for developmental improvements in cognition expected with age.
- ● **Fluid Cognition Composite:** The average of the age-corrected scores from the five fluid tasks (Flanker, DCCS, Picture Sequence, List Sorting, Pattern Comparison).
- ● **Crystallized Cognition Composite:** The average of the age-corrected scores from the two crystallized tasks (Picture Vocabulary, Oral Reading).
- ● **Total Cognition Composite:** A global summary score representing general intellectual functioning, derived by averaging the Fluid and Crystallized composite scores.

### Supplementary Materials: Methods

#### 2.1 Participants (ENIGMA-ASD)

Participants were drawn from the ENIGMA-ASD Working Group, a large-scale, multi-site effort, aggregating data from distinct international sites. See Supplemental Table S1 for full information regarding image acquisition parameters for each cohort. The van Rooij et al. (2018) study included a total of 1,571 individuals with autism and 1,651 healthy controls, with age ranges from 2–64 years old, drawn from 49 participating sites. Our updated dataset focused on a final analytic sample of 3,389 participants (1,655 autism and 1,734 neurotypical), restricted to ages 2–30 years. We analyzed data from 51 participating sites. Our current sample includes data from four independent cohorts not present in the Van Rooij et al. (2018) analysis: Barcelona, GENDAAR, SWEDEN, and DRESDEN (Total N of the new sites = 670). Additionally, our analysis includes the Nijmegen cohort (referenced as separate subsets: cattes, liehoe, and wougro), which was also present in the Van Rooij et al. (2018) study. Only the Toronto site (total N = 246) from the 2018 report was unavailable for this analysis. Ultimately, despite the stricter age exclusion criteria, the present study increases the sample size of individuals with autism from 1,571 to 1,655, providing a robust and expanded basis for direct comparison. The final analytic dataset included 3,389 individuals drawn from 51 independent sites.

#### 2.2 Cross-Cohort Reproducibility (COCORO-ASD)

Both studies utilized a meta-analytic approach for data harmonization. Specifically, effect sizes (unbiased Cohen’s d) were determined using a linear regression model calculated within each unique scanner and protocol group at each site, adjusting for age and sex as covariates. For subcortical volumes, intracranial volume (ICV) was included as an additional covariate. The individual site-level effect sizes were then combined using a random-effects model meta-analysis. We extracted these region-level effect sizes for cortical thickness (k=34 regions) and subcortical volumes (k=8 regions) from the published supplementary data (Matsumoto et al., 2023; Okada et al., 2023). Because the COCORO studies reported effects separately for left and right hemispheres, these values were averaged to match the bilateral region-of-interest (ROI) definitions used in the ENIGMA protocols (detailed in: http://enigma.ini.usc.edu/protocols/imaging-protocols/).The white matter RVI was constructed using regional effect sizes for fractional anisotropy (FA) reported by (Koshiyama et al., 2020). Because the COCORO studies reported effects separately for left and right hemispheres, these values were averaged to match the bilateral region-of-interest (ROI) definitions used in the ENIGMA protocols. We then performed Pearson correlation analyses to quantify the concordance of neuroanatomical profiles between the current updated ENIGMA dataset, the original ENIGMA findings, and the COCORO consortium results.

### Study II: Population-Based Investigation (ABCD Study)

#### 2.3 Participants (ABCD Study)

Participants for the ABCD study were also excluded at baseline if they presented with severe neurological, medical, or sensory impairments that would interfere with protocol compliance. Specific exclusionary diagnoses included schizophrenia, substance use disorders, and moderate-to-severe autism spectrum disorder or intellectual disability impacting the child’s ability to follow instructions. The final analytical sample was 54% male (n = 2,275) and 46% female (n = 1,926). Racial and ethnic composition included 60% White (n = 2,507), 16% Hispanic (n = 690), 14% Black (n = 591), 8.9% Other/Mixed (n = 372), and 1.0% Asian (n = 41). The ABCD study was designed to recruit from public schools to be representative of the U.S. child population. Written informed consent was provided by parents or guardians, and assent was provided by the children prior to participation. All data used in this study were acquired from the ABCD 5.0 data release. For the present study, data was limited to participants who had all data available (DTI and sMRI).

#### 2.4 Neuroimaging Data Acquisition and Preprocessing

Preprocessing of T1-weighted and diffusion tensor imaging (DTI) data was completed using the standardized Enhancing Neuro Imaging Genetics through Meta-Analysis (ENIGMA) consortium pipelines, which have been detailed previously (https://github.com/ENIGMA-git) using COCORO data The white matter RVI was constructed using regional effect sizes for fractional anisotropy (FA) reported by Koshiyama and colleagues (2020).

#### 2.5 Regional Vulnerability Index (RVI) Calculation

The ENIGMA consortium provided the meta-analytical ranks of the severity of differences associated with autism, and other disorders (e.g., SSD, BD, MDD, OCD) on gray matter thickness (33 cortical areas), subcortical volumes (7 structures) and fractional anisotropy (24 major white matter regions) (Boedhoe et al., 2020; Hibar et al., 2016, 2018, 2018; Kelly et al., 2018; Kochunov, Fan, et al., 2022; Piras et al., 2021; Schmaal et al., 2016, 2017; Van Erp et al., 2016; Van Velzen et al., 2020). The updated ENIGMA RVI-ASD results from this paper were provided from each working group and have not been added to the ENIGMA toolbox. For ABCD analysis the whole-brain RVI was constructed using updated ENIGMA-ASD cortical and subcortical and COCORO white matter effect sizes. Four dependent variables were selected as primary neuroanatomical outcomes: Gray Matter Thickness RVI, Gray Matter Volume RVI, whole-brain RVI-ASD, White Matter RVI-ASD.

These findings are provided as the regional effect sizes using Cohen’s *d* statistics after adjusting for age and sex and for intra-cranial volume for both subcortical and cortical (global correction was not used in the case of DTI-FA). RVI is a simple measure of agreement between an individual’s pattern of regional neuroimaging traits and the expected pattern of each specific disorder (such as autism, in this study) derived from ENIGMA meta-analyses in these traits. RVI is calculated as a single value per individual. The calculations for the unimodal RVIs are included below:

1. We calculated the residuals by regressing out effects of the covariates, including age, sex and intracranial brain volume using the sample in each scanning site, and then inverse-normalizing the residuals.
2. We performed a *z*-score transformation for each individual/tract measurement using the average and standard deviation values calculated in all subjects. This produced a phenotype vector of 24 z-scores for regional white matter measurement for every individual in the sample. The RVI is then calculated as a Pearson’s correlation coefficient between 24 region-wise *z* values for the subject and their corresponding effect sizes. This is repeated for gray matter thickness that is represented by a vector of 33 values and subcortical gray matter volume that is represented by a vector of 8 values.

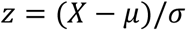

The multimodality whole-brain RVI-ASD was calculated following a rank normalization of ENIGMA’s Cohen’s *d* values for the cortical, subcortical, and white matter measures by sorting and assigning values between0.00to 1, i.e., the structure with the highest effect size had the absolute value of 1 and the structure with the lowest effect size had a value of 0. This normalization maintains the relative distance between individual effect sizes within each modality and is intended to prevent biases during the multimodality whole-brain RVI-ASD calculation due to potential differences in the magnitude of effect sizes across modalities. For example, it prevents the multimodality whole-brain RVI-ASD coefficient from being exaggerated by the difference in the direction and the magnitude of effect sizes in different modality clusters. The multimodal whole-brain RVI-ASD coefficient was calculated as the Pearson correlation between the full phenotypic vector of 64 values (33 cortical, 7 subcortical and 24 white matter) for each individual and the corresponding vector of normalized ENIGMA effect sizes.

#### 2.6 Genotyping (ABCD Study)

##### Empirical Kinship

The *pedifromsnps* function of the SOLAR-Eclipse software in a PLINK format (Gao et al., 2021), which uses weighted allelic correlation (WAC) (Hayes et al., 2009) (Eq. 1):

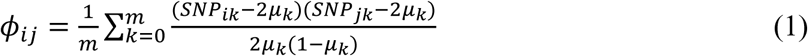

where *ϕij* is the empirical kinship matrix value between individual *i* and individual *j*. *m* is the total number of SNP loci for both individual *i* and individual *j*. *SNPik* and *SNPjk* are allelic scores (0, 1 or 2) for the *k*-th SNP in individuals *i* and *j*. *μk* is the frequency of the *k*-th major allele. Measuring the statistical power of pedigree using expected likelihood ratio test (ELRT). The ELRT method is used by SOLAR-Eclipse software to evaluate the statistical power of a pedigree for heritability analysis and to compare power between two pedigrees. This function is based on the functionality proposed by (Blijd-Hoogewys et al., 2014) and further generalized Raffa et al. (2016) (Raffa & Thompson, 2016). The ELRT is defined as the expectation of twice the difference of the log-likelihoods evaluated at the true parameter and at several different null-parameter values, respectively (Raffa & Thompson, 2016). It uses Taylor series approximations to summarize the relatedness in a pedigree to accurately approximate the expectation of the likelihood ratio test and expected confidence interval widths (Raffa & Thompson, 2016).

##### Heritability, genetic correlation and genotype-by-age interaction

We performed polygenic modeling of the source of individual variance to study the factors that contributed to the baseline and longitudinal changes in RVI for autism. Polygenic model parcellates the total variance in the trait into additive genetic (G), environmental (E) and genetic interactions, such as genetic by age (GxA). Genetic correlation analysis can further be used to interrogate the G and E components between two traits by calculating the contribution from suspected factors while adjusting for genetic variance shared among subjects (G). The longitudinal data allows for calculation of GxA interaction, as well as to screen potential environmental factors as contributors via E. For example, an increase in trait’s heritability (*h^2^*) between two longitudinal measurements indicates an increase in genetic control over variance. This can occur due to GxA interaction whereby additive genetic factors start to exert higher control, or because of reduction in E, which can be further screened using genetic correlation analyses to screen known environmental factors. This makes ABCD a powerful sample for discovery of genetic and environmental factors, where longitudinal data collection allows for evaluation of the nature vs nurture effects on development of different brain patterns (Blangero, 2009). Classically, these analyses were limited to twin- or -family study design (Blangero et al., 2013; Chouinard-Decorte et al., 2014; Glahn et al., 2013). Here, we used novel methods implemented in SOLAR-Eclipse to perform these analyses in the full ABCD data and compared traditional *h^2^* estimates derived from self-reported relationship vs. empirical derived from quantitative analyses of genetic data and observed a good agreement (Smith et al., 2023).

##### Standard polygenic model

A common analysis approach when analyzing data from related subjects use linear mixed models to account for genetic relatedness (non-independence) among the subjects (Eq. 1). For simplicity of the equations, we make two assumptions: effects of fixed covariates such as age, sex, scanner, site, etc., were removed at the point of preprocessing of the phenotypes and the data were inverse-normalized such that each trait follows a multivariate normal distribution. In this general case, the linear mixed effect model for effects of a measured genotype (SNP) in a sample of non-independent subjects is shown in Eq. 2:

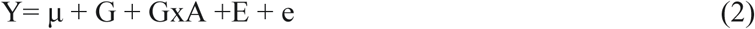

Two extensions of standard polygenic models using in this study are: bivariate genetic correlation and genotype by age interaction.

##### Bivariate genetic correlation

The overall phenotypic correlation (ρP) between two traits A and B (Eq 3) can be expressed using the correlation due to shared additive genetic effects (ρG) and the residual correlation (ρE) due to shared environmental effects.

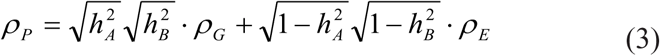

Where, h^2^A and h^2^B denote the heritability for each of the traits i.e., the proportion of the total phenotypic variance that is explained by additive genetic factors. If the genetic correlation coefficient (rG) is significantly different from zero, then the significant portion of the variance in two traits are considered to be influenced by shared genetic factors (Almasy et al., 1997; Kochunov, Glahn, Lancaster, et al., 2011; Kochunov, Glahn, Nichols, et al., 2011).

##### Genotype by age interaction

The degree of additive genetic control (h^2^) over trait can change as a part of normal development. This can happen due to change in proportion of variance due to environmental variance or biologically pre-programmed through genotype-by-age interaction. We define genotype by age interaction as a significant additive component of variance in response to the development (Blangero, 2009; Chouinard-Decorte et al., 2014; Glahn et al., 2013). This additive genetic variance in response (*σ*^2^*GΔ*) is a function of the additive genetic variance of the trait expressed during development e.g., baseline and follow up (Eq. 4):

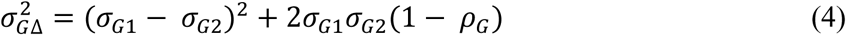

The null hypothesis is that the baseline and follow up trait have full pleiotropy (ρ*G* = 1) of no change in additive genetic variance between the two environments. In the context of G × A interactions, the requirement of full pleiotropy means that the same set of genes must account for the observed additive variance of the trait across age. We can test it by computing two genetic correlation models. The null model is calculated under the constraint that baseline heritability for the trait is same as the follow up heritability. The hypothesis model is the genetic correlation between base and follow up trait values. Once both models have been maximized and their corresponding log-likelihoods calculated, the p-value is computed first by taking the difference between the hypothesis log-likelihood and the null log-likelihood. Next, we multiply this value by negative two and enter it into the chi-squared cumulative distribution function, with one degree of freedom. Finally, the result is divided by two, because we are dealing with a single tailed p-value.

### Supplemental Information: ENIGMA-ASD Working Group Members and Affiliations

- **Celso Arango, MD, PhD** Child and Adolescent Psychiatry Department, Gregorio Marañón General University Hospital, School of Medicine, Universidad Complutense, IiSGM, CIBERSAM, Madrid, Spain
- **Guillaume Auzias, PhD** Institut de Neurosciences de la Timone, UMR 7289, Aix Marseille Université, CNRS, Marseille, France
- **Marlene Behrmann, PhD** Department of Psychology, Carnegie Mellon University, Pittsburgh, PA, USA
- **Sven Bolte, PhD** Center of Neurodevelopmental Disorders (KIND), Centre for Psychiatry Research; Department of Women’s and Children’s Health, Karolinska Institutet & Stockholm Health Care Services, Region Stockholm, Stockholm, Sweden; Child and Adolescent Psychiatry, Stockholm Health Care Services, Region Stockholm, Stockholm, Sweden; Curtin Autism Research Group, Curtin School of Allied Health, Curtin University, Perth, Western Australia
- **Paolo Bosco, PhD** National Institute for Nuclear Physics, Pisa Division, Largo B. Pontecorvo 3, 56124, Pisa, Italy
- **Jan Buitelaar, MD, PhD** Department of Cognitive Neuroscience, Donders Institute for Brain, Cognition and Behaviour, Radboud University Medical Centre, Nijmegen, The Netherlands
- **Geraldo Busatto, PhD** Department of Psychiatry, Faculty of Medicine, University of São Paulo, São Paulo, Brazil; Center for Interdisciplinary Research on Applied Neurosciences (NAPNA), University of São Paulo, São Paulo, Brazil
- **Sara Calderoni, MD, PhD** IRCCS Stella Maris Foundation, viale del Tirreno 331, 56128, Pisa, Italy
- **Rosa Calvo, MD, PhD** Department of Child and Adolescent Psychiatry and Psychology Hospital Clinic, Barcelona CIBERSAM, Universitat de Barcelona
- **Eileen Daly, PhD** Department of Forensic and Neurodevelopmental Sciences, Institute of Psychiatry, Psychology & Neuroscience, King’s College London, London, UK
- **Christine Deruelle, PhD** Institut de Neurosciences de la Timone, UMR 7289, Aix Marseille Université, CNRS, Marseille, France
- **Ilan Dinstein, PhD** Department of Psychology, Ben-Gurion University of the Negev, Beer Sheva, Israel
- **Fabio Duran, MD** Departamento de Neuropsiquiatria, UFSM - Universidade Federal de Santa Maria
- **Sarah Durston, PhD** Brain Center Rudolf Magnus, Department of Psychiatry, University Medical Center Utrecht, The Netherlands
- **Christine Ecker, PhD** Department of Child and Adolescent Psychiatry, Psychosomatics and Psychotherapy, University Hospital, Goethe University Frankfurt am Main, Frankfurt, Germany
- **Stephan Ehrlich, MD, PhD** Division of Psychological and Social Medicine and Developmental Neurosciences, Faculty of Medicine, TU Dresden, Germany
- **Damien Fair, PhD** Institute of Child Development, University of Minnesota, Minneapolis, MN, USA; Masonic Institute for the Developing Brain, University of Minnesota, Minneapolis, MN, USA; Department of Pediatrics, University of Minnesota, Minneapolis, MN, USA
- **Jennifer Fedor, BS** Department of Psychiatry, University of Pittsburgh, Pittsburgh, PA, USA
- **Jackie Fitzgerald, PhD** Department of Psychiatry, School of Medicine, Trinity College, Dublin, Ireland
- **Christine M. Freitag, PhD** Department of Child and Adolescent Psychiatry, Psychosomatics and Psychotherapy, University Hospital, Goethe University Frankfurt am Main, Frankfurt, Germany
- **Louise Gallagher, MD, PhD** Department of Psychiatry, School of Medicine, Trinity College, Dublin, Ireland
- **Ilaria Gori, MSc** National Institute for Nuclear Physics, Pisa Division, Largo B. Pontecorvo 3, 56124, Pisa, Italy
- **Shlomi Haar, PhD** Department of Brain and Cognitive Sciences, Zlotowski Center for Neuroscience, Ben-Gurion University of the Negev, Beer Sheva, Israel
- **Liesbeth Hoekstra, MSc** Department of Cognitive Neuroscience, Donders Institute for Brain, Cognition and Behaviour, Donders Centre for Cognitive Neuroimaging, Radboud University Medical Centre, Nijmegen, The Netherlands
- **Zachary Jacokes** School of Data Science, University of Virginia, Charlottesville, VA, USA
- **Maria Jalbrzikowski, PhD** Department of Psychiatry, School of Medicine, Trinity College, Dublin, Ireland
- **Joost Janssen, PhD** Child and Adolescent Psychiatry Department, Gregorio Marañón General University Hospital, School of Medicine, Universidad Complutense, IiSGM, CIBERSAM, Madrid, Spain
- **Gaon S. Kim, MS** Imaging Genetics Center, Mark and Mary Stevens Neuroimaging and Informatics Institute, Keck School of Medicine, University of Southern California, Marina Del Rey, CA, USA
- **Joseph King, PhD** Division of Psychological and Social Medicine and Developmental Neurosciences, Faculty of Medicine, TU Dresden, Germany
- **Katherine E. Lawrence, PhD** Imaging Genetics Center, Mark and Mary Stevens Neuroimaging and Informatics Institute, Keck School of Medicine, University of Southern California, Marina Del Rey, CA, USA
- **Noemi Gonzalez Lois** Child and Adolescent Psychiatry Department, Gregorio Marañón General University Hospital, School of Medicine, Universidad Complutense, IiSGM, CIBERSAM, Madrid, Spain
- **Beatriz Luna, PhD** Department of Psychiatry, University of Pittsburgh, Pittsburgh, PA, USA
- **Luisa Lázaro, MD, PhD** Department of Child and Adolescent Psychiatry and Psychology Hospital Clinic, Barcelona CIBERSAM, IDIBAPS
- **Mauricio Martinho, MD** Department of Psychiatry, Faculty of Medicine, University of São Paulo, São Paulo, Brazil; Center for Interdisciplinary Research on Applied Neurosciences (NAPNA), University of São Paulo, São Paulo, Brazil
- **Jane McGrath, PhD** Department of Psychiatry, School of Medicine, Trinity College, Dublin, Ireland
- **Filippo Muratori, PhD** IRCCS Stella Maris Foundation, viale del Tirreno 331, 56128, Pisa, Italy
- **Declan G.M. Murphy, MD, FRCPsyc** The Sackler Institute for Translational Neurodevelopment, Institute of Psychiatry, Psychology & Neuroscience, King’s College London, London, UK
- **Janina Neufeld, PhD** Center of Neurodevelopmental Disorders (KIND), Centre for Psychiatry Research; Department of Women’s and Children’s Health, Karolinska Institutet & Stockholm Health Care Services, Region Stockholm, Stockholm, Sweden
- **Benjamin Franklin McGartland Newman, PhD** Department of Psychology, University of Virginia, Charlottesville, VA, USA; VA School of Medicine, University of Virginia, Charlottesville, VA, USA
- **Bob Oranje, PhD** Brain Center Rudolf Magnus, Department of Psychiatry, University Medical Center Utrecht, The Netherlands
- **Kirsten O’Hearn, PhD** Department of Physiology and Pharmacology, Wake Forest School of Medicine, Winston-Salem, NC, USA
- **Mara Parellada, MD, PhD** Child and Adolescent Psychiatry Department, Gregorio Marañón General University Hospital, School of Medicine, Universidad Complutense, IiSGM, CIBERSAM, Madrid, Spain
- **Jose Pariente** Department of Child and Adolescent Psychiatry and Psychology Hospital Clinic, Barcelona CIBERSAM
- **Karl Lundin Remnélius, PhD** Center of Neurodevelopmental Disorders (KIND), Centre for Psychiatry Research; Department of Women’s and Children’s Health, Karolinska Institutet & Stockholm Health Care Services, Region Stockholm, Stockholm, Sweden
- **Alessandra Retico, PhD** National Institute for Nuclear Physics, Pisa Division, Largo B. Pontecorvo 3, 56124, Pisa, Italy
- **Pedro Rosa, MD** Department of Psychiatry, Faculty of Medicine, University of São Paulo, São Paulo, Brazil; Center for Interdisciplinary Research on Applied Neurosciences (NAPNA), University of São Paulo, São Paulo, Brazil
- **Katya Rubia, PhD** Institute of Psychiatry, Psychology and Neuroscience, Kings College London, London, UK
- **Devon Shook, PhD** Brain Center Rudolf Magnus, Department of Psychiatry, University Medical Center Utrecht, The Netherlands
- **Kristiina Tammimies, PhD** Center of Neurodevelopmental Disorders (KIND), Centre for Psychiatry Research; Department of Women’s and Children’s Health, Karolinska Institutet & Stockholm Health Care Services, Region Stockholm, Stockholm, Sweden
- **Michela Tosetti, PhD** IRCCS Stella Maris Foundation, viale del Tirreno 331, 56128, Pisa, Italy
- **John D. Van Horn, PhD** School of Data Science, University of Virginia, Charlottesville, VA, USA; Department of Psychology, University of Virginia, Charlottesville, VA, USA
- **Daan van Rooij, PhD** Experimental Psychology, Helmholtz Institute, Utrecht University, Netherlands
- **Gregory L. Wallace, PhD** Department of Speech and Hearing Sciences, The George Washington University, Washington, DC, USA

## Notes

### Competing Interest Statement

The authors have declared no competing interest.

## References

1. American Psychiatric Association. Diagnostic and Statistical Manual of Mental Disorders. DSM-5-TR. American Psychiatric Association Publishing; 2022.

2. Maenner MJ, Warren Z, Williams AR, Amoakohene E, Bakian AV, Bilder DA, et al. Prevalence and Characteristics of Autism Spectrum Disorder Among Children Aged 8 Years — Autism and Developmental Disabilities Monitoring Network, 11 Sites, United States, 2020. MMWR Surveill Summ. 2023;72:1–14.

3. Karst JS, Van Hecke AV. Parent and Family Impact of Autism Spectrum Disorders: A Review and Proposed Model for Intervention Evaluation. Clin Child Fam Psychol Rev. 2012;15:247–277.

4. Duan X, Shan X, Uddin LQ, Chen H. The Future of Disentangling the Heterogeneity of Autism With Neuroimaging Studies. Biological Psychiatry. 2025;97:428–438.

5. Lenroot RK, Yeung PK. Heterogeneity within Autism Spectrum Disorders: What have We Learned from Neuroimaging Studies? Front Hum Neurosci. 2013;7.

6. Lord C, Elsabbagh M, Baird G, Veenstra-Vanderweele J. Autism spectrum disorder. The Lancet. 2018;392:508–520.

7. Constantino JN, Todd RD. Autistic Traits in the General Population: A Twin Study. Arch Gen Psychiatry. 2003;60:524.

8. Robinson EB, St Pourcain B, Anttila V, Kosmicki JA, Bulik-Sullivan B, Grove J, et al. Genetic risk for autism spectrum disorders and neuropsychiatric variation in the general population. Nat Genet. 2016;48:552–555.

9. Cruz Puerto M, Sandín Vázquez M. Understanding heterogeneity within autism spectrum disorder: a scoping review. AIA. 2024;10:314–322.

10. Masi A, DeMayo MM, Glozier N, Guastella AJ. An Overview of Autism Spectrum Disorder, Heterogeneity and Treatment Options. Neurosci Bull. 2017;33:183–193.

11. Happé F, Frith U. Annual Research Review: Looking back to look forward – changes in the concept of autism and implications for future research. Child Psychology Psychiatry. 2020;61:218–232.

12. Sucksmith E, Roth I, Hoekstra RA. Autistic Traits Below the Clinical Threshold: Re-examining the Broader Autism Phenotype in the 21st Century. Neuropsychol Rev. 2011;21:360–389.

13. Warrier V, Zhang X, Reed P, Havdahl A, Moore TM, Cliquet F, et al. Genetic correlates of phenotypic heterogeneity in autism. Nat Genet. 2022;54:1293–1304.

14. Abu-Akel A, Allison C, Baron-Cohen S, Heinke D. The distribution of autistic traits across the autism spectrum: evidence for discontinuous dimensional subpopulations underlying the autism continuum. Molecular Autism. 2019;10:24.

15. Bethlehem RAI, Seidlitz J, White SR, Vogel JW, Anderson KM, Adamson C, et al. Brain charts for the human lifespan. Nature. 2022;604:525–533.

16. Waterhouse L. Heterogeneity thwarts autism explanatory power: A proposal for endophenotypes. Front Psychiatry. 2022;13:947653.

17. Kochunov P, Hong LE, Dennis EL, Morey RA, Tate DF, Wilde EA, et al. ENIGMA-DTI: Translating reproducible white matter deficits into personalized vulnerability metrics in cross-diagnostic psychiatric research. Human Brain Mapping. 2022;43:194–206.

18. Koshiyama D, Fukunaga M, Okada N, Morita K, Nemoto K, Usui K, et al. White matter microstructural alterations across four major psychiatric disorders: mega-analysis study in 2937 individuals. Mol Psychiatry. 2020;25:883–895.

19. Van Rooij D, Anagnostou E, Arango C, Auzias G, Behrmann M, Busatto GF, et al. Cortical and Subcortical Brain Morphometry Differences Between Patients With Autism Spectrum Disorder and Healthy Individuals Across the Lifespan: Results From the ENIGMA ASD Working Group. AJP. 2018;175:359–369.

20. Boedhoe PSW, Van Rooij D, Hoogman M, Twisk JWR, Schmaal L, Abe Y, et al. Subcortical Brain Volume, Regional Cortical Thickness, and Cortical Surface Area Across Disorders: Findings From the ENIGMA ADHD, ASD, and OCD Working Groups. AJP. 2020;177:834–843.

21. Thompson PM, Jahanshad N, Schmaal L, Turner JA, Winkler AM, Thomopoulos SI, et al. The Enhancing NEUROIMAGING Genetics through Meta-Analysis Consortium: 10 Years of Global Collaborations in Human Brain Mapping. Human Brain Mapping. 2022;43:15–22.

22. Matsumoto J, Fukunaga M, Miura K, Nemoto K, Okada N, Hashimoto N, et al. Cerebral cortical structural alteration patterns across four major psychiatric disorders in 5549 individuals. Mol Psychiatry. 2023;28:4915–4923.

23. Okada N, Fukunaga M, Miura K, Nemoto K, Matsumoto J, Hashimoto N, et al. Subcortical volumetric alterations in four major psychiatric disorders: a mega-analysis study of 5604 subjects and a volumetric data-driven approach for classification. Mol Psychiatry. 2023;28:5206–5216.

24. Ramaswami G, Geschwind DH. Genetics of autism spectrum disorder. Handbook of Clinical Neurology, vol. 147, Elsevier; 2018. p. 321–329.

25. Woodbury-Smith M, Scherer SW. Progress in the genetics of autism spectrum disorder. Develop Med Child Neuro. 2018;60:445–451.

26. Sandin S, Lichtenstein P, Kuja-Halkola R, Larsson H, Hultman CM, Reichenberg A. The Familial Risk of Autism. JAMA. 2014;311:1770.

27. Modabbernia A, Velthorst E, Reichenberg A. Environmental risk factors for autism: an evidence-based review of systematic reviews and meta-analyses. Molecular Autism. 2017;8:13.

28. Shelton JF, Tancredi DJ, Hertz-Picciotto I. Independent and dependent contributions of advanced maternal and paternal ages to autism risk. Autism Research. 2010;3:30–39.

29. Wu S, Wu F, Ding Y, Hou J, Bi J, Zhang Z. Advanced parental age and autism risk in children: a systematic review and meta-analysis. Acta Psychiatr Scand. 2017;135:29–41.

30. Janecka M, Mill J, Basson MA, Goriely A, Spiers H, Reichenberg A, et al. Advanced paternal age effects in neurodevelopmental disorders—review of potential underlying mechanisms. Transl Psychiatry. 2017;7:e1019–e1019.

31. Reichenberg A, Gross R, Weiser M, Bresnahan M, Silverman J, Harlap S, et al. Advancing Paternal Age and Autism. Arch Gen Psychiatry. 2006;63:1026.

32. Sandin S, Schendel D, Magnusson P, Hultman C, Surén P, Susser E, et al. Autism risk associated with parental age and with increasing difference in age between the parents. Mol Psychiatry. 2016;21:693–700.

33. Taylor JL, Debost J-CPG, Morton SU, Wigdor EM, Heyne HO, Lal D, et al. Paternal-age-related de novo mutations and risk for five disorders. Nat Commun. 2019;10:3043.

34. Wang C, Geng H, Liu W, Zhang G. Prenatal, perinatal, and postnatal factors associated with autism: A meta-analysis. Medicine. 2017;96:e6696.

35. Lord C, Risi S, Lambrecht L, Cook, Jr. EH, Leventhal BL, DiLavore PC, et al. The Autism Diagnostic Observation Schedule—Generic. Journal of Autism and Developmental Disorders. 2000;30:205–223.

36. Constantino JN. Social Responsiveness Scale. In: Volkmar FR, editor. Encyclopedia of Autism Spectrum Disorders, New York, NY: Springer New York; 2013. p. 2919–2929.

37. Barch DM, Albaugh MD, Baskin-Sommers A, Bryant BE, Clark DB, Dick AS, et al. Demographic and mental health assessments in the adolescent brain and cognitive development study: Updates and age-related trajectories. Developmental Cognitive Neuroscience. 2021;52:101031.

38. Casey BJ, Cannonier T, Conley MI, Cohen AO, Barch DM, Heitzeg MM, et al. The Adolescent Brain Cognitive Development (ABCD) study: Imaging acquisition across 21 sites. Developmental Cognitive Neuroscience. 2018;32:43–54.

39. Hagler DJ, Hatton SeanN, Cornejo MD, Makowski C, Fair DA, Dick AS, et al. Image processing and analysis methods for the Adolescent Brain Cognitive Development Study. NeuroImage. 2019;202:116091.

40. Volkow ND, Koob GF, Croyle RT, Bianchi DW, Gordon JA, Koroshetz WJ, et al. The conception of the ABCD study: From substance use to a broad NIH collaboration. Developmental Cognitive Neuroscience. 2018;32:4–7.

41. Fan CC, Loughnan R, Wilson S, Hewitt JK, ABCD Genetic Working Group, Agrawal A, et al. Genotype Data and Derived Genetic Instruments of Adolescent Brain Cognitive Development Study® for Better Understanding of Human Brain Development. Behav Genet. 2023;53:159–168.

42. Hibar DP, Westlye LT, Van Erp TGM, Rasmussen J, Leonardo CD, Faskowitz J, et al. Subcortical volumetric abnormalities in bipolar disorder. Mol Psychiatry. 2016;21:1710–1716.

43. Hibar DP, Westlye LT, Doan NT, Jahanshad N, Cheung JW, Ching CRK, et al. Cortical abnormalities in bipolar disorder: an MRI analysis of 6503 individuals from the ENIGMA Bipolar Disorder Working Group. Mol Psychiatry. 2018;23:932–942.

44. Kelly S, Jahanshad N, Zalesky A, Kochunov P, Agartz I, Alloza C, et al. Widespread white matter microstructural differences in schizophrenia across 4322 individuals: results from the ENIGMA Schizophrenia DTI Working Group. Mol Psychiatry. 2018;23:1261–1269.

45. Kochunov P, Fan F, Ryan MC, Hatch KS, Tan S, Jahanshad N, et al. Translating ENIGMA schizophrenia findings using the regional vulnerability index: Association with cognition, symptoms, and disease trajectory. Human Brain Mapping. 2022;43:566–575.

46. Piras F, Piras F, Abe Y, Agarwal SM, Anticevic A, Ameis S, et al. White matter microstructure and its relation to clinical features of obsessive–compulsive disorder: findings from the ENIGMA OCD Working Group. Transl Psychiatry. 2021;11:173.

47. Schmaal L, Veltman DJ, Van Erp TGM, Sämann PG, Frodl T, Jahanshad N, et al. Subcortical brain alterations in major depressive disorder: findings from the ENIGMA Major Depressive Disorder working group. Mol Psychiatry. 2016;21:806–812.

48. Schmaal L, Hibar DP, Sämann PG, Hall GB, Baune BT, Jahanshad N, et al. Cortical abnormalities in adults and adolescents with major depression based on brain scans from 20 cohorts worldwide in the ENIGMA Major Depressive Disorder Working Group. Mol Psychiatry. 2017;22:900–909.

49. Van Erp TGM, Hibar DP, Rasmussen JM, Glahn DC, Pearlson GD, Andreassen OA, et al. Subcortical brain volume abnormalities in 2028 individuals with schizophrenia and 2540 healthy controls via the ENIGMA consortium. Mol Psychiatry. 2016;21:547–553.

50. Van Velzen LS, Kelly S, Isaev D, Aleman A, Aftanas LI, Bauer J, et al. White matter disturbances in major depressive disorder: a coordinated analysis across 20 international cohorts in the ENIGMA MDD working group. Mol Psychiatry. 2020;25:1511–1525.

51. Gao S, Kochunov P, Hatch K, Ma Y, Talib F. Package ‘RVIpkg’. 2025. 2025.

52. Team P. RStudio: integrated development environment for R. Posit Software, PBC, Boston, MA. 2023.

53. Team RC. R: A language and environment for statistical computing. R Foundation for Statistical Computing, Vienna, Austria. Http://Www.R-Project.Org/. 2016. 2016.

54. Fan CC, Loughnan R, Wilson S, Hewitt JK, ABCD Genetic Working Group, Agrawal A, et al. Genotype Data and Derived Genetic Instruments of Adolescent Brain Cognitive Development Study® for Better Understanding of Human Brain Development. Behav Genet. 2023;53:159–168.

55. Weintraub S, Dikmen SS, Heaton RK, Tulsky DS, Zelazo PD, Bauer PJ, et al. Cognition assessment using the NIH Toolbox. Neurology. 2013;80.

56. Bartko JJ. The Intraclass Correlation Coefficient as a Measure of Reliability. Psychol Rep. 1966;19:3–11.

57. Coupé P, Catheline G, Lanuza E, Manjón JV, for the Alzheimer’s Disease Neuroimaging Initiative. Towards a unified analysis of brain maturation and aging across the entire lifespan: A MRI analysis. Human Brain Mapping. 2017;38:5501–5518.

58. Kochunov P, Ryan MC, Yang Q, Hatch KS, Zhu A, Thomopoulos SI, et al. Comparison of regional brain deficit patterns in common psychiatric and neurological disorders as revealed by big data. NeuroImage: Clinical. 2021;29:102574.

59. Kochunov P, Gao S, Salminen L, Jahanshad N, Nir T, Thompson P, et al. Alzheimer’s Disease-Like Brain Pattern Biomarker: Capturing Risks and Predicting Disease Onset. 2025.

60. Kochunov P, Hong LE, Dennis EL, Morey RA, Tate DF, Wilde EA, et al. ENIGMA-DTI: Translating reproducible white matter deficits into personalized vulnerability metrics in cross-diagnostic psychiatric research. Human Brain Mapping. 2022;43:194–206.

61. Kochunov P, Jahanshad N, Marcus D, Winkler A, Sprooten E, Nichols TE, et al. Heritability of fractional anisotropy in human white matter: A comparison of Human Connectome Project and ENIGMA-DTI data. NeuroImage. 2015;111:300–311.

62. Lenroot RK, Schmitt JE, Ordaz SJ, Wallace GL, Neale MC, Lerch JP, et al. Differences in genetic and environmental influences on the human cerebral cortex associated with development during childhood and adolescence. Human Brain Mapping. 2009;30:163–174.

63. Elvevag B, Goldberg TE. Cognitive Impairment in Schizophrenia Is the Core of the Disorder. Crit Rev Neurobiol. 2000;14:21.

64. Kochunov P, Ganjgahi H, Winkler A, Kelly S, Shukla DK, Du X, et al. Heterochronicity of white matter development and aging explains regional patient control differences in schizophrenia. Human Brain Mapping. 2016;37:4673–4688.

65. Kochunov P, Coyle TR, Rowland LM, Jahanshad N, Thompson PM, Kelly S, et al. Association of White Matter With Core Cognitive Deficits in Patients With Schizophrenia. JAMA Psychiatry. 2017;74:958.

66. Kochunov P, Ma Y, Hatch KS, Gao S, Jahanshad N, Thompson PM, et al. Brain-wide versus genome-wide vulnerability biomarkers for severe mental illnesses. Human Brain Mapping. 2022;43:4970–4983.

67. Charman T, Jones CRG, Pickles A, Simonoff E, Baird G, Happé F. Defining the cognitive phenotype of autism. Brain Research. 2011;1380:10–21.

68. Tillmann J, San José Cáceres A, Chatham CH, Crawley D, Holt R, Oakley B, et al. Investigating the factors underlying adaptive functioning in autism in the EU-AIMS Longitudinal European Autism Project. Autism Research. 2019;12:645–657.

69. Owen MJ, O’Donovan MC. Schizophrenia and the neurodevelopmental continuum:evidence from genomics. World Psychiatry. 2017;16:227–235.

70. Rapoport JL, Giedd JN, Gogtay N. Neurodevelopmental model of schizophrenia: update 2012. Mol Psychiatry. 2012;17:1228–1238.

71. Ecker C, Bookheimer SY, Murphy DGM. Neuroimaging in autism spectrum disorder: brain structure and function across the lifespan. The Lancet Neurology. 2015;14:1121–1134.

